# A lack of distinct cell identities in single-cell measurements: revisiting Waddington’s landscape

**DOI:** 10.1101/2022.06.03.494765

**Authors:** Breanne Sparta, Timothy Hamilton, Juan V. Najar, Serena Hughes, Gunalan Natesan, Eric J. Deeds

## Abstract

The prevailing interpretation of Waddington’s landscape is that attractors in gene expression space produce and stabilize distinct cell types. This notion is often applied in single-cell omics, where transcriptome data is clustered to produce models of cell types. Here we apply graph theory to characterize the distribution of cells in epigenetic space, using data from various tissues, organisms, and single-cell omics technologies. We find that cells of different types exist in the same regions of epigenetic space, with highly heterogeneous density distributions that are inconsistent with expected densities near an attractor. The lack of attractor structure could not be explained by technical noise, scale variance among genes, nor the subset of genes that were used; nor could it be rescued by any standard set of transformations. These findings pose a challenge for the robust analysis of single-cell data and open the possibility for alternative explanations of canalization during development.

## Introduction

Many multicellular taxa exhibit extreme cellular differentiation, where, despite possessing the same genetic material, different cells within an individual perform highly divergent and specialized functions. For instance, human neutrophils are amoeboid and excel at pursuing and engulfing pathogens, while neurons are non-motile and adopt elaborate dendritic and axonic structures. The question of how difference is generated *within* an individual organism has confronted the biological imagination for the last 150 years^1–6^. For instance, in the late 1800s August Weismann posited that stable differentiation was achieved through irreversible conversion of cellular components from the “germ plasm” into somatic structures^7^. Weissman’s view, while compelling, was of course incorrect, as all cells in most multicellular organisms contain more-or-less the same genome.

Our current understanding for the molecular basis of development rests on ideas from dynamical systems theory that are schematized in Waddington’s “epigenetic landscape”^8–10^. In this picture, cell differentiation is depicted as movement through a landscape of epigenetic constraints, descending through valleys that guide the production of terminally differentiated cell fates, which correspond to attractors in the landscape (Fig. 1A)^8–10^. In its original conception, Waddington’s notion of the state variables in these epigenetic landscapes was abstract. Yet, in the 1950’s, the emergence of the central dogma of molecular biology narrowed the conceptual framework. The landscape shifted from an abstract representation of cell state to one where the state variables of cell-fate attractors could be quantitatively represented by mRNA and protein concentrations^11–17^. This leap was prompted in part by the growing popularity of Jacob and Monod’s wiring diagrams for differential gene regulation, which provided a clear mechanistic hypothesis for epigenesis^13,18–21^. In parallel, developments in nonlinear dynamical systems theory demonstrated that systems with complex interactions could have multiple stable point attractors, each with their own corresponding basins of attraction (Fig. 1B)^12,22–25^.

**Figure 1.**
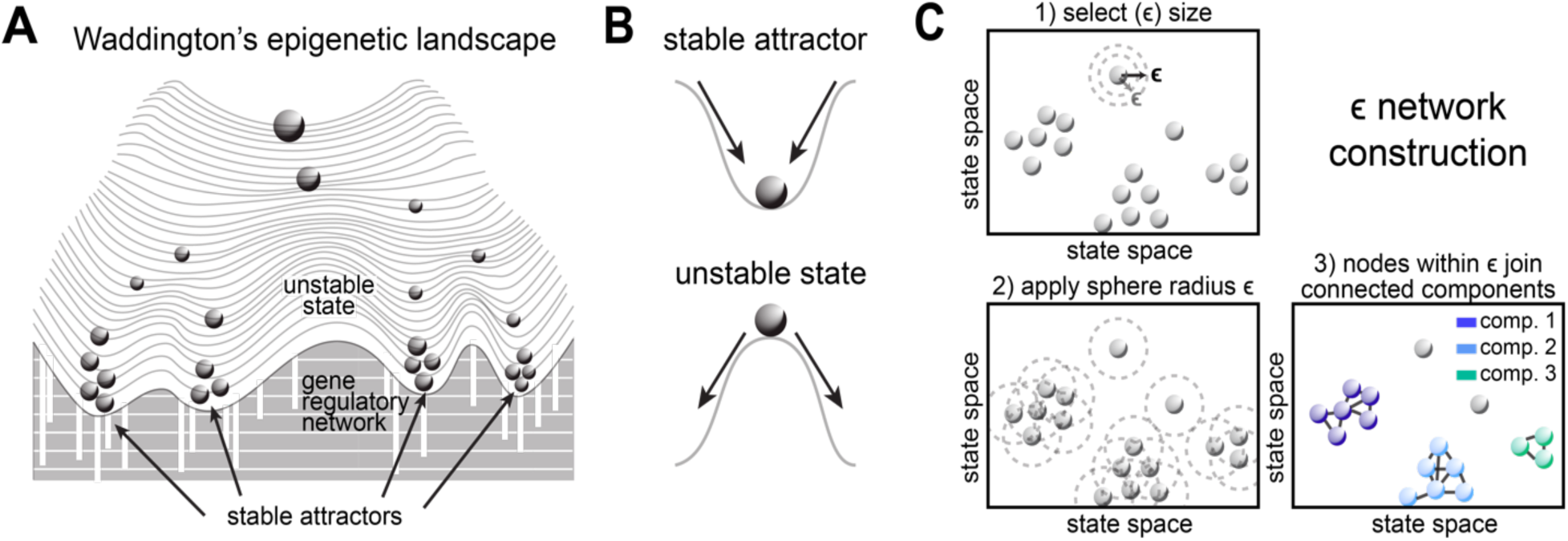
Waddington’s Epigenetic Landscape draws predictions from dynamical systems theory. **A.** Schematic of Waddington’s landscape, where the epigenetic state of developing cells is *canalized* by interactions in gene regulatory networks. Cell types are thought to represent different “basins of attraction” in this landscape, and cells of the same type should have similar gene expression patterns. **B.** Schematic of the states that construct the hills and valleys of Waddington’s Landscape. The valleys represent stable attractors, and function to buffer a cell from perturbations. The hills correspond to unstable states, where any perturbation results in increasing divergence from the original state. **C.** Schematic of *∈* network construction. First, we consider a hypersphere of radius *∈* around any given cell in the dataset (panel 1). We consider a similar hypersphere for every cell in the dataset (panel 2). For each cell, an edge is drawn between the cell and any neighbor that is within the *∈* radius. This generates a network that connects cells (panel 3). Note that, at any given *∈*, the network naturally forms a set of clusters of cells that are all connected to each other, called “components” in graph theory. The largest of these is the giant component.

The contemporary interpretation of Waddington’s landscape is clear: different cell types correspond to different attractors in a landscape where the state variables are either mRNA levels or protein levels, depending on the specific context^11,13,26^. A large body of mathematical and experimental work has characterized how dynamical systems models of gene regulatory networks can generate these attractors, and how bifurcations can arise during development to allow cells to proceed down different developmental paths^12,27–29^. This underlying picture for differentiation during development is compelling, as it provides a generative model for cell types as well as a molecular basis for “canalization” in development. Canalization ensures the developmental process is robust, such that cell lineages and terminally differentiated cell types do not switch fates despite internal gene expression noise or small environmental perturbations. One of the key predictions of this modern interpretation of Waddington’s landscape is that phenotypically similar cells should have similar molecular compositions^27,30–32^.

During the past few decades, new measurement technologies have enabled characterization of the molecular state of individual cells, with increasing precision and sampling capacities. In particular, single-cell RNA-sequencing (scRNA-seq) provides transcriptome-wide information about mRNA levels across hundreds of thousands of individual cells^33^. This data provides a unique opportunity to characterize attractors in the gene expression landscape and to test our conceptual frameworks of cell differentiation and biological organization. Yet, the majority of studies employing scRNA-seq technology take *a priori* the assumption that clustering in gene expression space can produce models of cell types, upon which analyses like differential cell-type composition or differential gene expression can be performed. In practice, however, the analysis of this data has revealed incredible heterogeneity in cell state, regardless of the particular measurement technology employed^34–36^. Moreover, analytical pipelines that attempt to find cell type clusters (Fig. 1C) involve many nonlinear transformations and dimensionality reduction steps, and the clusters obtained are often very sensitive to the parameters (Fig. S1)^37–41^. For example, we found that even arbitrary parameter choices such as removing the smallest cluster from a dataset, or even just removing 1% of cells at random, dramatically changes the clustering membership results (Fig. S1). This practical difficulty in identifying cell-type clusters raises the question of whether developmentally robust and stable cell-type attractors can be identified through measurements of the transcriptome to begin with.

In this work, we applied a classic approach that we term “*ϵ* networks” to a variety of single-cell genomics data to directly characterize their attractor structures. This approach allows us to analyze epigenetic spaces without the need to resort to nonlinear transformations and dimensionality reduction steps that can introduce distortion^42,43^. Using both simulated and experimental data, we validate our *ϵ* network approach and demonstrated its ability to detect separation using data from cell lines, and other biological controls, despite the high dimensionality of the data and the presence of noise. Yet, despite an extensive analysis of multiple experimental platforms, multiple gene sets, and multiple analytical pipelines, we found essentially no evidence of an attractor-like structure in available single-cell epigenetic data.

Rather than finding distinct cell states (“valleys”) separated from one another (e.g. by the “hills” on the landscape), we find that cells of very distinct phenotypes and lineages occupy more-or-less the same region of the space. Moreover, in all cases, we find that the data does not have the density distribution one would expect in the region of an attractor, with most cells near the bottom and fewer cells at the top (Figs. 1A and B). Instead, we find that the local density of cells in the epigenetic landscape varies across orders of magnitude and are distributed as an approximate power law^44^. The vast majority of cells are found in very low-density regions while a few cells are found in very high-density regions. As with our observations above, we found this “fractal-like density” distribution in all of the data we analyzed, including a large number of 10x scRNA-seq datasets, data from spatial transcriptomics platforms, data on protein levels, and data on chromatin accessibility.

Our extensive analysis reveals that essentially no single-cell data conforms to the predictions of Waddington’s landscape, at least in its contemporary interpretation where cell types are maintained by dynamical attractors in gene expression space^27^. There are several possible explanations for our findings. For one, a particular confluence of noise in technical measurements might simply prevent us from observing attractors that really are “there” in the underlying biological systems in question. This explanation would require that a vast array of different experimental modalities all suffer from so much noise that a robust attractor structure in real macromolecular levels cannot be observed in the actual experimental data, but this could nonetheless be the case. This scenario suggests an urgent need for the development of new experimental techniques that are accurate enough to characterize cell-type attractors. Alternatively, our findings raise the possibility that the mechanistic basis of canalization lies elsewhere, in a yet-to-be described epigenetic space, and a more principled approach could be developed to operationally map transcriptomes into meaningful cell types. Another possibility is that cellular phenotypes are more continuous than has been previously appreciated, such that there may be no attractors whatsoever, but rather a different form of cellular specification that admits incredible molecular and phenotypic diversity to enable organismal integrity. Regardless of which mechanistic hypothesis can better explain differentiation in development, this work suggests a critical need for the revisitation of our conceptual frameworks for understanding, analyzing, and interpreting emerging single-cell data.

## Results

### A lack of distinct cell groups in single-cell data

Our initial goal in this work was to characterize the variation of cells around the cell-type attractors in gene expression space. The first step in characterizing that variation is to identify these attractors themselves (Fig. 1A). We applied a classic approach, which we term “*ϵ* networks”, that iterates through a range of distance thresholds and measures how cells are distributed relative to one another in epigenetic space^45–49^. For any given dataset, we can calculate the distance between any two cells using their corresponding feature vectors (i.e. gene expression values). To start, we used the simple Euclidean distance (i.e. the *ℓ*^2^ norm) to define the distance, since this is definition of distance used in the vast majority of scRNA-seq and single-cell genomics studies.^50,51^ This is also the natural notion of distance used in the analysis of dynamical systems (see, e.g., the notion of “divergence” in the definition of the Lyapunov exponent, the standard definition of an attractor, and a host of other definitions and theorems)^52,53^. We should emphasize that the Euclidean distance applied to raw gene expression space is here just a starting point; we considered a wide array of other metrics and transformed versions of these datasets, as described below.

Regardless of the metric, to construct an “*ϵ* network”, we take an n-dimensional hypersphere of radius *∈*, where each dimension represents a gene, and we apply this hypersphere to each cell in the data (Fig. 1C, panels 1 and 2). If two cells are closer to each other than this radius *ϵ* (i.e., they lie within each other’s hyperspheres), then we connect those cells in the network (Fig. 1C, panel 3 and detailed in Fig. S2.1). For any given value of *ϵ*, we can thus convert our original high-dimensional dataset into a standard undirected graph. This allows us to use the powerful tools of graph theory to analyze networks at different values of *ϵ*, giving us insight into how the cells are distributed in gene expression space.

At any given value of *ϵ*, the network will naturally partition into different groups of cells that are all connected to each other, giving rise to a set of clusters or “components” (Fig. 1C, panel 3). If the cells are arranged in gene expression space according to the Waddington paradigm, we should see groups of similar cells in distinct regions of the space (see Fig. 2A for a schematic). As *ϵ* increases, cells of the same type should group together first; cells of different types should join together into the same cluster or component only after the radius *ϵ* exceeds the distance between the groups (Fig. S2.1iii). To track this behavior, we measured the size of the largest cluster in the graph (called the “giant component”) as a function of increasing *ϵ*. We should see that the giant component first includes only one of the cell types in the system (Fig. 2B). At larger *ϵ* radii, other groups of cells are expected to join all at once. This should give rise to a characteristic step-like behavior in the giant component (Fig. 2B). While the 2-D schematic in Figure 2A is useful, it is easy to prove mathematically that, if Waddington’s landscape is a good metaphor for the distribution of cells in epigenetic space, then there will be a discrete “jump” in the size of the giant component, regardless of the dimensionality of the data (see Supp. text 3, 4 and 5).

**Figure 2.**
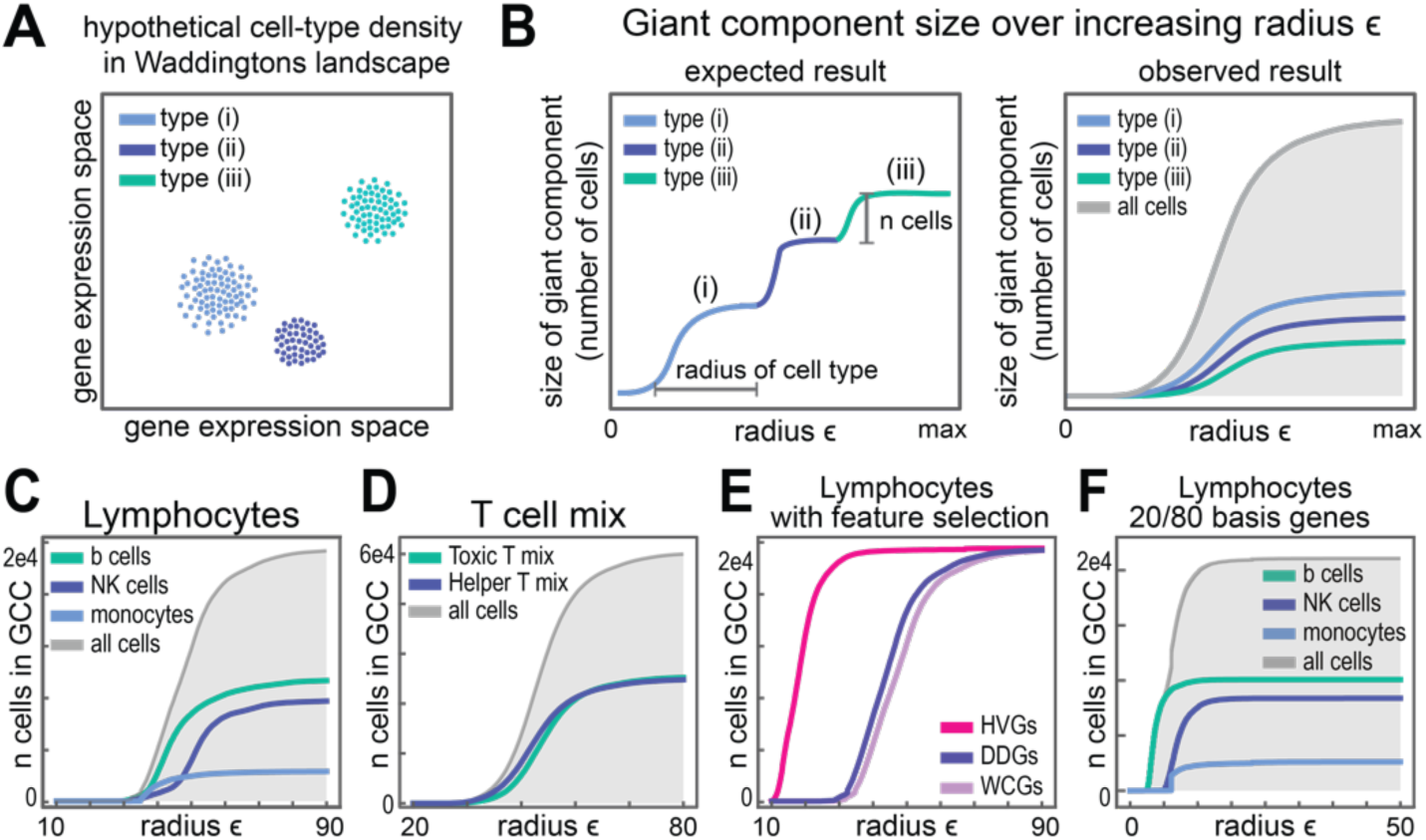
*∈* network analysis of FACS-separated lymphocytes. **A)** Schematic showing the expected distribution of cells in gene expression space, where each group exists in a distinct valley of Waddington’s Landscape and thus occupies a distinct region. **B)** Schematic showing how the size of the giant component changes as a function of *∈*. The expected behavior, illustrated on the left, demonstrates homogenous, dense groups of cells being sequentially added to the giant component in a step-wise fashion. On the right is an illustration of the observed behavior, where cells of different types occupy more-or-less the same region of gene expression space and join the giant component together. **C-F)** Size of the giant component as a function of *∈* for Lymphocytes sorted by FACS and then sequenced on the 10x genomics scRNA-seq platform. Giant component analysis using the raw UMI counts of all detected genes for **C)** cell types from distinct groups and **D)** cell types from different branches of T cells. In **E)** & **F)** b cells, NK cells, and monocytes, were analyzed using a subset of feature selected genes. **E)** uses either Highly Variable Genes (HVGs), Differentially Distributed Genes (DDGs), and genes determined to be different on average between the cell types using the Wilcoxon rank-sum test (WCGs). **F)** uses genes determined using a supervised, distance-based approach (20/80 basis genes).

We first applied this approach to a classic scRNA-seq data set from human Peripheral Blood Monocytes (PBMCs)^54^. In this data, cells were sorted by FACS using cell surface markers to separate them into distinct cell-type groups prior to sequencing. We thus know the “true” cell-type label for each cell in the dataset^55^, which allows us to not only monitor the size of the giant component, but also its composition. We first analyzed a subset of these data comprised of three cell types: B cells, Natural Killer (NK) cells, and monocytes. These terminally-differentiated cell types represent distinct developmental lineages of PBMCs, and should thus be easily separable in gene expression space. Yet, as can be seen from Fig. 2C, the size of the giant component (gray line) over increasing radius *ϵ*’s shows no step-like behavior whatsoever. Instead, the transition in the giant component is smooth, as cells of the different types join the giant component roughly at the same time (Fig. 2C). We see the same behavior when evaluating various subsets of T cells from the PBMC data set, suggesting that all the T cells purified in this experiment occupy the same region of gene expression space (Fig. 2D).

The findings presented in Fig. 2 clearly do not show the step-like behavior we expected to see, but there are many possible explanations for this. One simple alternative is the fact that this analysis was carried out on *all* the genes in the genome. The dataset in this case is likely to contain many genes that do not vary significantly between the different cell types, and it is possible this could mask the presence of separable groups in the data. Indeed, in scRNA-seq analysis, cell-type clustering is typically performed on only a subset of “feature selected” genes that are expected to vary significantly between different cell types. To address this question, we first considered the most popular approach of using “Highly Variable Genes” (HVGs) whose variance is higher than one would expect given the average expression level^56^. Interestingly, HVGs did not separate these lymphocytes into different groups (Fig. 2E). We also considered an alternative, more statistically grounded feature selection method that we recently developed (Differentially Distributed Genes or DDGs), and found that they similarly could not separate the groups of cells (Fig. 2E)^57^.

Interestingly, even when we employed *supervised* feature selection approaches, which take advantage of the fact that we know the cell types *a priori*, we could not separate the cells into distinct attractor-like groups. For instance, we used the Wilcoxon rank-sum test to find genes that are differentially expressed in one of the cell types compared to the other two (WCGs)^58^. Even these genes could not provide the expected step-like behavior (Fig. 2E). Indeed, a more detailed supervised analysis, which used a per-gene distance criteria, could not find a single gene that could be used to separate all three groups of cells with reasonable cutoffs (Figs. 2F, S2.2), and sets of genes selected using this approach did not result in well-separated groups (Fig. 2F). Regardless of which set of genes we used, we found that the populations of cells remain overlapping in gene expression space, with, for instance, many B cells more similar to monocytes and NK cells than they are to other B cells (Fig. S2.3).

### Validation of ∈ network sensitivity

Another possible explanation of the above findings is that the *∈* network approach itself is not sensitive enough to detect different groups of cells, especially given the high dimensionality, high levels of sparsity, and high levels of noise in the data^57,59–62^. The first issue we considered was dimensionality. An scRNA-seq dataset can easily have 10,000 or more dimensions, and one version of the “curse of dimensionality” states that, as dimensionality increases, distances become more similar to one another^63^. This result suggests that perhaps there really are distinct groups of cells in the data, but the distance metric we use cannot distinguish them because of the high dimensionality. We note that this issue with high dimensionality only holds when points are drawn from the *same* distribution in the high-dimensional space^63^. Waddington’s landscape^31,32,64,65^, however, clearly posits that cells of different types are drawn from *different* distributions in epigenetic space. It is straightforward to prove mathematically that this curse of dimensionality does not hold when data is drawn from different distributions, and so high dimensionality is not *a priori* a problem for the approach used here (Supp text 3).

To empirically test the sensitivity of *ε* networks in high-dimensional spaces, we first generated a set of synthetic datasets in 10,000 dimensions, particularly a Negative Binomial Mixture Model (NBMM) and a Poisson Mixture Model (PMM) and a Gaussian Mixture Model (GMM). In both cases, we designated a subset of 250 genes to be “marker” genes that are highly expressed just in one “cell type,” resulting in a total of 750 marker genes. The remaining 9,250 genes were expressed at a low level in all cells. We found that *∈* networks could detect separation across a wide range of parameters (Fig. 3A, Figs. S3.1-S3.3). We saw similar results for a simple Gaussian Mixture Model (GMM) as well (Figs. S3.1).

**Figure 3.**
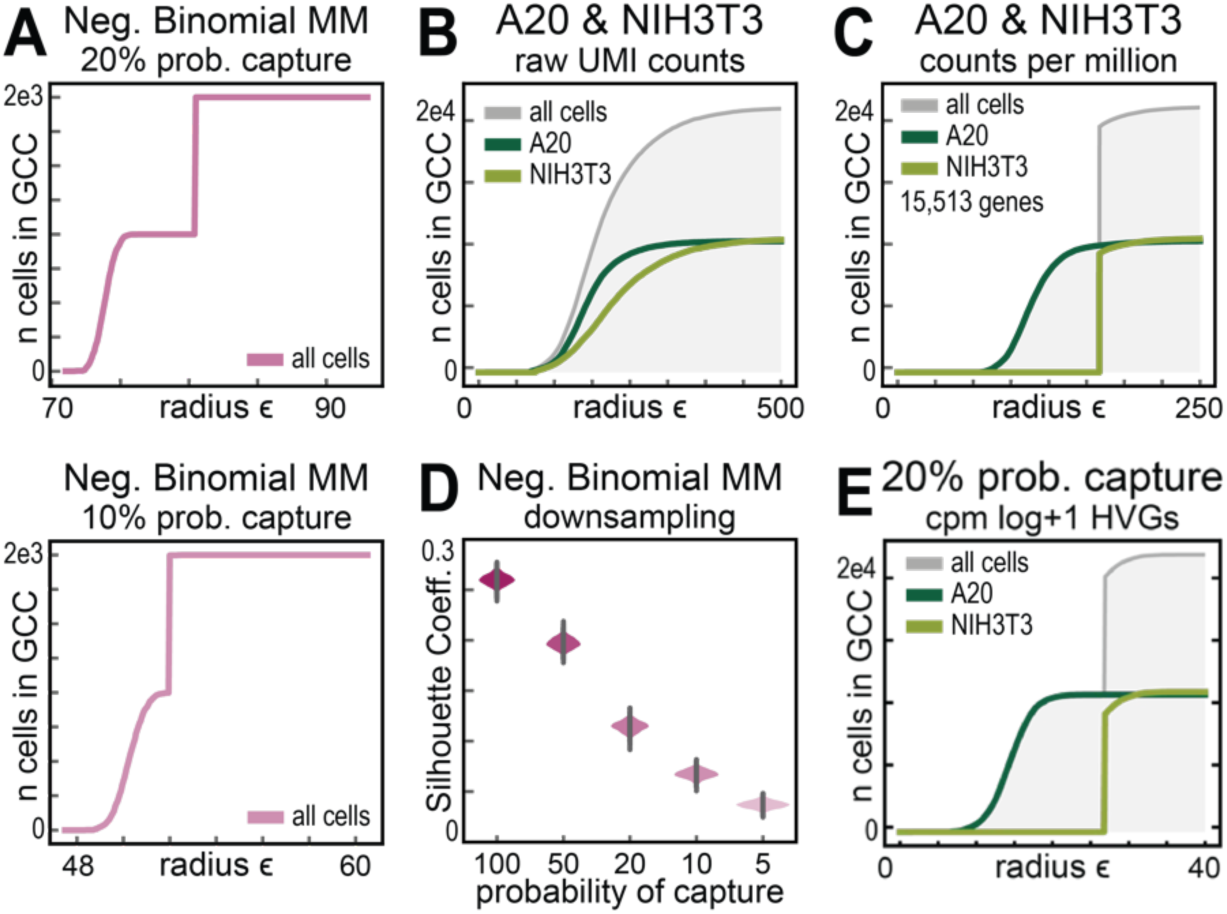
*∈* network analysis can detect separation despite high-dimensionality, sparseness, or noise. **A)** Size of the giant component as a function of *∈*, when *∈* networks were constructed from simulated data, where the values for 10,000 genes were sampled from negative binomial distributions. For each of the three panels, a mixture of two cell type are simulated, each of which only differ by the expression levels of 250 genes. These ‘marker gene’ values are sampled from a negative binomial distribution with the indicated means. **B,C)** Size of the giant component as a function of *∈* for *∈* networks generated from B) the raw UMI counts of multiplexed cell line data, or C) the counts-per-million transformed counts of multiplexed cell line data, where all 15,513 genes are included in the graph. **D)** Silhouette coefficients for the marker gene titration illustrated in (A). **E)** Size of the giant component as a function of *∈* for *∈* networks generated from the CPM log+1 transformed HVGs of a downsampled version of the cell line dataset.

To determine whether *∈* networks could identify separation in real, high-dimensional biological data, we applied our approach to scRNA-seq data from cell lines, which we reasoned would have the highest chance of showing separation. For this analysis we used multiplexed 10x scRNA-seq data that was generated from A20 and NIH3T3 cell lines, where again we know the identity of each cell *a priori*^66^. When we applied our *ε* networks to the raw UMI count data, we were unable to detect separation even in this dataset (Fig. 3B). However, when we applied simple counts per million normalization to this data, we did observe separation, even when we measured the distance between cells using the Euclidean distance between all 15,513 genes in the dataset (Fig. 3C).Taken together with our results on various Mixture Models, these results demonstrate that *ε* networks are indeed sensitive enough to detect separation even in exceedingly high-dimensional datasets.

Another possible explanation for our observation is that the experimental platforms used to generate scRNA-seq data are highly noisy, and result in data that is “sparse” in that most genes are not expressed in most cells. The combination of noise and sparsity could prevent the *ε* network approach from detecting the separation between cell types even if such separation exists in the original cells. To explore this possibility, we first used our synthetic data from the NBMM, since NB distributions are similar to the count distributions observed in scRNA-seq data^67–69^. To model noise due to “dropouts” in library preparation, we included a Binomial sampling process where each mRNA originally present in any given simulated cell had a small chance of being captured and retained in the final dataset (further detailed in the methods section). In these models, we found that the *ε* network approach found separation even when average gene expression levels, after the dropout sampling process, were relatively low (less than 10%) (Figs. 3A and 3D, S3.2 A-C). When noise is so extreme that the averages approach 0, then we can no longer observe separation, because there is essentially no data left (Figs. S3.2 A-C).

Intriguingly, CPM normalization alone does not generate separation in the lymphocyte data considered in Fig. 2 (Supp. text 5, 6 and Fig. S5.1), or in essentially any other dataset we considered (see below). So, while the cell line data provides a positive control, it is still possible that real separation in biological data could be obscured by low sampling rates during the measurement process. To address this possibility, we down-sampled the multiplexed A20 and NIH3T3 cell line data by performing a Bernoulli trail for each UMI count in the data, randomly retaining only a fraction of the counts in the original data. Even if we throw away 50% of the counts in this (already sparse) dataset, we still see robust separation between the groups; indeed, separation is still detectable (if weaker) when we remove even 80% of the data (Fig. S3.3A). If we quantify sparsity of as the fraction of “0” counts in the data or the number of UMI counts per gene per cell, this 50-80% down-sampled version of the A20/NIH3T3 dataset covers the range of other datasets we considered in this work (Fig. S3.3 D-I). Indeed, when we consider the more highly transformed version of these downsampled datasets, we see separation in the data even in cases where the sparsity is much higher than the vast majority of datasets considered here (Fig. 3E, TABLE S4). Taken together, this suggests that the *ϵ* network approach can detect separation even in noisy, high-dimensional data, and that many scRNA-seq datasets should have sufficient data quality for these networks to detect separation, if it is present.

Another possible explanation for a lack of separation could be that the data forms multiple separate groups that simply overlap on their edges. In this scenario, the main groups of the cells would be separated, but *ϵ* networks would not be able to detect that due to a few cells from each cell type that just happen to be near cells from the “edge” of the basin of cells from another type. To test this possibility, we calculated the “betweenness centrality” for all the cells in the dataset. This measures the extent to which the shortest paths between nodes use any given node in the graph. Peripheral bridge cells should have high betweenness centrality, since they connect two large groups of cells, but fewer neighbors, since they are on the edges of the cluster. We do not see cells with few neighbors but high betweenness centrality in this case, and visualization of the corresponding *ϵ* networks also shows that the cell-type groups are not connected on their edges, but rather entirely enmeshed (Fig. S3.5). Interestingly, in the A20/NIH3T3 dataset, when we down-sample by 80%, we actually do see these “bridge” nodes emerging, indicating that the lack of separation between these two groups is due to two separate groups starting to merge on their peripheries (Fig. S3.3). The lack of such behavior in the majority of scRNA-seq datas suggests the groups are fundamentally overlapping, consistent with our findings from Fig. 2.

### A lack of distinct cell groups in single-cell data across modalities and organisms

All the data used above was obtained using the popular 10X genomics platform. Although the NIH3T3/A20 and simulation data shown in Fig. 2 suggest that the lack of separation in the lymphocyte data is not purely due to noise and sparsity inherent in the platform, it could still be that biased sources of noise in the 10X approach end up destroying the real separation between cell type groups^70^. To test this possibility, we next analyzed many different types of data taken from different measurement approaches, including scRNA-seq techniques that have much higher capture probabilities. For instance, the BD Rhapsody platform captures 50-75% of mRNAs in each cell, compared to ∼35% for modern 10X approaches, although it can only provide data for ∼1,000 genes in the genome^71^. We applied our analysis to BD Rhapsody data for PMBCs, and again found no evidence of distinct cellular groups (Fig. 4A)^71^. We also considered data from Parse Biosciences, a plate-based platform that uses combinatorial barcoding, as well as data from the 10X version 3 chemistry with improved sensitivity^72,73^. Even when we applied feature selection methods to these data, we were unable to find evidence of distinct cellular groups (Fig. 4A).

**Figure 4.**
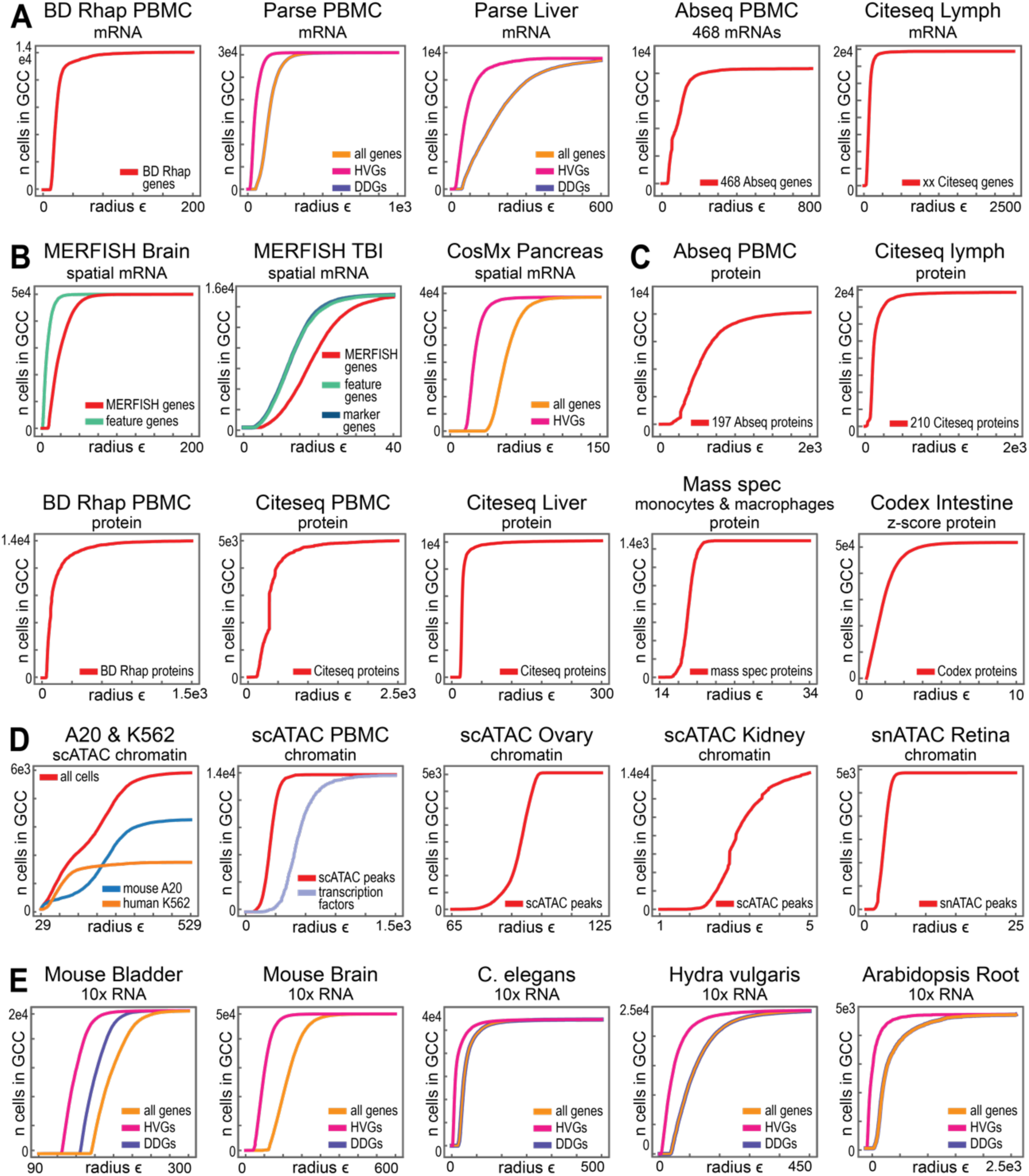
*∈* network analysis of data from different single-cell measurement platforms. **A-E)** Size of the giant component as a function of *∈* for data generated using targeted and genome wide **A)** RNA-seq platforms, **B)** spatial transcriptomics platforms, **C)** single-cell proteomics platforms, **D)** single-cell chromatin accessibility platforms, and **E)** 10x single-cell RNA seq experiments from different phylogenies. Additional details of each dataset are available in Table S1.

Alternative approaches have also been developed that allow for the quantification of mRNA levels in single cells that do not rely on the “capture-and-sequence” paradigm. For instance, the MERFISH technique uses a combinatorial set of labeled probes to identify individual mRNA molecules from a subset of genes in the genome in a microscopy image^74^. We analyzed MERFISH data consisting of a mouse brain slice with ∼50,000 cells with measurements for around ∼600 genes in each cell (Fig. 4B)^75^. We also analyzed another MERFISH dataset of around 15,000 mouse brain cells following a simulated Traumatic Brain Injury (TBI). This data contains measurements for 160 distinct genes per cell, of which 80 genes were *specifically selected* as marker genes for cell types^76^. We found no evidence of distinct cell type groups in either the entire TBI dataset or a dataset in which we just considered these 80 marker genes (Fig. 4B). To further test whether appropriate subsets of “cell-type determining genes” could separate groups of cells, we identified genes in the two MERFISH datasets with apparently more multimodal distributions (Fig. S4.1A), reasoning that these genes might comprise different sets of genes with distinct expression levels in different subsets of cells. However, using these sets of 23 “feature genes”, we again found no evidence of distinct cell type groups (Fig. 4B,C). Together, these findings strongly suggest that our observations are not a consequence of either low capture probabilities, noise, or any property specific to a given experimental platform.

We also considered additional epigenetic modalities, including single-cell measurements of proteins (Fig. 4C) and scATAC-seq (Fig. 4D), which provides genome-wide information on chromatin accessibility^71,77–80^. *ε* network analysis of protein levels from the BD-rhapsody, Ab-seq, cite-seq, mass-spec, and CODEX platforms did not recover any distinct groups (Fig. 4C, S4.2B,C). scATAC-seq data from PBMCs, Ovary, Kidney, Retina, Fallopian tube, and Heart data yields essentially identical results to those obtained from scRNA-seq (Fig. 4D, Fig. S4.2D). Because scATAC-seq assays DNA accessibility, rather than mRNA levels, we inspected whether evaluating just the subset of genes that encode transcription factors would recover distinct groups of cells. Once again, considering only transcription factors did not produce distinct regions of cells (Fig. 4D). Thus, regardless of the single-cell platform, epigenetic modality measured, or subset of genes considered, we could not find any published single-cell dataset on epigenetic state that was consistent with well-separated attractors for individual cell types as predicted by Waddington’s landscape (Fig. 1A and 2B).

We next extended our analysis to characterize scRNA-seq data from samples with distinct developmental and phylogenetic origins. In each case we constructed *ϵ* networks for the original, high dimensional data and different subsets of genes. We first analyzed data from the mouse bladder, as an example of a tissue with moderate complexity, and found no evidence of step-like behavior in any of the feature selected data (Fig. 4E)^81^. Similarly, regardless of how we selected the feature space, we saw a smooth transition in the much more complex tissue, the mouse brain (Fig. 4E). We then considered scRNA-seq data from the developing *C. elegans* embryo^82^. Again, we saw the same behavior (Fig. 4E), despite the deterministic nature of *C. elegans* development, where the distinct cellular fates of various lineages are extremely well-characterized^83^. This behavior persists even when we just look at transcription factor genes in *C.* elegans, which are thought to drive fate transitions within these lineages and thus should show some evidence of the expected attractor structure (Fig. 1A and S4.1D). We saw the same behavior for every data set we considered, including whole-organism data from *Hydra vulgaris* (Fig. 4E) and even data for the model plant *Arabidopsis thaliana*, suggesting that this is a feature of all complex multicellular organisms (Fig. 4E)^84^.

### Popular non-linear transformations cannot reliably separate groups of cells

In the scRNA-seq field, analyses like cell-type clustering are essentially never performed on the raw data, but rather only on data that has been subjected to a set of non-linear transformation and dimensionality reduction steps^50^. A typical pipeline for scRNA-seq data would start with the raw UMI counts, perform “counts per million” (CPM) or an equivalent normalization, perform a log transformation (e.g. log (CPM +1)), identify a set of HVGs, perform PCA on the log (CPM+1) transform of that subset of genes, and then use non-linear techniques like t-SNE or UMAP to visualize the data in two or three dimensions ^50^. Clustering is usually carried out using the Louvain or Leiden community-detection algorithms after either the PCA or tSNE/UMAP step^50^. Given that many groups report successful clustering of their data using this broadly similar set of approaches, we considered whether these transformations could reliably separate cells into distinct groups and, if so, which parts of this pipeline seem critical for that separation.

As above, we first considered the case of the FACS-separated Lymphocyte data. Interestingly, we found that most combinations of transformations in this pipeline did not result in distinct cell groups (including CPM normalization on its own, PCA on its own, log transformation, or selection of HVGs, Figs. S5.1). This is particularly interesting, since CPM normalization on its own was sufficient to generate separation in the NIH3T3/A20 cell line data (Fig. 2). Combining all of these transformations did, however, generate the step-like behavior we originally expected to see, regardless of the set of genes the pipeline of transformations was applied to (Fig. 5A). This is encouraging, suggesting that, while there are not distinct attractors in the raw count data, applying this popular pipeline of transformations might be able to recover distinct groups. While application of these transformations to control cell line data was able to produce separation (Fig. 5A, S5.2, S5.3), application of these transformations to data from intact biological organisms, however, produced either no separation between cell groups or very little step-like behavior (Fig. 5A, S5.4, S5.5, S5.6, S5.7, S5.8). For instance, when applying the standard analysis pipeline to the mouse bladder, the transformed data gives essentially the same pattern as the raw data, regardless of the feature selection technique employed (Fig. 5A and S5.4A). This suggests that the standard pipeline is not a universally reliable approach to generate separation.

**Figure 5.**
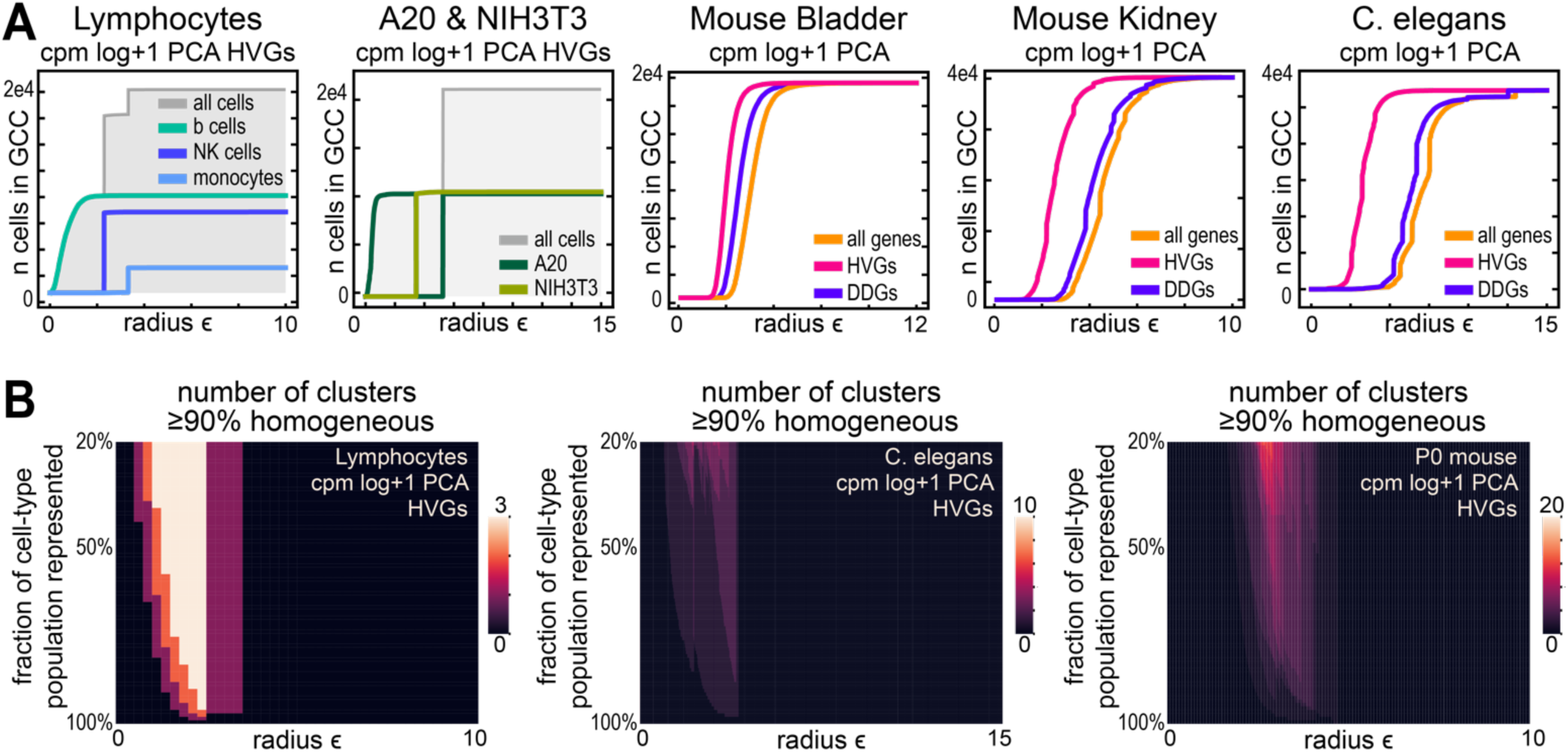
Common non-linear transformations do not recover cell-type groups. **A.** Size of giant component vs *ϵ* for 10x scRNA-seq data generated from FACS sorted Lymphocytes, multiplexed A20 & NIH3T3 cell lines, mouse bladder, mouse kidney, or *C. elegans* after Counts Per Million (CPM) normalization, log CPM + 1transformation, selection of Highly Variable Genes (HVGs) or Differentially Distributed Genes (DDGs), and dimensionality reduction using PCA. **B.** Heatmap characterizing the size and composition of components in the *ϵ* networks for the FACS sorted lymphocytes, the *C. elegans* data, or data from the “one well” P0 embryo from the Mouse Atlas. The x-axis indicates increasing radius *ϵ*, the y-axis represents the fraction of particular cell types represented in each cluster, relative to the total number of that particular cell type in the data. The color of the heatmap indicates the number of components meeting a 90% homogeneity criteria. For the Lymphocyte data, we see a range of *ϵ*’s where 3 components are more than 90% homogeneous and have collected nearly 100% of the cells of that type. In *C. elegans* and P0 mouse, we see very few cases where clusters that are 90% homogeneous contain a large fraction of the cells of that type.

Most of the other data sets we considered fell in between the behavior seen for PBMC lymphocytes and the mouse bladder (Figs. 5A, S5.4, S5.5, S5.6, S5.7, S5.8). For instance, for the *C. elegans* embryo, we do see some distinct jumps, but not the characteristic step-like behavior we expect for well-separated cell type groups (Fig. 5A and S5.4C). In most data sets, we lack orthogonal cell-type annotations, so we cannot tell if these individual “jumps” correspond to single cell types, or groups of cell types, joining the giant component all at once (as is the case for the lymphocytes, Fig. 5A). To test this possibility, we focused on the *C. elegans* data, where we have operationally standard annotations for the cell type of each cell in the data set^82^. At each value of *ϵ*, the graph will contain multiple components or clusters (Fig. 1B). The moderate step-like behavior we observe here could be due to the fact that cell type groups are coalescing into separate clusters, joining one another, and then adding to the giant component all at once. To test this, we quantified the composition of all of the clusters in the graph as a function of *ϵ*, not just the giant component.

In the transformed *C. elegans* data, we see very few components that are both highly homogeneous and contain most of the cells of a given type (Fig. 5B, S5.9D, and S5.10 A,B). For instance, we never see more than 5 clusters that contain more than 50% of the cells of any given annotated cell type, regardless of the *ϵ* value we consider. This is in contrast to the PBMC data, where we do indeed see three distinct clusters that mostly consist of a single cell type and that collect most of the cells of that type in the dataset (Fig. 5B, S5.9B). Given that there are 36 annotated cell types in the *C. elegans* data, this suggests that, even though we see discrete “jumps” in the sizes of the giant component for many data sets, this does not correspond to clusters of single cell types first joining together with one another, and then with the giant component. Instead, we see small isolated “islands” of cells of similar type, but those islands are generally closer to cells of a different cell type than to other islands of cells of the same type (Fig S5.11). We extended this approach to characterize the cluster composition of data generated from Hydra as well as a single embryo from the Mouse Atlas, again using the operational standard cell type annotations^85^. In both additional experiments, the standard set of transformations fail to recover a large clusters of homogenous cell type groups (Fig 5B, S5.9F). Thus, while the transformations applied here do seem to generate small groups of “similar cells,” they certainly do not result in the expected attractor structure predicted by Waddington’s landscape.

Interestingly, in a few cases, such as the 10x v3.1 PBMC data, we observe that CPM transformation of HVGs alone (i.e. without log transformation or PCA) produces distinct jumps in the *ϵ* network analysis (Fig. S5.8, and S5.12). To distinguish between the possibility that the two groups of cells clusters correspond to distinct PBMC lineages, or the possibility that CPM-HVG transformation artificially introduces separation in the data^24^, we took advantage of the well-established marker genes in this tissue and evaluated marker gene expression across the two groups. Analysis of gene expression in the two clusters suggested that, in the v3.1 PBMC data, the two groups produced by CPM-HVG transformation appear to correspond more to “read-depth” than any pattern of marker gene expression (Fig S5.12). Indeed, CPM transformation of a selection of 34 random genes also produced two distinct clusters in this data (Fig S5.12D), raising the possibility that the generation of apparent clusters can occur spuriously when non-linear transforms are applied (Supp. text 6)^61^.

Since the standard pipeline or CPM-HVG transformation does not consistently separate groups of cells, we evaluated whether other common transformations of the data could produce the expected separation. It is also often argued that due to differences in relative expression levels, different genes in the dataset will have intrinsically different scales of variation, and thus a vastly different impact on the distance. One simple way to deal with this problem is to perform a z-score transformation of the data^50^.

However, z-score transformation failed to generate separate groups of cells for any of the feature-selected single-cell data we tested (Fig. S5.13). Further, neither the combination of CPM then z-score normalization, or vice-versa, succeeded in separating data into distinct groups, regardless of which set of genes were tested (Fig. S5.14).

In addition to accounting for “sampling differences”, cell-size, and gene weights, we also tested a number of alternative feature spaces and clustering techniques. We tested cell-cycle regression methods^86^, different metrics of distance (i.e. the *P*^1^ and other norms), as well as non-single linkage clustering algorithms (Figs. S5.15, S5.16, S5.17, S5.18, and S5.19). However, across all of the single-cell data we tested, none of these various scaling, measurement, or clustering techniques produced the expected separation of cells into distinct groups.

### The density distribution of cells is inconsistent with attractors in epigenetic space

In addition to being separated from one another, Waddington’s picture predicts that cell types should correspond to attractors in epigenetic space ^32,87–89^. A key property of an attractor is the fact that, if a cell is perturbed away from the attractor by noise or an environmental fluctuation, the natural dynamics of the gene regulatory network will induce the cell to move back towards the attractor^52,53^. This is usually represented using a “potential well” picture, which explains how cells near an attractor will tend to return to that attractor if they are perturbed while cells in unstable regions (say, at the top of a hill between two attractors) will tend to move away (Fig. 1B). In the presence of noise, the structure of these potential wells suggests that the probability of finding a cell in a certain region of the landscape should vary with the height of the landscape, leading to a large number of cells near the attractors at the bottom of the valleys and few cells on the hills in between them (Fig. 1A and S6.1). Indeed, this principle has been used to attempt to infer the shape of the landscape from scRNA-seq data^32^. Waddington’s landscape thus leads us to expect that most cells will be in regions of similar density, with similar numbers of neighbors in the *ϵ* networks we constructed.

To investigate this phenomenon we calculated the “density distribution” for various single-cell datasets. For any given value of *ϵ*, we can count how many neighbors each cell has (Fig. 1C), and then plot a histogram of this data. In graph theory, this histogram is known as the “degree distribution” of the network, and is often instructive about the topology of the network itself^90,91^. In this work, we will refer to this distribution of connectivities as a density distribution, as in most practical cases it measures the number of neighbors a cell has within the volume of the *ϵ*-sphere around that cell. We should note that, since UMI counts can only have non-zero values, the exact volume of “allowed” gene expression space may vary from cell to cell, and so this notion of degree as a measure of density is approximate. If most cells are near the center of an attractor (the bottom of the well in Fig. 1B), they should have more-or-less the same number of neighbors, which should give rise to a roughly binomial (i.e. approximately Gaussian or Gaussian-like) distribution of neighborhood sizes (Fig. 6B).

**Figure 6.**
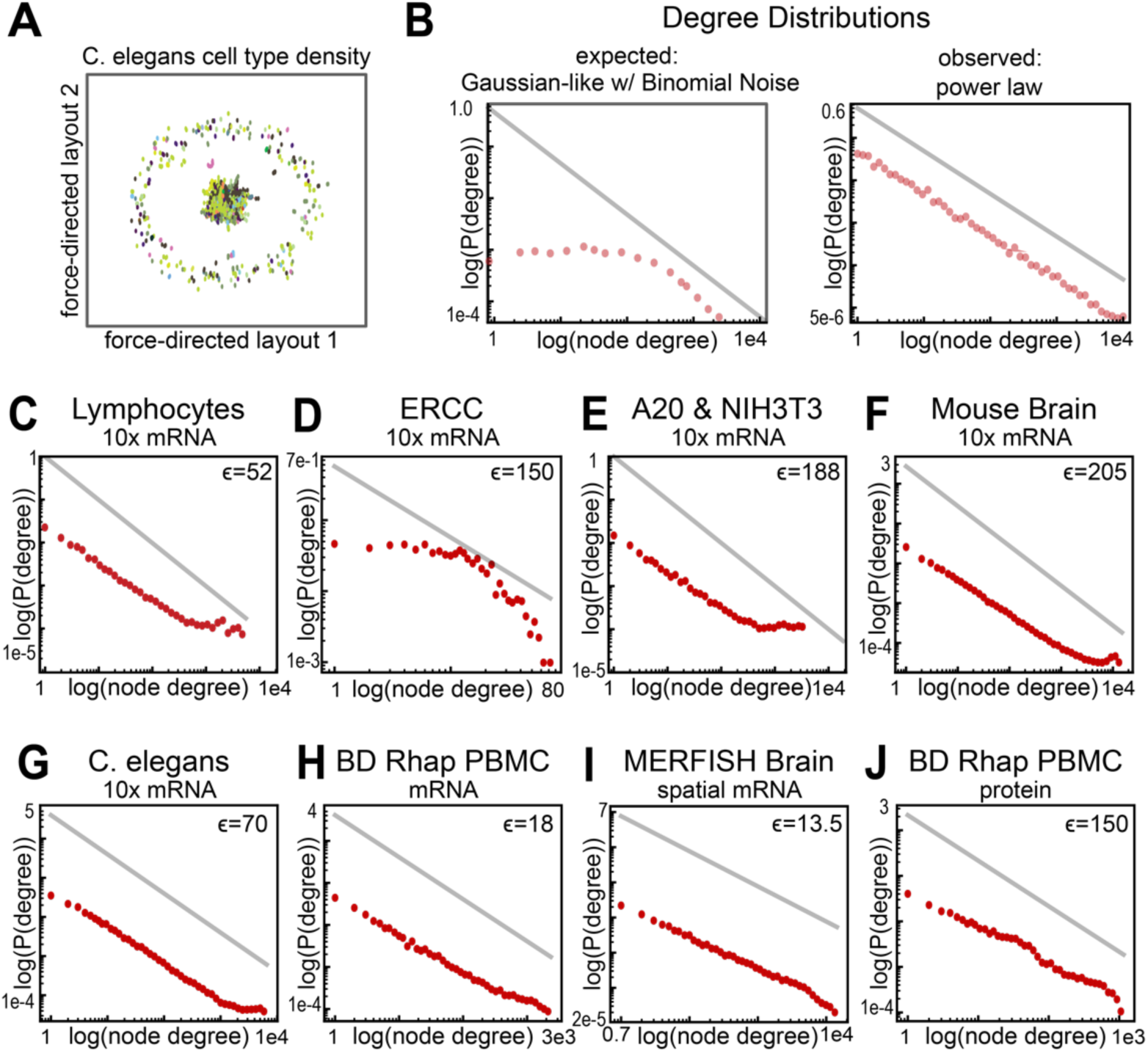
Density distributions of single-cell datasets indicate the absence of attractor structures. **A.** Force Directed layout of the *∈*-network (*∈* = 70) for data from *C. elegans* embryos collected on 10X platform (all “orphan” cells with no neighbors were removed for visual clarity). Cells are colored according to the annotated cell type. **B.** A schematic of the degree distribution expected to be produced by attractors (left) vs the degree distribution expected when sampling from a power law distribution (right). **C-J)** Degree Distribution for the *∈*-network of various datasets at the indicated *∈*: C) FACS-separated Lymphocytes D) ERCC control data E) A20 & NIH3T3 cell lines, F) Mouse Brain, G) *C. elegans,* H) BD Rhapsody PBMC mRNA, I) MERFHISH Brain, J) BD Rhapsody PBMC protein. The gray line represents a power-law with an exponent of −1 for reference.

We first considered the 10X PBMC FACS purified lymphocyte scRNA-seq data described above and found an approximately power-law distribution of local densities (Fig. 6C). Although the density distribution shown in Fig. 6C is only for one particular value of *ϵ*, we find the same scaling behavior across a wide range of *ϵ* values, suggesting that this is not simply an artifact of choosing precisely the right radius for the neighborhoods in question (Fig. S6.2A). In this case, we find a small number of cells that are in extremely dense regions of gene expression space, with thousands of neighbors; most cells, however, are found in very low-density regions, with either no neighbors or just one or two (Fig. 6C). The *ϵ* networks we observe are thus similar to classical “scale-free” networks, though we should note that our analysis here is insufficient to determine if these distributions are truly scale-free or simply similar to a power law^92^. Regardless, this highly heterogeneous density distribution is completely inconsistent with the distributions we would naturally expect to see in the neighborhood of a stable attractor (Fig. 6B). Indeed, in all our our Negative Binomial Mixture Models, we observe the expected gaussian-like distribution, regardless of the amount of sampling noise in the experiment (Fig. S3.2).

As mentioned above, the 10X platform is known to be noisy, so it could be that the heterogeneous density distribution we observe is a reflection of the platform and not the underlying biology. To test this, we considered an artificial dataset in which a set of 92 cDNA standards were introduced to the 10X platform at defined concentrations that span orders of magnitude^54^. In this “ERCC control” data, we do not observe any evidence of scaling behavior, suggesting that powerlaw densities are not simply an artifact of the scRNA-seq technique (Fig. 6D). Interestingly, we observed highly heterogeneous density distributions for all the 10x data sets discussed above, including the A20 & NIH3T3 cell line data (Fig. 6E), mouse brain (Fig. 6F), *C. elegans* (Fig. 6G), *H. vulgaris*, *A. thaliana,* etc. (Figs. S6.3A). Interestingly, downsampling the A20& NIH3T3 cell line data did not reduce the extend of scaling behavior (Fig. S3.3B). We also observe similar density distributions in the BD Rhapsody PBMC data (Fig. 6D), further suggesting that this observation is not an artefact of the low capture probability entailed by the 10x platform.

To test whether this observation was specific to the scRNA-seq paradigm, we also analyzed several available spatial transcriptomic datasets. Interestingly, the Vizgen MERFISH data on the mouse brain showed striking power-law behavior across four orders of magnitude in local densities (Fig. 6I). We saw similar behavior for the TBI (considering either the entire dataset or just marker genes) and cultured MCF10A MERFISH data, (Fig. S6.3C) and CosMx genome-wide spatial mRNA data from the Pancreas (Fig. S6.3C)^93^. As with our analysis of the giant component, these findings strongly suggest that these power-law densities are not a simple consequence of either the high levels of noise nor the inherently high dimensionality of scRNA-seq data. The fact that this observation holds across quite different subsets of genes (chosen in each data set for completely different reasons), as well as in feature-selected scRNA-seq data (Fig. S6.4-S6.9), also suggests that power-law densities are a general feature of the distribution of cells in gene expression space regardless of the subset of genes considered^71,76,81,82,84,93,94^. Interestingly, we also see this scaling behavior in other epigenetic datasets, including all of the protein data (Figs. 6J, S6.3D) and scATAC-seq data, including when we only consider the subset of transcription factors (Fig. S6.3C), and if we use the *ℓ*^1^ norm or the Hamming distance instead of the standard *P*^2^ norm^95^ (Fig. S6.11, S6.12). Further, we applied various normalization and dimensionality reduction techniques to feature and non-feature selected data to test whether these transformations could recover locally dense regions of cells. While some transformations altered the curvature of the distributions, none of the approaches recovered gaussian-like density distributions (Fig. S6.4-6.10, S6.13). As with the lack of separation between cell types, non-gaussian density distributions are thus a universal feature of the single-cell data we analyzed.

## Discussion

For over 80 years, Waddington’s landscape has been the dominant picture used to explain canalization in ontogeny^28,29,32,87,89,96^. Since the 1960s, the near-universal interpretation of this landscape has been that individual cell types should correspond to attractors in gene expression space. Indeed, the landscape picture itself was influential in developing the language used in modern dynamical systems theory (for instance, the notion of an attractor’s “basin of attraction” was directly inspired by Waddington’s ideas)^28,32,87,89,94,96^. Mathematical models of development and cellular differentiation universally cast the process as a set of bifurcations, whereby the number of attractors in gene expression space change during development, constraining and producing different phenotypic fates^26,28,32,87,89,97^. These concepts are deployed extensively in the study of cancer, stem cells and stem cell reprogramming, and other areas^28,87,89,97^. Despite its widespread acceptance, it has only been recently that single-cell measurements have allowed us to directly test this picture.

The predictions of Waddington’s landscape are clear: cell types should be drawn from different distributions in gene expression space, corresponding to the different attractors on the landscape (Fig. 1A, Supp text 4 and 5)^26,28,32,87^. Our analysis demonstrates that most available single-cell measurements are inconsistent with these predictions. For one, rather than occupying distinct regions of gene expression space, or even slightly overlapping regions of space, cells of very distinct types and lineages occupy the same region of that space (Fig. 2, Fig. 4, Fig. S4.2), regardless of how we select the subset of genes on which to focus our analysis. We found this not just for scRNA-seq data taken from the 10X platform, but also for more targeted technologies that provide higher-quality data for an ostensibly biologically informative subset of genes (e.g. the MERFISH data from mouse brains in Fig. 4B). This lack of separation is also present in chromatin accessibility data and data on protein levels (Figs. 4C,D).

Moreover, the other clear prediction of Waddington’s landscape, that the cells should be found in attractor states no matter how well separated they might be, also does not conform to our findings. Instead of seeing cells clustered around a “typical” gene expression state corresponding to the center of an attractor, we find that cells are extremely heterogenous, leading to an approximately power-law or “fractal-like” density distribution (Fig. 6 and Fig. 7A).

**Figure 7.**
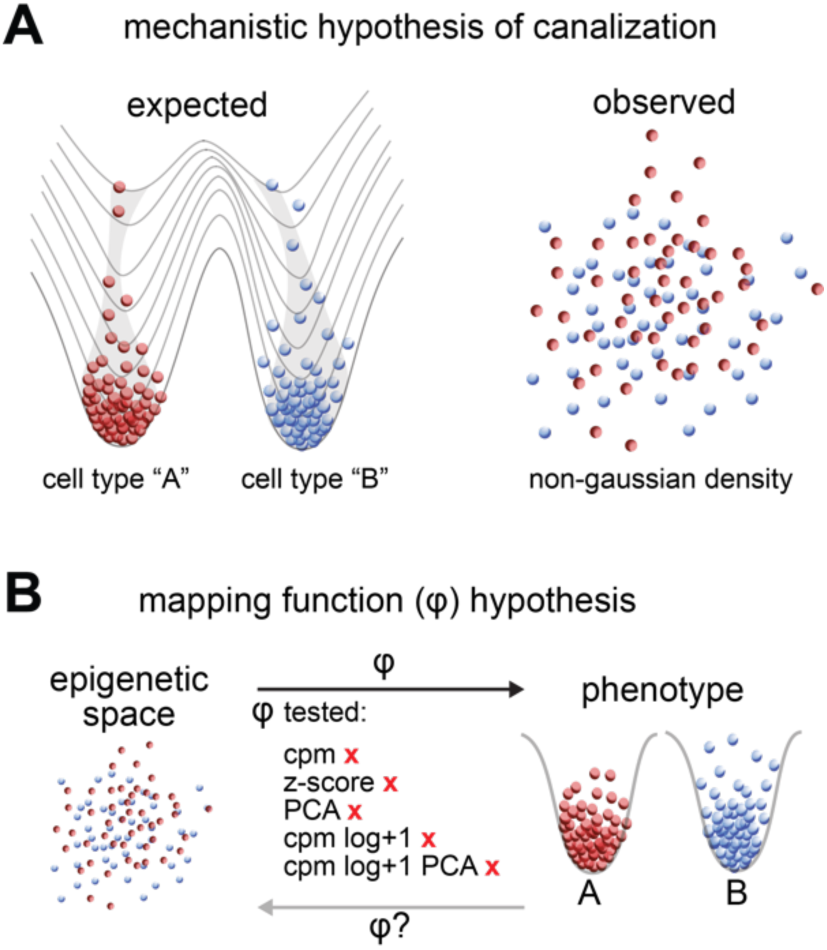
Current single-cell epigenetic data does not support our current conception of cell types. A) Schematic illustrating cell-type densities predicted from the mechanistic hypothesis of canalization depicted in Waddington’s Landscape compared to the observed non-gaussian densities. B) Schematic illustrating the inability to find a function that maps transcriptome space back into a phenotypically separable cell-type space.

Since its inception, the analysis of single-cell genomics data, and particularly scRNA-seq, has relied on a series of nonlinear transformations and dimensionality reduction steps that are applied before any attempt is made to cluster cells into cell types^37,50^. In order to make sense as a biochemical explanation for differentiation and development, the state variables of the dynamical system schematized in Waddington’s landscape should represent the concentrations of the mRNAs and proteins (see Supp text 5)^11,26,29,32,87,89^. As such, we should see separation directly in the space of mRNA (or protein) levels, or potentially in the CPM data (see Supp text 5 & 6), which we clearly do not see across most datasets. Nonetheless, nonlinear transformations have been operationally useful in the analysis of scRNA-seq, since no one can deny the sheer volume of papers that have derived meaningful biological insights from this data. We tested the hypothesis that this “standard pipeline” has discovered a way to approximate a function that maps the epigenetic state to phenotype, and thus recovers physiologically distinct groups of cells (Fig. 7B). We found that application of the standard pipeline of scRNA-seq analysis sometimes does separate cells into discrete, homogenous groups (Fig. 5A). In the vast majority of cases, however, this approach simply does not “work:” these transformations generally project groups of physiologically distinct cells into essentially the same region of state space, with highly heterogeneous density distributions (Fig. 5, Fig. 6 and Fig. S5.2-S5.7).

One way to explain our results is to posit that the “real,” underlying epigenetic spaces of cells actually do have an attractor-like structure, but essentially all experimental tools currently used are so noisy that they destroy that attractor structure. Even though our *ε* network analysis clearly demonstrates that the standard transformations do not reliably invert this sampling process (Fig. 7), it could be that popular clustering approaches like Louvain and Leiden detect subtle differences in gene expression patterns and thus detect residual signatures of the true Waddington-like structure that was destroyed by the experiment. Interestingly, scRNA-seq analyses can be very difficult to replicate, and in some studies, even with considerable effort, the authors cannot cluster the data into reasonable, distinct groups of cells^43,98^. Indeed, extremely small perturbations to a dataset, such as removing a few hundred cells, results in radically different cell type clusters (Fig. S1, echoing recent results that demonstrate that even changing the random seed of the clustering algorithm can significantly change the results^40^. This lack of statistical robustness indicates that, if there is a true cell type identity for any given cell in the dataset, Louvain and Leiden cannot reliably recover that identity. Interestingly, both Louvain and Leiden are algorithms that detect “modules” in the shared *k*NN graphs generated from the data^99,100^. When we look at Force-Directed Layouts (FDLs) of these graphs, we immediately see that there are few obvious modules in the data (Fig. S1). This is because the cells essentially completely overlap in the underlying transformed space, making the problem of identifying clusters challenging. Interestingly, when we take a case like the A20/NIH3T3 cell line datasets, where there is actual separation in the underlying space (Fig. 3), clustering results are instead robust to small perturbations in the set of cells we include in the analysis (Fig. S1). Our work thus suggests that persistent difficulties with the reliability and reproducibility of cell type clustering likely arise from the lack of underlying cluster structure in the data itself.

We should note that our findings should not be taken to suggest that clustering approaches in scRNA-seq cannot provide meaningful insights into the data. It is clear that a combination of nonlinear transformations, dimensionality reduction and graph clustering can generate useful models of cell types, especially when guided by a user who can evaluate clustering performance using marker genes and modify parameters of the pipeline accordingly, as best practices guides and clustering tutorials explicitly prescribe^50,51^. So, it is not that the standard pipeline is not useful; it just must be carefully managed in order to generate results that can be used to generate biological insights or suggest hypotheses that can be later tested. Waddington’s landscape, however, does not state that there should be some effective procedure for using epigenetic data to generate cell type clusters in a guided clustering process. Rather, it puts forward a mechanistic explanation for how differences in epigenetic state arise and are expressed phenotypically, and this explanation relies on attractors in the corresponding epigenetic space.

Of course, the fact that the current pipeline does not reliably invert the sampling process to re-generate the original Waddington-like groups does not mean that sampling noise is not ultimately the cause for the lack of distinct cell-types we see here. All we can currently say is that the majority of datasets we analyzed do not conform to the Waddington picture. Further improvements in experimental platforms, particularly the development of more reliable measurement techniques, will be needed to completely rule out this “sampling noise” hypothesis. This issue with noise, and experimental improvements that could help in this regard, are discussed in more detail in the “Limitations of the current study” section below. If the observations made here are *not* due to widespread inaccuracy or bias in experimental techniques, then our findings would suggest that Waddington’s landscape does not hold for mRNA levels within cells. Interestingly, the landscape originally conceptualized by Waddington made reference to an abstract epigenetic space quite different from the modern interpretation, in part because the proposal itself was made long before the advent of modern molecular biology^28,29^. One hypothesis that emerges from our work is that Waddington’s explanation for canalization is fundamentally correct, but it is just that the epigenetic space is not a space of merely mRNA levels. While we have considered protein levels and patterns of chromatin accessibility in this work, it is possible that further advances in the technologies that generate these data could reveal a more Waddington-like pattern. In particular, advances in tools that allow single-cell characterization of protein levels, especially at the genome-wide level, could eventually allow us to determine if the patterns we observe here hold for a larger set of proteins and across a wider range of systems^101–103^. In this scenario, since mRNA molecules form the instruction set for the production of proteins, it is not clear how a regulatory system would achieve a stable set of distinct attractors in protein levels based on an exceedingly heterogeneous distribution of mRNA levels. It is also unclear why a system would evolve to have no attractors in mRNA space, but then evolve a regulatory system that generates such attractors in protein space.

An alternative possibility could be that Waddington’s landscape involves a completely different epigenetic state of the cell. In this scenario, cell types would correspond to attractors in this (heretofore undescribed) epigenetic landscape, and there is some non-linear projection from those attractors into the spaces that current single-cell technologies allow us to access experimentally (Fig. 7B). If this hypothesis holds true, it may be possible to find a more principled approach to inverting the projection that performs more reliably than the set of transformations that form the current standard of practice in the field^37,50^.

It is possible, however, that canalization in development takes a very different form from that originally envisioned by Waddington. One hypothesis would be that, rather than representing stable attractors in epigenetic space, cell types could be specified as a set of constraints on the *dynamic responses* of cells. In this view, instead of corresponding to different point attractors on an epigenetic landscape with corresponding basins of attraction, cell types would instead correspond to sets of “allowable dynamics” in response to perturbations. So, instead of point attractors, cell types would correspond to sets of “paths” or “trajectories” in state space. Indeed, a large amount of previous work has demonstrated that the dynamics of cellular response can extremely biologically informative^104,106,108,110,112^. That being said, a great deal of work will need to be done in order to fully establish a theoretical framework based on constrained dynamics that is as well-developed as the Waddington attractor theory^32,114,116^.

Another alternative is that cell state is far more complex and continuous than the framework of discrete cell types can admit. For instance, single-cell measurements of physiological responses of cells ranging from breast tissue to the immune system reveal incredible diversity in those responses^34–36,98^. In other words, the biological responses of individual cells that correspond to our classical notion of a “cell type” is itself quite heterogeneous, suggesting that categorizing cells into discrete groups in the first place may mask critical aspects of their physiology. Emerging functional data at the single-cell level thus suggest that the “final step” in development may not be a discrete set of cell types, as Waddington’s landscape posits (Fig. 1A), but rather a more continuous spectrum of states and phenotypes that is not well-approximated by the attractor picture. Regardless of whether or not there are cell-type attractors to be found in some as of yet uncharacterized epigenetic space, it is clear that careful analysis of available data, and a willingness to test even well-established paradigms against that data, is critical to the future of single-cell biology.

### Limitations of the current study

Our analysis covered a wide range of techniques for measuring epigenetic state at the single-cell level, encompassing multiple sequencing-based tools like 10X, ParseBio and BD Rhapsody, microscopy-based techniques like MERFISH, measurements of chromatin accessibility and protein levels, among many others^73,75,76,84,93,94,105,107,109,111^. The fact that we see essentially identical results across these divergent experimental platforms makes it less likely that those results represent purely technical artifacts. It is nonetheless still possible that all of these methods lack the accuracy needed to characterize cell-type attractors at the single-cell level. If that is the case, this would suggest a critical need to develop new experimental techniques that can accurately characterize the epigenetic states of cells. In particular, improving effective capture rates (i.e., ensuring that more of the transcripts that are actually present in cells are reflected in the data) would help reduce the potential impact of sampling noise on the types of analyses presented here. In addition, current estimates of the capture probability vary widely and are generally based either on cDNA spike-ins (which may have different properties from endogenous mRNA) or on ballpark calculations based on rough ideas of the number of mRNA molecules that should be present in cells. More experimental effort targeted at understanding the precision and accuracy of single-cell techniques would be extremely helpful in establishing whether the lack of separation observed here is due to technical limitations of the techniques themselves.

Another limitation of this work is the fact that, as mentioned above, current data on protein levels in single cells is much less extensive than that available for mRNA levels, both in terms of the numbers of macromolecules measured in each study (∼100-1000 compared to genome-wide) and the number of tissues and organisms that have been considered. Further advancements in single-cell measurement technologies for proteins, and for other epigenetic characteristics of cells, will be necessary to fully test the predictions of Waddington’s landscape in those spaces.

A final limitation we will mention is the fact that we have very few datasets where we know the cell type identity of each cell *a priori*, which makes it difficult to directly compare epigenetic states for groups of cells that we would classify as the same “type” based on well-validated orthogonal markers. Most of the examples of orthogonally-validated cell types that we have are from cell lines. Of course, cells that have been immortalized and transformed to grow well in an *in vitro* setting may exhibit many changes in epigenetic characteristics from their native counterparts. The primary example of data for known cell types is from a 2017 study by Zheng *et al.* on FACS-separated PBMCs; indeed, this is one of the few “gold standard” datasets and is still used in many benchmarking studies of clustering algorithms^40,119,120^. This study is now almost 10 years old, however, and focuses on just a single tissue type. More examples of this kind of data, generated with more modern technologies, will be critical to testing any theory on the epigenetic basis for cell type specification and maintenance. These examples will also be of great help in training AI foundation models to predict cell types based on epigenetic data^121–123^.

## Methods

### Datasets

The vast majority of data we analyzed was taken directly from freely-available repositories on the internet. Supplementary Table S1 summarizes each of these datasets, including the name we have given to each data set, number of cells, number of features measured, the paper citation associated with the dataset and where we obtained the data. CSV files for all of the datasets we used, specific to each transformation and analysis, are also provided for each dataset as additional supplementary material through Github (https://github.com/DeedsLab/Epsilon-Network).

The MERFISH data that we analyzed is the only data that is not publicly accessible in an appropriate format online. The MERFISH data for the mouse Traumatic Brain Injury dataset was obtained using the protocols and analysis described in ref.^76^. The count-by-cell matrix for this dataset was kindly provided by Zach Hemminger and Roy Wollman. Similarly, the Vizgen corporation has made image data from their MERFISH experiments on mouse brain slices freely available on the web^75^. Data for one of these slices was obtained through image analysis as described in, and again kindly provided by Zach Hemminger and Roy Wollman. All the MERFISH count matrices are provided as additional supplementary information.

### Pre-processing and transformations

For any scRNA-seq dataset we considered, we either performed our analysis on post-QC data provided by the original authors, or performed standard quality control filtering based on the standard practice in the field^50,51^. For all data used in this study, we inspected the joint distributions of three quality control metrics: 1) the number of genes per cell, 2) the UMI count per cell, and, if available, 3) the fraction of mitochondrial gene counts per cell. We used scatter plots with histograms to visualize the joint and individual distributions of these QC covariates, and to manually determine a threshold for removing outliers. For example, in a typical QC procedure aiming to remove low quality or dying cells, we use relatively permissive thresholds to remove cells with a relatively low number of genes identified per cell, and a relatively low number of UMI counts per cell, but a high proportion of mitochondrial genes. To filter out potential doublets, we visually inspected the joint distributions of the number of genes per cell and the UMI counts per cell. If we identified a small subset of cells that appeared to exist multiple standard deviations away from the main mass of the joint distribution, then these outliers were removed.

We applied an array of normalization and transformation approaches: namely counts per million normalization (CPM), CPM and log+1 transformation, CPM-log+1-PCA, PCA on the raw data itself, Zscore normalization, Z-score then CPM, CPM then Z-score, cell-cycle regression, and Sanity normalization^113^. For the “Standard scRNA-seq analysis approach” we first CPM normalize the data using a scale factor of 10,000. Then we apply a log+1 transformation, followed by sub setting the genes into feature sets^50^. The standard analysis pipeline uses the Highly Variable Gene feature selection model, which we applied using the Scanpy software with the ‘highly_variable_genes’ function, with n_top_genes = 2000, and flavor = ‘seurate_v3’^50,51^. In addition to the HVG feature selection, we also used the Differentially Distributed Gene feature selection model with the capture probability parameter, p_cap set to 5%^57^. When orthogonal cell type labels were available, we used these labels to generate a supervised set of Wilcoxon Rank Sum feature genes. Finally, we also used references from the literature to define marker genes to use as feature genes for the 10x v3 PBMC data^115^. After feature selection, we apply PCA, and select the number of components by applying the “elbow method” to detect the inflection point in which including additional components no longer explains additional variance in the data.

In the absence of one particular standardized method for analyzing scATAC-seq data, we relied on how the available data from previous studies had been processed by the original authors. Details on how scATAC-seq data had been QC filtered and pre-processed to generate raw chromatin availability scores, can be found in the method sections of the original studies [see Supplementary Table S1]. For a subset of the scATAC data, we performed our analysis on both the raw chromatin availability score, as well as data that had been ‘feature selected’ using Latent Semantic Indexing by the original authors of the study. Whether we analyzed the raw or the Latent Semantic Indexing data is indicated in each sub panel of our figures, and in Table 1.

### Simulated and down-sampled data

To test the sensitivity of our *ϵ* networks we generated several synthetic datasets where we could sample “cells” from distinct distributions. The case we focused on in the Main Text (Fig. 3) was a Negative Binomial Mixture Model in which we sampled 1,000 cells from a distribution for “cell type A” and 1,000 cells from a different distribution for “cell type B.” We chose the Negative Binomial distribution as it has been suggested as a reasonable model for the types of count distributions observed in scRNA-seq data^67,68^. To generate realistic parameters for this experiment, we analyzed the A20 & NIH3T3 10x scRNA-seq data. In this dataset, we first identified all the “differentially expressed” genes between the two cell types using a standard Wilcoxon rank-sum test (note this was based on the raw UMI count data and not any transformed space). This generated a list of genes with statistically significant differences between the two groups. We then ranked them by their average expression in the highly expressing cell type, reasoning that more highly expressed genes would be more helpful for separating groups. We found that typical gene expression levels for marker genes had around 8 counts in the highly expressing cell type and around 1 in the lowly-expressing cell type. We found around 250 genes or so were markers for each cell type based on this rough criterion (see Supporting Data Tables S2 and S3).

Each cell in this simulation had a total of 10,000 genes, approximating the dimensionality of the A20/NIH3T3 data, and as mentioned above we sampled 2,000 cells total, 1,000 from each cell type.

Based on the above analysis, we chose to have 250 genes as “markers” for each cell type, leading to a total of 500 marker genes (i.e. 250 marker genes for cell type A and 250 for cell type B) with the expression levels described above. The remaining 9,500 genes were the same for all cells, and were given the “low” expression state to model the fact that many genes in the data are not expressed at a high level. So, only 5% of the genes in this dataset are markers, and the rest are essentially “noise.”

In the first 250 features, cells in for type A were drawn from a Negative Binomial (NB) distribution with a probability of success was set at 0.5. In this case, we can set the average expression of the distribution simply by changing the parameter for the number of successes in the distribution. We chose values of 10, 8, 6, …, 2, 1 for this parameter. This means that we modulated the average number of counts in the distribution for the marker genes across that range, which is similar to the range of real average counts observed in the A20/NIH3T3 data (Supporting Data Table S2). For this first 250 features, expression values for type B cells were drawn from a NB distribution with a success probability of 0.5 and a success number of 1, corresponding to a gene with mean expression of 1. In the next 250 features, cells for cell Type B had the higher level of average expression (10, 8, 6, …, 2, 1) and cells for cell type A had the low expression (success probability 0.5, number of successes of 1). For the remaining 9,500 features, values for cell types A and B were drawn from the *same* distribution with a probability of success of 0.5 and a number of successes of 1 (modeling the “low” expression state). Results from this model are shown in Figs. 3 and S3.2.

We generated a similar synthetic dataset using a Poisson, rather than NB, distribution to simulate the counts. Here again, we had 250 features as “marker” genes for cell type A, with parameters chosen to have average expression levels of 10, 8, 6, …, 2, 1 depending on the simulation. Similar values were used for the next 250 features for cell type B. In each case, an average expression level of 1 was used for the corresponding “other” cell type, and for the 9,500 genes that were not used as markers. The results for this model may be found in Fig. S3.2. We also considered several simple Gaussian Mixture Models with varying structures and parameters and found similar results (Fig. S3.1).

In addition to this purely simulated data, we also simulated what might happen if we take a real dataset where we observe separation and increase sparsity by down-sampling that data. To do this, we focused on the A20/NIH3T3 scRNA-seq data mentioned above. For each simulation, we first set a down-sampling probability *p_D_*; this is the probability that any given count in any given cell would be retained. To down-sample the data, we went through every cell *i* and every gene *j*; call the UMI counts for that gene in that cell *n_i_*_,j_. We then sampled a *new* value, *n*\**_i_*_,j_ from a Binomial distribution with *n_i_*_,j_ trials and a proibability of success *p_D_*. The results for *p_D_* = 0.5, 0.2, 0.1, 0.05 are shown in Fig. S3.3

### ∈ network construction, analysis and visualization

For every dataset and every transformation, the data consists of a matrix where the rows correspond to each individual cell and the columns to the features measured in the single-cell experiment (note that the data is sometimes represented with the columns as cells and the rows as features, depending on the specific study in question). For raw scRNA-seq data, these features are all the genes in the genome, and the entries in the matrix are the number of UMI counts for that gene in that cell. All of the other data considered here (MERFISH, BD Rhapsody mRNA and protein, etc.) ultimately consists of a similar matrix, just with either fewer genes measured (in the case of MERFISH, for example) or a different type of measurement for each entry (for instance, protein levels rather than UMI counts in the BD Rhapsody protein data). Transformations of the data result in similar matrices, but in those cases the entries in the matrix correspond to different quantities (CPM normalized data, log CPM+1, etc.). Application of dimensionality-reduction tools like PCA of course result in fewer features in the resulting matrix.

For any given dataset, we can easily calculate the distance between any two cells using their corresponding feature vectors. The figures in the main text focused on the simple Euclidean distance (i.e. the *ℓ*^2^ norm) to define the distance, since this is definition of distance used in the vast majority of scRNA-seq and single-cell genomics studies. We considered other types of metrics, including the *ℓ*^1^ norm and Hamming distance, as described in the text and supplementary information (Supp. text Section 2 and Figures S5.16, S5.17, S6.11 and S6.1). Calculating the distance between every pair of cells results in a symmetric distance matrix with a number of elements equal to the number of cells in the dataset squared.

Once we calculate this distance matrix, we can use that matrix to generate an epsilon network. To do so, we first define the cutoff ɛ to be some value. Then, for each cell *i* in the dataset, we go through the row in the distance matrix and consider every other cell *j*. If the distance between these cells is less than the cutoff (*d_i_*_,j_ < ɛ), then we add cell *j* to the list of cells that are connected (or adjacent) to cell *i*. Doing this for every cell in the dataset generates a standard adjacency list representation of the *∈* network.

We analyzed the resulting *ϵ* network using a standard set of algorithms on graphs. For instance, we used a Depth-First-Search (DFS) to determine all of the components in each of our *ϵ* networks. The largest such component is the giant component, and the number of cells in that largest component is the size of the giant component. Similarly, the “degree” of any cell *i* for a given value of *ϵ* is just the size of the adjacency list for that node in the graph^91^. Degree distributions were calculated as a histogram across these individual degrees. Note that the histograms in Fig. 3 and in the Supplementary Information use a standard logarithmic binning approach for approximately scale-free distributions.

Distance matrices were calculated either using the scanpy package in Python^51^ or using custom-built C++ software (particularly for larger datasets). Analysis of the giant component size and composition as a function of *ϵ* was performed using custom-built C++ software. Degree distributions and force-directed layouts were calculated using the NetworkX package in Python, and all plots were generated using the matplotlib package in Python^117,118^. All software used in this work is available on Github at the following link: https://github.com/DeedsLab/Epsilon-Network.

## Acknowledgements

The authors thank Tom Kolokotrones, Walter Fontana, Alexander Hoffmann, Roy Wollman, Jukka Keranen, Pavak Shah, Zach Hemminger, Ivy Xiong, Gunalan Natesan, Nina Gilshteyn, Hannah Tamura, and other members of the Deeds lab for many helpful discussions and comments. We also thank Zach Hemminger and Roy Wollman with their assistance in analyzing MERFISH data. This work was supported by an NIH IRACDA postdoctoral fellowship 2K12GM106996-06 to BS and NIH R01-GM143378 to EJD.

