## Supplemental Figures for "A lack of distinct cell identities in single-cell measurements: revisiting Waddington’s landscape"

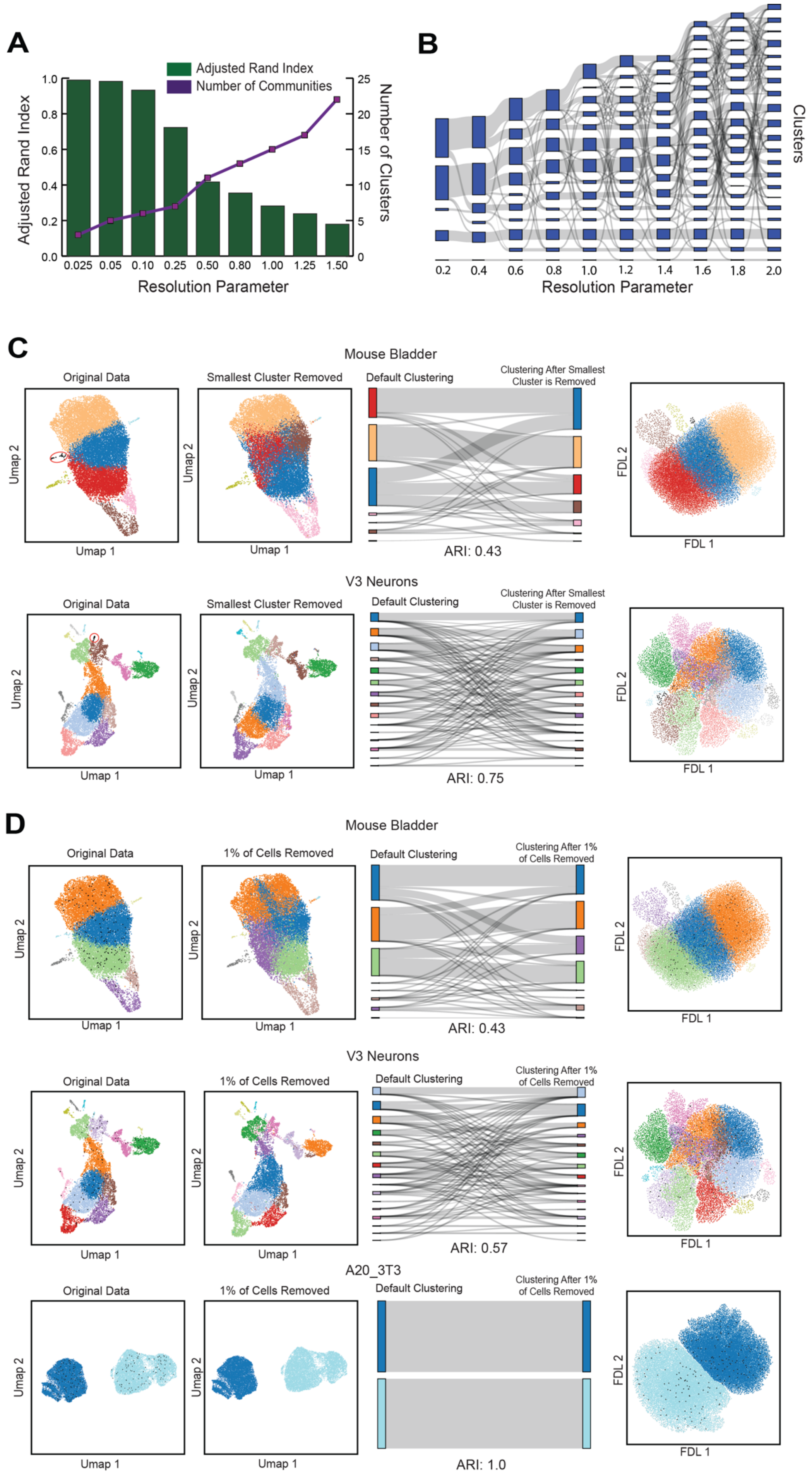
A
Adjusted Rand Index
Number of Communities
Adjusted Rand Index
Number of Clusters
Resolution Parameter
0.025 0.05 0.10 0.25 0.50 0.80 1.00 1.25 1.50
B
Clusters
Resolution Parameter
0.2 0.4 0.6 0.8 1.0 1.2 1.4 1.6 1.8 2.0
C
Mouse Bladder
Original Data
Smallest Cluster Removed
Default Clustering
Clustering After Smallest Cluster is Removed
Umap 1
Umap 2
FDL 1
FDL 2
ARI: 0.43
V3 Neurons
Original Data
Smallest Cluster Removed
Default Clustering
Clustering After Smallest Cluster is Removed
ARI: 0.75
D
Mouse Bladder
Original Data
1% of Cells Removed
Default Clustering
Clustering After 1% of Cells Removed
ARI: 0.43
V3 Neurons
Original Data
1% of Cells Removed
Default Clustering
Clustering After 1% of Cells Removed
ARI: 0.57
A20_3T3
Original Data
1% of Cells Removed
Default Clustering
Clustering After 1% of Cells Removed
ARI: 1.0

**Figure S1.** Identification of meaningful cell types by community detection algorithms is parameter dependent and extremely unstable. **A)** Adjusted Rand Index (ARI) and Number of Communities as a function of the resolution parameter in Louvain clustering implemented in Seurat. Louvain clustering was performed on the filtered, raw UMI counts of10x genomics scRNA-seq data from three FACS-purified lymphocyte types: B cells, Monocytes and Natural Killer cells. The ARI was computed by comparing the identity of cells within the Louvain-determined clusters to the ground truth cell type identities determined by FACS. As shown, there is no natural minimum by which to determine a reasonable clustering parameter to use, even when orthogonal labels can be used to validate clustering. **B)** River plot of clustering obtained by the applying the Louvain clustering algorithm after the standard pipeline (implemented in scanpy) to the same group of cells in part **A.** with varying resolution values. Rather than showing a clear hierarchical structure where increasing resolution allows for larger clusters to split cleanly into multiple clusters, increasing resolution results in cells from separate clusters merging to form new clusters. **C)** UMAPs of both the Mouse Bladder and V3 Neurons where the data is clustered via Leiden as part of the standard analysis pipeline, with the same number of clusters found as the original authors. The left most panels show the clustering based on the standard pipeline, while the panels in the second column from the left show the clustering after the smallest cluster found when clustering the data originally is removed (around 35 cells total in both cases). The cells removed are colored black and circled the leftmost panel. If cells exist in well-defined basins, removal of one small group should minimally affect the other clusters. Instead, as shown on the Sankey Diagram to the right, the removal of the smallest cluster results in pairs of cells that were previously in the same cluster to split while pairs of cells in different clusters are then joined together in the same cluster. This results in a remarkably low ARI value. On the far right, we show a Force Directed Layout (FDL) of the k-Nearest Neighbor graph used as the input to Leiden clustering. The completely connected and highly overlapping nature of this graph likely contributes to the instability in clustering. **D)** UMAPs of both the Mouse Bladder, V3 Neurons and NIH 3T3/A20 Cell Line data, where the data is clustered via Leiden as part of the standard analysis pipeline, with the same number of clusters found as the original authors. The left most panels show the clustering based on the standard pipeline, while panels in the second columns from the right show the clusterings after 1% of cells are *randomly* removed. The cells that were removed are colored in black in the leftmost panels. In the Mouse Bladder and V3 Neurons data, removal of these cells results in dramatically different clustering results, as highlighted in the Sankey diagrams in the middle. In contrast, however, removal of random cells in the Cell Line data has no effect on the clustering results. As we can see in the FDL results on the right, the Cell Line data actually forms two discrete groups that just barely touch on their edges, and so removing random cells does not influence the final result. Comparison of the FDL for this case with those from the Mouse Bladder and V3 Neuron datasets suggests that the structure of the data has a strong influence on the statistical robustness of clustering results.

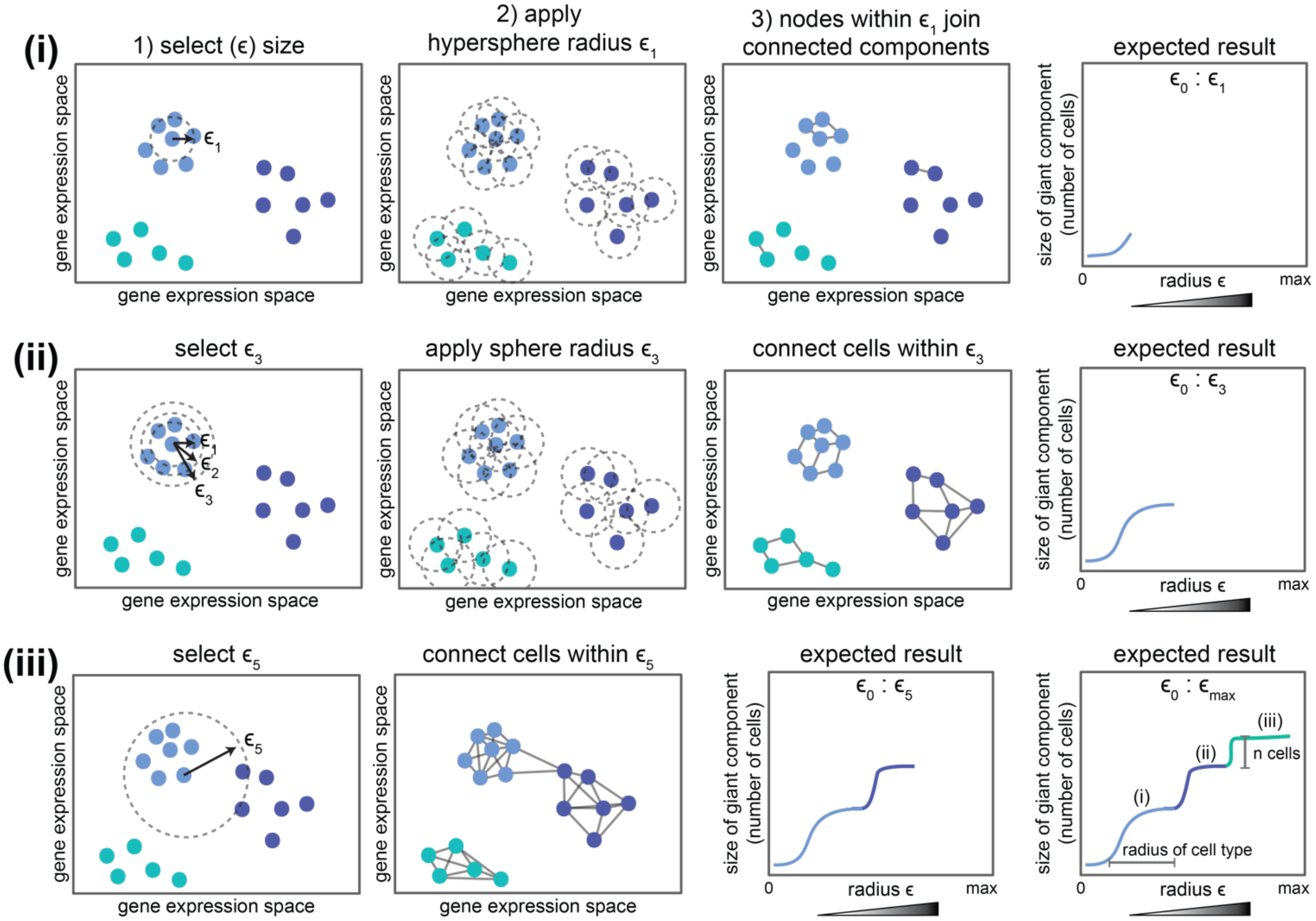


**Figure S2.1** Quantifying the size of the giant component in "$\epsilon$ network" analysis. **(i-iii)** Schematic illustrating the steps in the "$\epsilon$ network" analysis. For any single-cell dataset we can calculate the distance between any two cells using their feature vectors (i.e. gene expression values). To perform the $\epsilon$ network analysis, we **(i)** first select an n-dimensional hypersphere of radius $\epsilon$, where each dimension represents a gene (1). We then apply this hypersphere to each cell in the data (2). If two cells are closer to each other than this radius $\epsilon$, i.e. they lie within each other's hypersphere, then those two cells are connected in the network (3). For any given value of $\epsilon$, cells will partition into groups of cells, or "components", that are connected to each other (3). The component containing the highest number of cells is known as the giant component (3). In many of our $\epsilon$ network analyses, we plot the size of the giant component as we increase the size of radius $\epsilon$ (rightmost panel). **(ii)** If cells are distributed in gene expression space according to the Waddington paradigm, increasing values of $\epsilon$ will at first lead to an increase in the size of the giant component, as the densest group of cells becomes fully connected in the network. As $\epsilon$ continues to increase, we expect the size of the giant component to plateau, as no additional cells are nearby to be added to the largest cluster. **(iii)** After a certain threshold of $\epsilon$, other groups of cells are expected to join the giant component all at once, giving rise to a characteristic step-like behavior in the giant component.

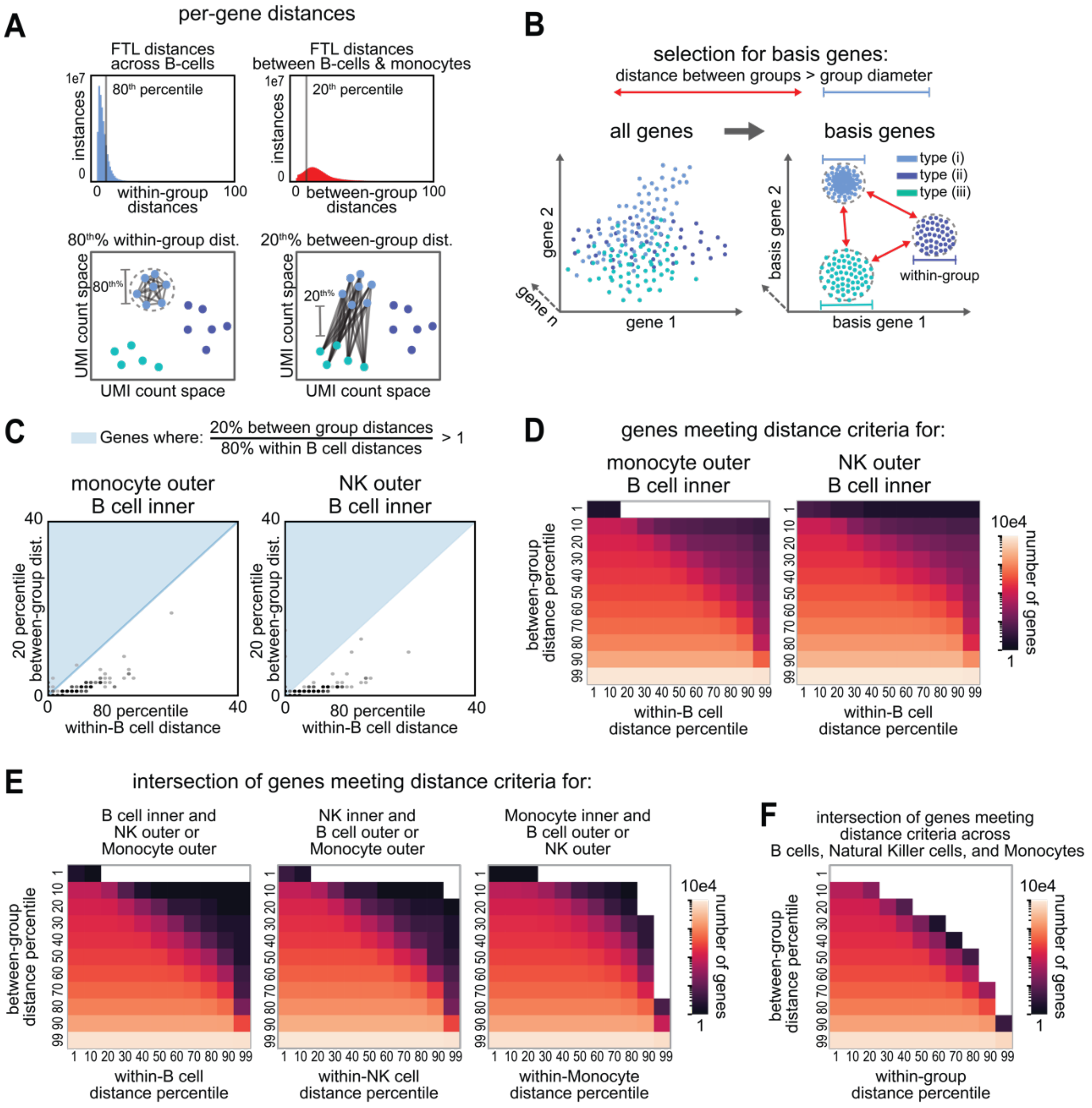
A
per-gene distances
FTL distances across B-cells
1e7
instances
80th percentile
within-group distances
0
100
FTL distances between B-cells & monocytes
20th percentile
between-group distances
80th% within-group dist.
20th% between-group dist.
UMI count space
B
selection for basis genes:
distance between groups > group diameter
all genes
basis genes
gene 1
gene 2
gene n
basis gene 1
basis gene 2
type (i)
type (ii)
type (iii)
within-group
C
Genes where: 20% between group distances / 80% within B cell distances > 1
monocyte outer B cell inner
NK outer B cell inner
20 percentile between-group dist.
80 percentile within-B cell distance
40
D
genes meeting distance criteria for:
monocyte outer B cell inner
NK outer B cell inner
between-group distance percentile
within-B cell distance percentile
1 10 20 30 40 50 60 70 80 90 99
10e4
number of genes
1
E
intersection of genes meeting distance criteria for:
B cell inner and NK outer or Monocyte outer
NK inner and B cell outer or Monocyte outer
Monocyte inner and B cell outer or NK outer
between-group distance percentile
within-B cell distance percentile
within-NK cell distance percentile
within-Monocyte distance percentile
10e4
number of genes
F
intersection of genes meeting distance criteria across B cells, Natural Killer cells, and Monocytes
within-group distance percentile

**Figure S2.2.** A distance-based, supervised feature selection method does not identify a set of genes that can reasonably separate B-cells, monocytes, and Natural Killer cells. **A)** Schematic of distance criteria for gene selection. Percentiles for *within-group* distances were defined on a per-gene basis by measuring, for each individual gene, the Euclidean distance between each cell *of the same type*. Percentiles for *between-group* distances were defined on a per-gene basis by measuring, for each individual gene, the Euclidean distance between each cell of two *different cell-type groups.* For example, in A), two histograms display the distance percentiles for the gene "FTL". On the left, (in blue), is the distribution for how far apart each B cell is from other B cells, based on their expression of FTL (i.e. the *within-group* distance). The grey vertical line marks the 80$^{th}$ percentile distance, a conservative approximation for the "diameter" of FTL expression across B cells. The histogram on the right displays the distribution, (in red), of the difference in FTL expression between all of the B cells and monocytes (i.e. the *between-group* distance). Here the grey vertical line marks the 20$^{th}$ percentile distance, a conservative approximation for the minimum distance of FTL between two cell types. **B)** Schematic illustrating the rationale for a distance-based feature selection approach. Basis genes are defined as those genes where the differences in gene expression (or Euclidean distance) *between* two cell type groups is greater than the differences in gene expression *within* a single cell type group. This set of genes are expected to produce cell type-groups that are separable in gene expression space. **C)** Scatter plot illustrating the set of genes for which the 20$^{th}$ % between-group distance is larger than the 80$^{th}$ % within-B cell distance. Genes that meet this criteria lie in the region shaded in blue. For both panels, B cells are used to define the within-cell type distance. The left panel displays gene distances between B cells and monocytes while the right panel displays gene distances between B cells and Natural Killer cells. **D)** Heat maps illustrating the number of genes where the *between-group* distance for a given percentile (on the y-axis) is bigger than the *within-group* distance for a given percentile (on the x-axis). For both panels, B cells are used to define the *within-group* distance. The left panel displays gene distances between B cells and monocytes while the right panel displays gene distances between B cells and Natural Killer (NK) cells. **E)** Heat maps quantifying the intersection of genes across two pairs of cell types, where a gene meets the indicated distance criteria (*between-group* % > *within-group* %) for each pair of inner and outer cell type. For example, in the first panel, B cells are used to measure *within-group* distances, and either the distance between B-cells and NK cells, or B-cells and Monocytes are used to define the *between-group* distances. The number of intersecting genes is the count of those genes where the distance criteria is met across both groups. **F)** Heat map illustrating the number of genes meeting the distance criteria across all six pairs of cells for the three cell-types. For example, there is just one gene where the 40$^{th}$ percentile distance between cells of different cell types is larger the 70$^{th}$ percentile distance within a given cell type group.

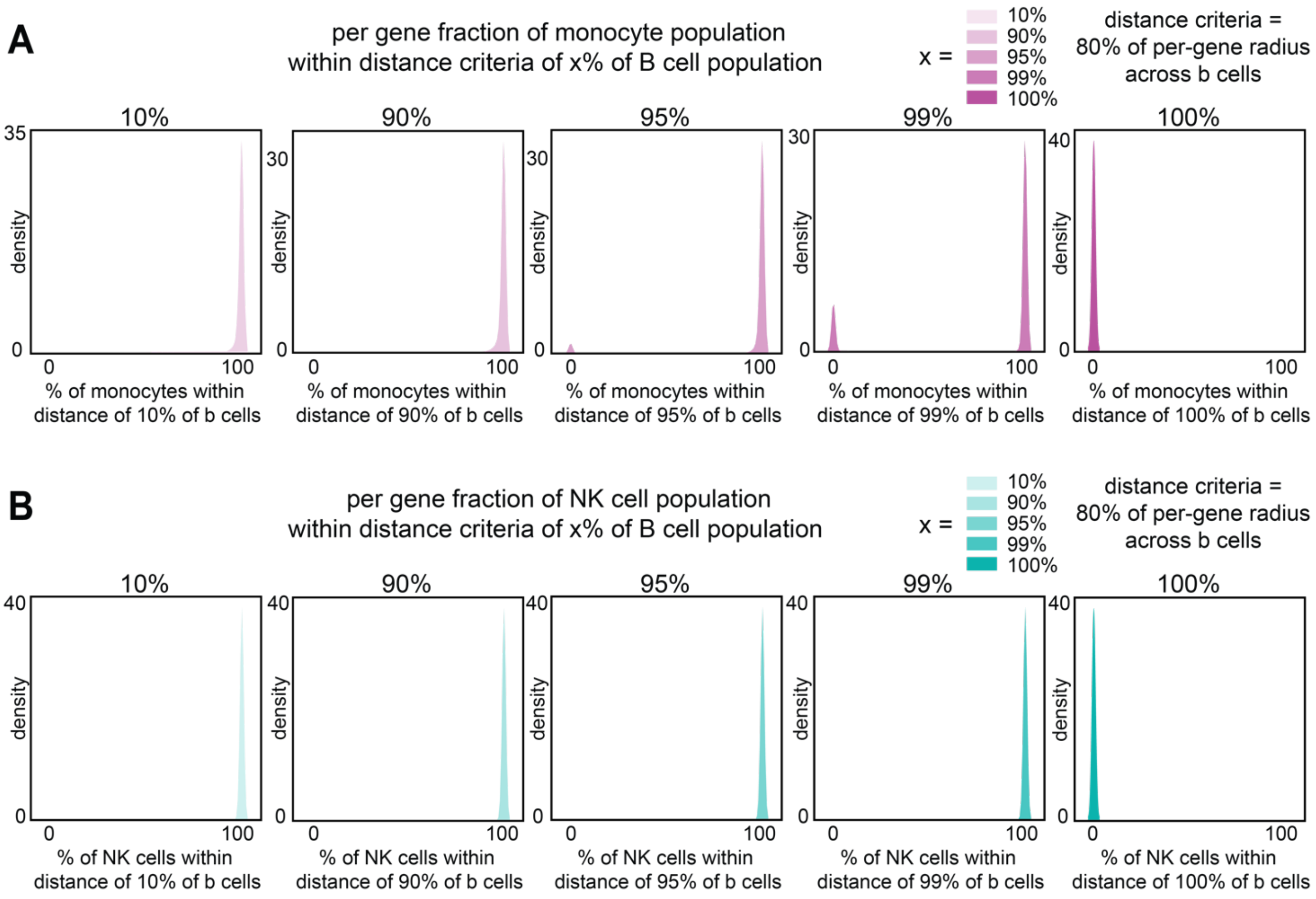


**Figure S2.3.** Quantifying the fraction of cells from different cell types in proximity to B cells using a per-gene distance threshold. **A)** Density distributions for the percent of monocytes that are approximately as close to a given fraction of B cells, as other B cells are to B cells. For example, the first panel illustrates that ~100% of monocytes are as close to at least 10% of B cells, as other B cells are to B cells, for each gene measured. *For each individual gene*, the diameter across B cells is approximated as the 80$^{th}$ percentile distance in the distribution of distances between B cells. The density distributions represent the fraction of the monocyte population that is in relatively close proximity to B cells across each *individual gene* in the data. For example, the fourth panel illustrates that for each gene, the majority of monocytes are as close to *99% of B cells,* as other B cells are to B cells. However, there is a small fraction of genes where monocytes are only as close to 95% of B cells as other B cells, but not 99%. **B)** Density distributions for the percent of NK cells that are approximately as close to a given fraction of B cells as other B cells are to B cells (approximated using the 80$^{th}$ % within B cell distance for each individual gene). While there are zero genes that place any NK cells within the same distance to 100% of B cells, as other B cells are to B cells, *all* of the genes measured place NK cells within the same distance to 99% of B cells, as other B cells are to B cells.

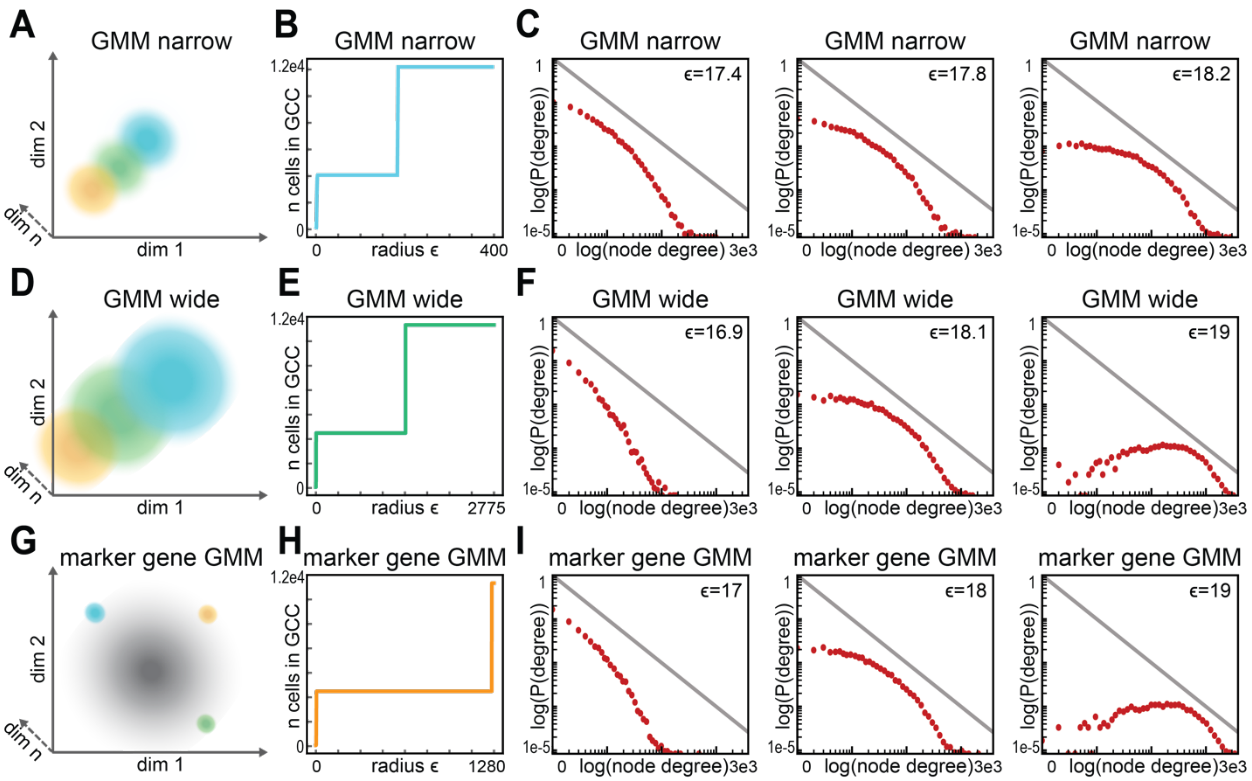


**Figure S3.1.** $\epsilon$ network analysis can identify distinct gaussian distributions in gaussian mixture models with overlap and noise. **A)** Schematic illustrating the “narrow” gaussian mixture model (GMM) analyzed in panels B and C. “GMM narrow” is a mixture of three different 200-dimensional gaussian distributions with identity covariance. The mean of the first gaussian is 100, the mean of the second gaussian is 114, and the mean of the third gaussian is 128. 4000 points were sampled from each. **B)** Plot of the size of the giant component over increasing radius $\epsilon$ for the $\epsilon$ networks constructed using the narrow GMM described in (A). Note that, since all three Gaussian clusters are equidistant from one another, two of the clusters join at once, generating a single large jump in this and all the other models considered here. **C)** Degree distributions at the indicated $\epsilon$ for the $\epsilon$ networks of the narrow GMM described in (A). **D)** Schematic illustrating the “wide” gaussian mixture model (GMM) analyzed in panels E and F. “GMM wide” is a mixture of three different 200-dimensional gaussian distributions with identity covariance. The mean of the first gaussian is 100, the mean of the second gaussian is 200, and the mean of the third gaussian is 300. 4000 points were sampled from each. **E)** Plot of the size of the giant component over increasing radius $\epsilon$ for the $\epsilon$ networks constructed using the wide GMM described in (D). **F)** Degree distributions at the indicated $\epsilon$ for the $\epsilon$ networks of the wide GMM described in (D). **G)** Schematic illustrating the “marker gene” gaussian mixture model (GMM) analyzed in panels H and I. “Marker gene GMM” is a mixture of three different 200-dimensional gaussian distributions with identity covariance. In the first gaussian, the mean of the first component is 1000, while the mean of the other components is 100. In the second gaussian, the mean of the second component is 1000, while the mean of the other components is 100. In the third gaussian, the mean of the third component is 1000, while the mean of the other components is 100. The center of each gaussian is equidistant from the others, and 4000 points were sampled from each. **H)** Plot of the size of the giant component over increasing radius $\epsilon$, for the $\epsilon$ networks constructed using the marker gene GMM described in (G). **I)** Degree distributions at the indicated $\epsilon$ for the $\epsilon$ networks of the marker gene GMM described in (G).

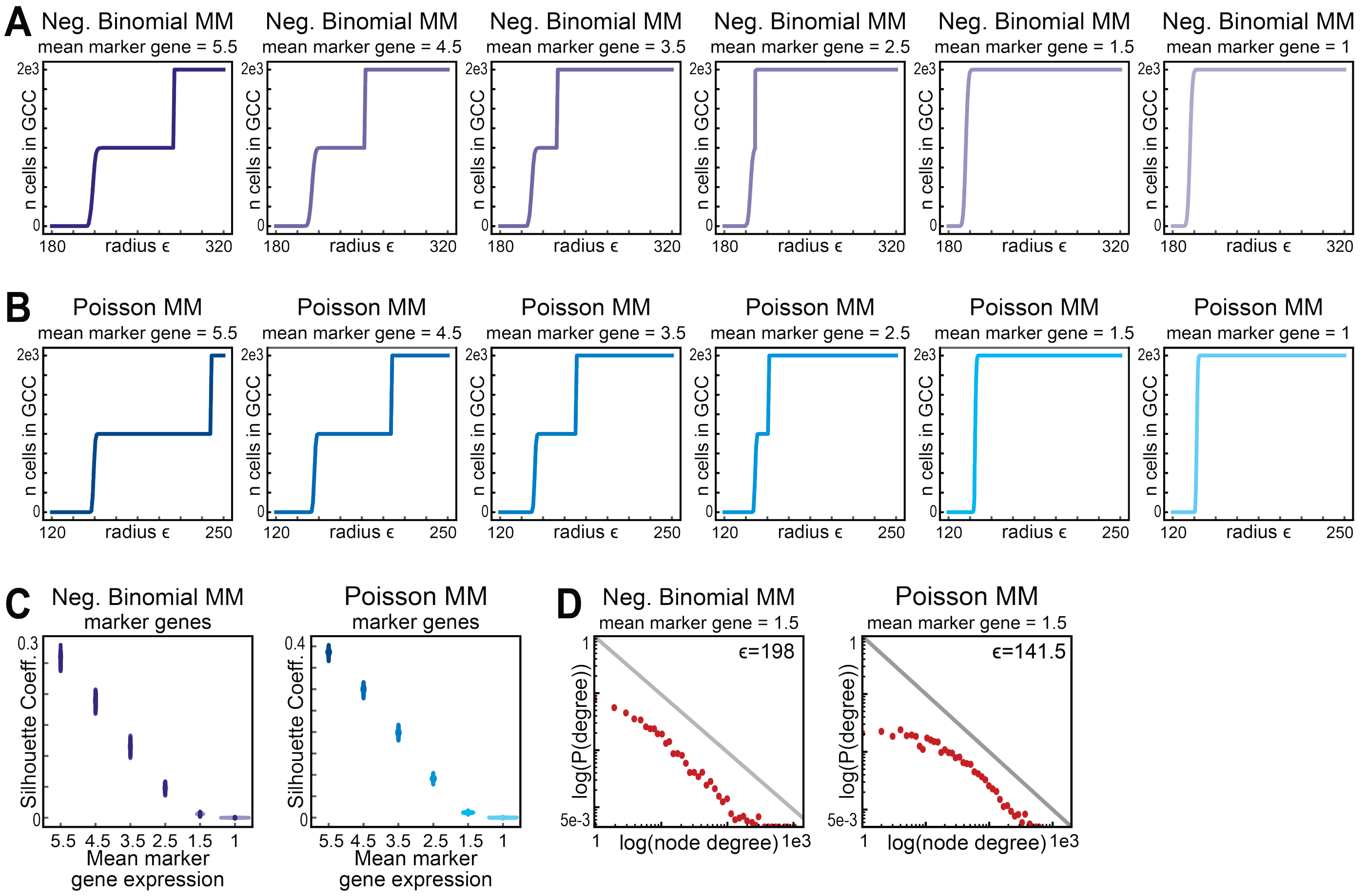


**Figure S3.2.** $\epsilon$ network analysis can identify distinct distributions in mixture models with noise and low marker gene expression. Briefly, each synthetic dataset modeled a mixture of two distributions. In each distribution, 1000 points were sampled from a 10,000 dimensional feature space. For each group of 1,000 points, only 250 of the 10,000 features varied in magnitude, in order to model a situation where only a fraction of marker genes demonstrate differential expression levels. **A)** Plot of the size of the giant component over increasing radius $\epsilon$ for $\epsilon$ networks constructed using a Negative Binomial Mixture model with varying levels of marker gene expression. Model construction details are provided in the Methods section description of Synthetic Negative Binomial Data. **B)** Plot of the size of the giant component over increasing radius $\epsilon$ for $\epsilon$ networks constructed using a Poisson Mixture model with varying levels of marker gene expression. Model construction details are provided in the Methods section description of Synthetic Poisson Data. **C)** Silhouette coefficient for the Negative Binomial and Poisson Mixture models described in (A) and (B). **D)** Density distributions for the marker gene models described in (A) and (B) at the indicated $\epsilon$.

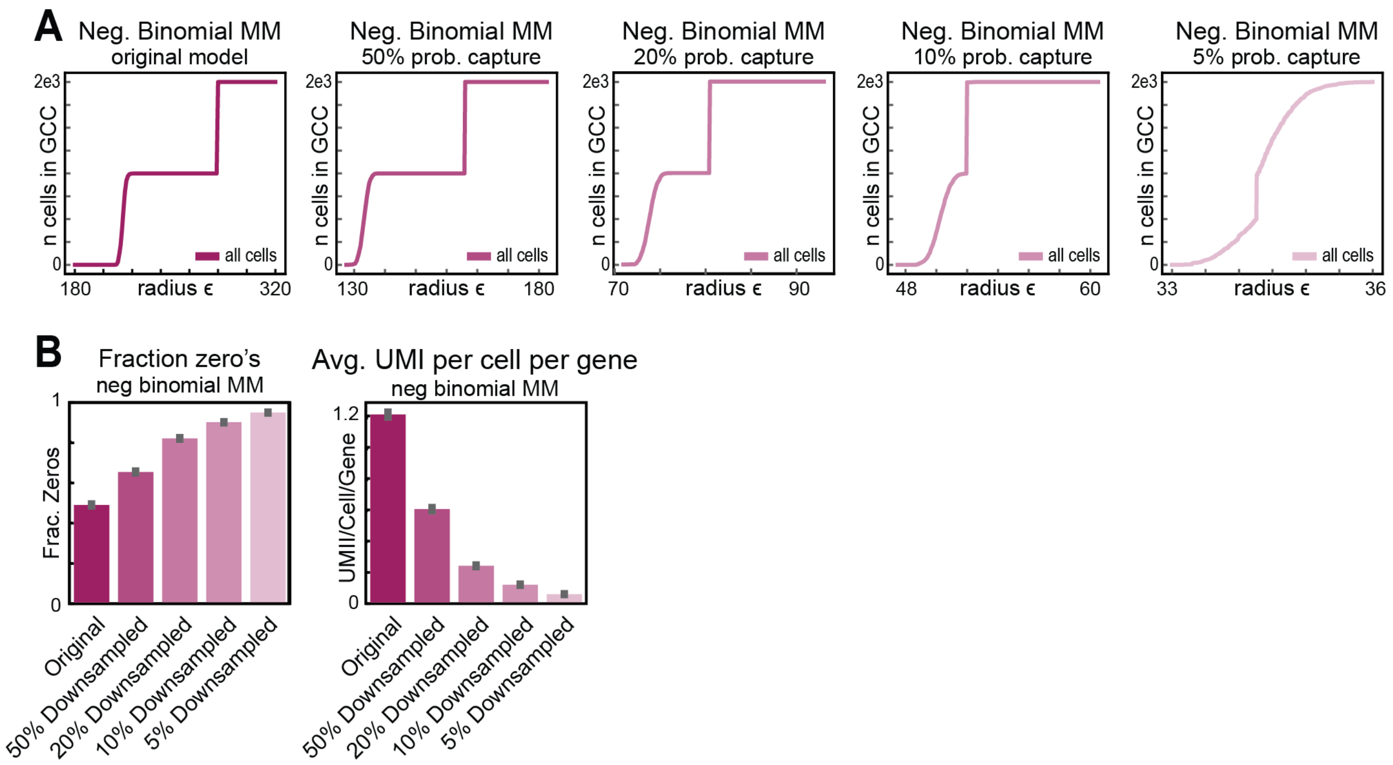


**Figure S3.3.** $\epsilon$ network analysis can identify distinct distributions in mixture models with noise and low marker gene expression. Briefly, each synthetic dataset modeled a mixture of two distributions. In each distribution, 1000 points were sampled from a 10,000 dimensional feature space. For each group of 1,000 points, only 250 of the 10,000 features varied in magnitude across two 'cell type' groups, in order to model a situation where only a fraction of marker genes demonstrate differential expression levels. To simulate different levels of sparsity, the data was downsampled such that each UMI count was subject to a Bernoulli trial at the indicated capture probability. **A)** Plot of the size of the giant component over increasing radius $\epsilon$ for $\epsilon$ networks constructed using a Negative Binomial Mixture model with varying levels of downsampling. Model construction details are provided in the Methods section description of Synthetic Negative Binomial Data. **B)** Fraction of zeros and average UMI count per cell per gene were calculated for each model and plotted as a bar graph.

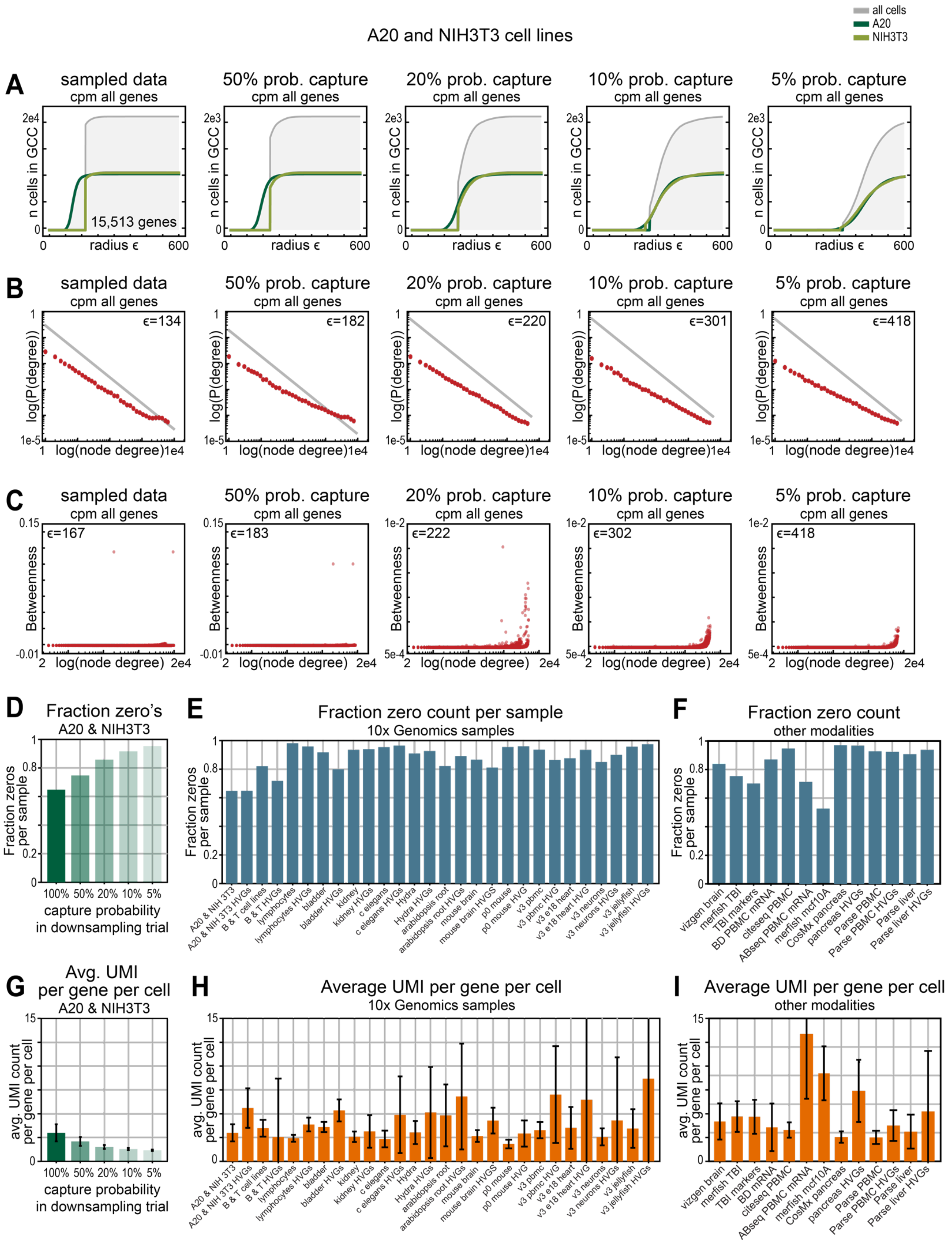

all cells
A20
NIH3T3
A20 and NIH3T3 cell lines
A
sampled data
cpm all genes
50% prob. capture
20% prob. capture
10% prob. capture
5% prob. capture
n cells in GCC
2e4
2e3
0
radius ε
600
15,513 genes
B
log(P(degree))
log(node degree)
1e-5
1e4
ε=134
ε=182
ε=220
ε=301
ε=418
C
Betweenness
0.15
-0.01
1e-2
5e-4
2e4
ε=167
ε=183
ε=222
ε=302
ε=418
D
Fraction zero's
A20 & NIH3T3
Fraction zeros per sample
100% 50% 20% 10% 5%
capture probability in downsampling trial
E
Fraction zero count per sample
10x Genomics samples
A20 & NIH 3T3
A20 & NIH 3T3 HVGs
B & T cell lines
B & T HVGs
lymphocytes
lymphocytes HVGs
bladder
bladder HVGs
kidney
kidney HVGs
c elegans
c elegans HVGs
Hydra
Hydra HVGs
arabidopsis root
arabidopsis root HVGs
mouse brain
mouse brain HVGS
p0 mouse
p0 mouse HVG
v3 pbmc
v3 pbmc HVG
v3 e18 heart
v3 e18 heart HVG
v3 neurons
v3 neurons HVGs
v3 jellyfish
v3 jellyfish HVGs
F
Fraction zero count
other modalities
vizgen brain
merfish TBI
TBI markers
BD PBMC mRNA
citeseq PBMC
ABseq PBMC mRNA
merfish mcf10A
CosMx pancreas
pancreas HVGs
Parse PBMC
Parse PBMC HVGs
Parse liver
Parse liver HVGs
G
Avg. UMI per gene per cell
A20 & NIH3T3
avg. UMI count per gene per cell
H
Average UMI per gene per cell
10x Genomics samples
I
Average UMI per gene per cell
other modalities

**Figure S3.4.** Analysis of how sparsity effects separability of biological data. **A)** Plot of the size of the giant component over increasing radius $\epsilon$ for $\epsilon$ networks constructed using the CPM transformed counts from the A20 and NIH 3T3 cell line data. For each panel, the original raw count matrix was down-sampled by performing a Bernoulli trial for each UMI count at the indicated capture probability. After down-sampling, the count matrices were CPM transformed. **B)** Density distributions for the down-sampled CPM-normalized A20 and NIH 3T3 cell line data described in (A). **C)** Plots of node degree versus betweenness centrality versus for the down-sampled CPM-normalized A20 and NIH3T3 cell line data across varying capture probabilities. **D-F)** Fraction of zero counts in **D)** the down-sampled A20 and NIH 3T3 cell line data across varying capture probabilities, **E)** feature-selected scRNA-seq data generated from the 10x Genomics platform, **F)** feature-selected single-cell transcriptomic data from other experimental modalities. **G-I)** Average UMI count per gene per cell in **G)** the down-sampled A20 and NIH 3T3 cell line data across varying capture probabilities, **H)** feature-selected scRNA-seq data generated from the 10x Genomics platform, **I)** feature-selected single-cell transcriptomic data from other experimental modalities

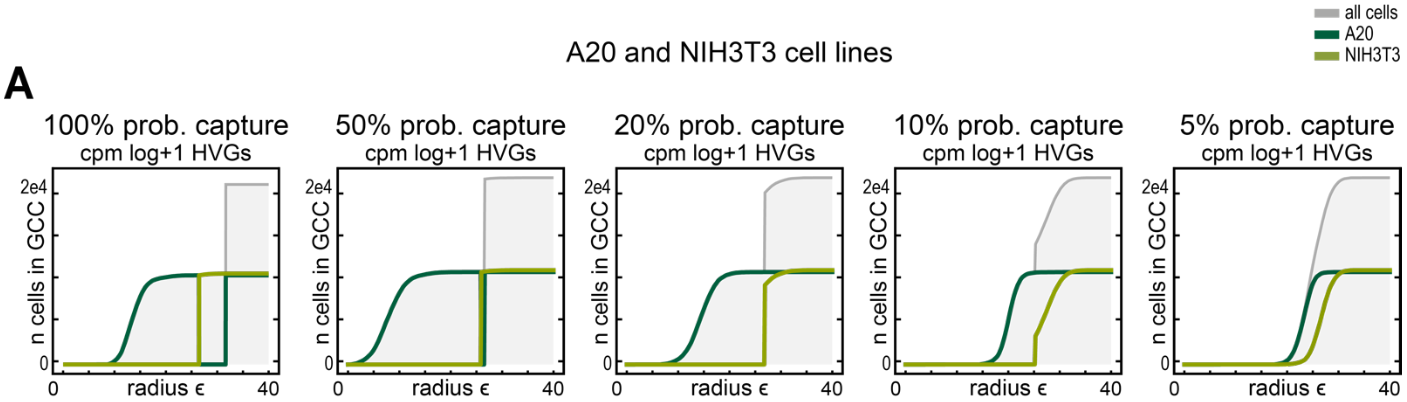


**Figure S3.5.** Analysis of how sparsity effects separability of biological data in HVG-selected, transformed spaces. **A)** Plot of the size of the giant component over increasing radius $\epsilon$ for $\epsilon$ networks constructed using the CPM-Log+1 transformed HVG counts from the A20 and NIH 3T3 cell line data. For each panel, the original raw count matrix was down-sampled by performing a Bernoulli trial for each UMI count at the indicated capture probability. After down-sampling, the original set of HVGs were subset, then the count matrices were CPM-Log+1 transformed.

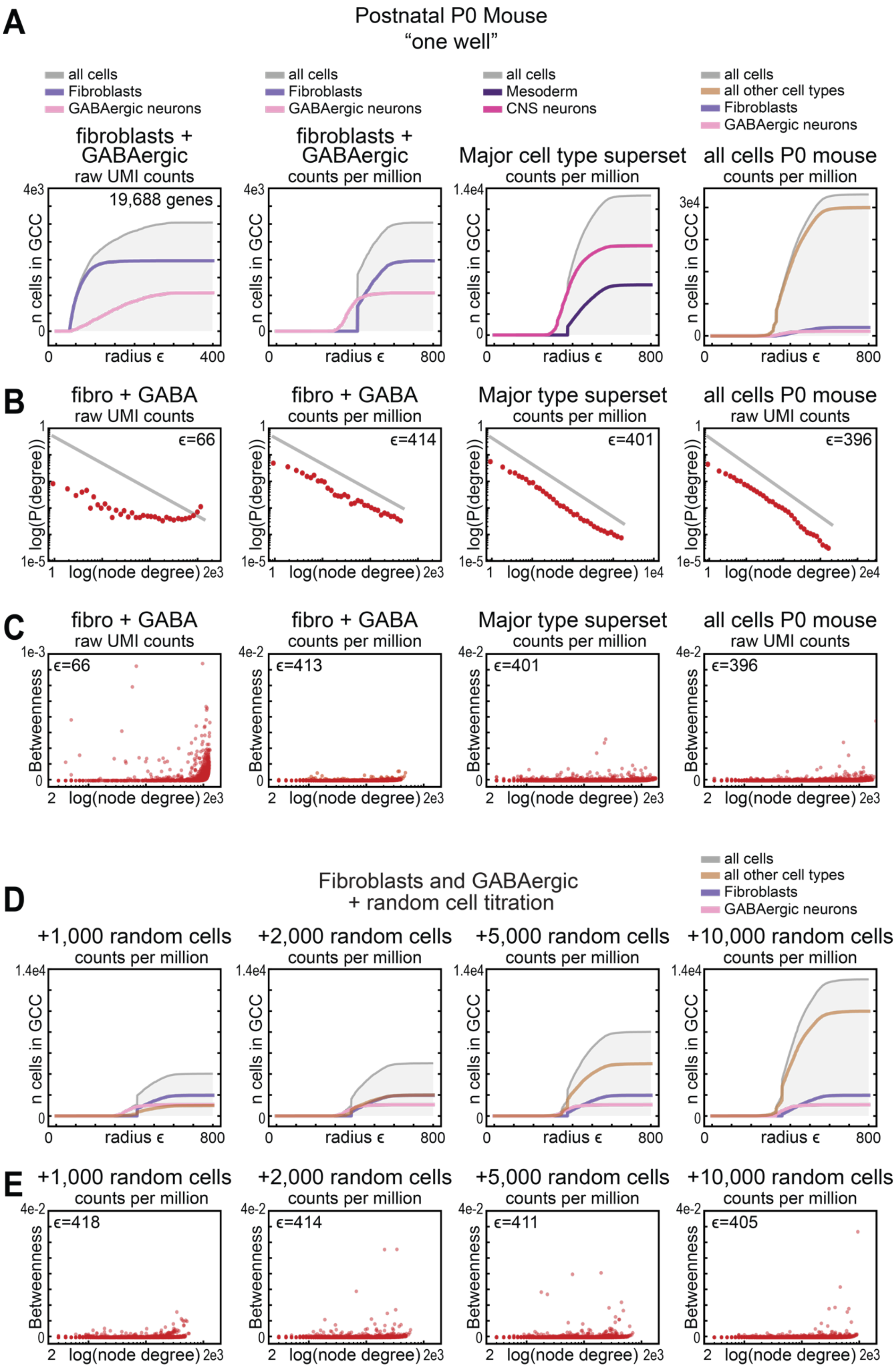
A
Postnatal P0 Mouse
"one well"
all cells
Fibroblasts
GABAergic neurons
fibroblasts + GABAergic
raw UMI counts
19,688 genes
counts per million
Mesoderm
CNS neurons
Major cell type superset
all other cell types
all cells P0 mouse
n cells in GCC
radius ϵ
B
fibro + GABA
Major type superset
log(P(degree))
log(node degree)
ϵ=66
ϵ=414
ϵ=401
ϵ=396
C
Betweenness
ϵ=413
D
Fibroblasts and GABAergic
+ random cell titration
+1,000 random cells
+2,000 random cells
+5,000 random cells
+10,000 random cells
E
ϵ=418
ϵ=411
ϵ=405

**Figure S3.6.** Cell type separability in single-cell atlas mouse organogenesis data. **A-E)** For each panel $\epsilon$ networks were constructed using scRNA-seq data from a single P0 stage and a single well. In the atlas, cell type labels were determined by applying the standard CPM-log+1-HVG-PCA set of transforms followed by two rounds of Louvain clusters. The first round produced major supersets which were then subclustered to identify specific cell-types. **A)** The size of the giant component was plotted over increasing radius $\epsilon$, when the $\epsilon$ networks were constructed using the following data from left to right: (i) raw UMI counts from Fibroblast and GABAergic cells, (ii) CPM-transformed counts from Fibroblast and GABAergic cells, (iii) CPM-transformed counts from the superset containing Fibroblast and the superset containing GABAergic cells, (iv) all cells from one well of the P0 embryo. **B)** Density distributions plotting the node degree versus the probability of observing cells at that node degree and the indicated $\epsilon$ for the datasets described in (A). **C)** Scatter plots displaying the betweenness centrality versus node degree at the indicated $\epsilon$ for the datasets described in (A). **D)** $\epsilon$ network analysis for Fibroblasts and GABAergic cells with titrated numbers of randomly selected cells included from the same well and the same P0 mouse embryo. The size of the giant component was plotted over increasing radius $\epsilon$, when the $\epsilon$ networks were constructed using CPM-transformed counts from Fibroblasts and GABAergic cells and an additional 1,000, 2,000, 5,000, or 10,000 random cells included. **E)** Density distributions plotting the node degree versus the probability of observing cells at that node degree and the indicated $\epsilon$ for the random cell titration data described in (D).

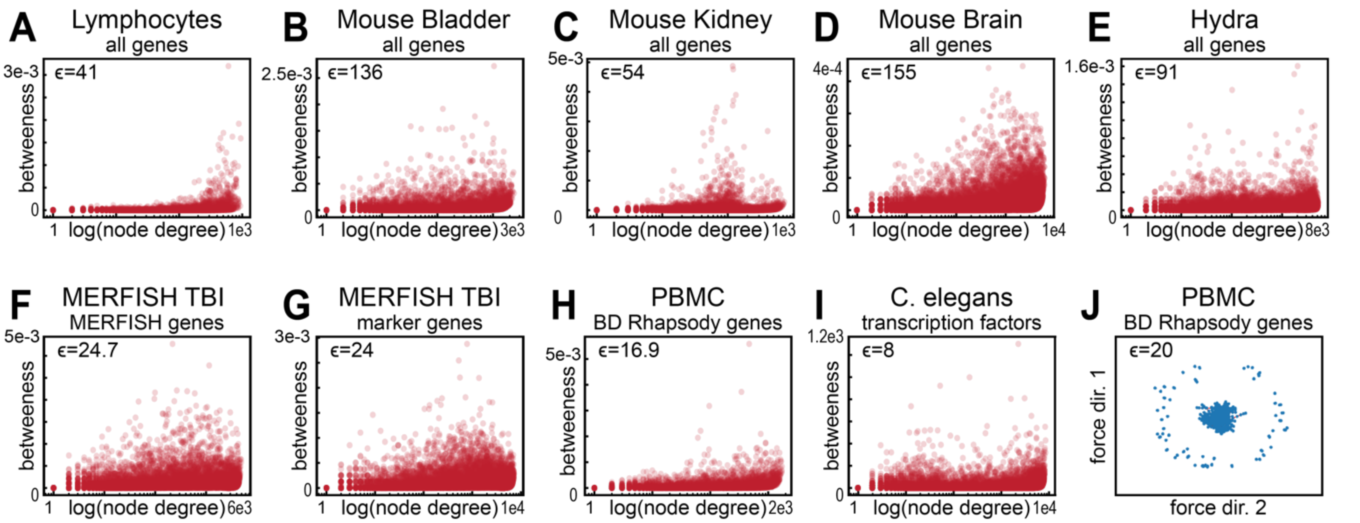


**Figure S3.7.** A lack of evidence for modularity in the giant component of single cell data. **A-I)** Scatter plots displaying the betweenness centrality versus node degree at the indicated $\epsilon$ for **A)** FACS-separated lymphocyte scRNA-seq data, **B)** Mouse Bladder scRNA-seq data, **C)** Mouse Kidney scRNA-seq data, **D)** Mouse Brain scRNA-seq data, **E)** Hydra scRNA-seq data, **F)** 170 targeted MERFISH genes from Traumatic Brain Injury (TBI) MERFISH data, **G)** 96 targeted brain marker genes from TBI MERFISH data, **H)** 499 targeted immune panel genes sequenced in Peripheral Blood Mononuclear Cells (PBMC) on the BD Rhapsody platform, and **I)** 677 transcription factors identified in C. elegans scRNA-seq data. **J)** Force Directed Layout of PBMC data sequenced on the BD Rhapsody platform.

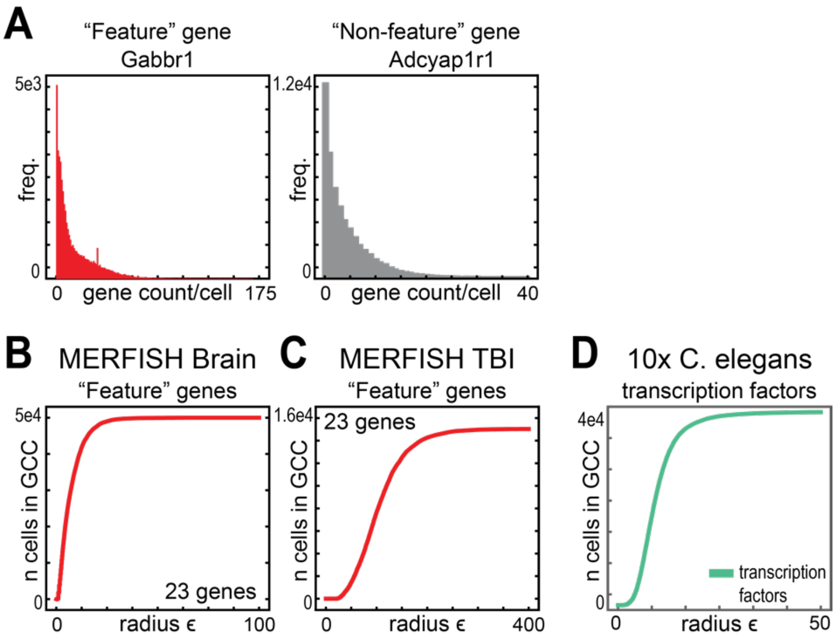


**Figure S4.1.** Visual selection of multi-modal genes does not produce distinct clusters of cells. **A)** Histograms displaying the distribution of UMI counts across all cells in the MERFISH brain data for the indicated gene. In the left panel, the Gabbr1 gene appears multimodal, and was included in the set of what is defined as "Feature" genes. In the right panel, the Adcyap1r1 has a unimodal distribution, and is excluded from the set of "Feature" genes. **B,C)** Plot of the size of the giant component over increasing radius $\epsilon$, when the $\epsilon$ networks were constructed using only the 23 multimodal "Feature" genes identified in **B)** MERFISH Brain data, **C)** MERFISH Traumatic Brain Injury (TBI) data an **D)** when only transcription factors from C. elegans were for $\epsilon$ network construction.

### Figure S4.2

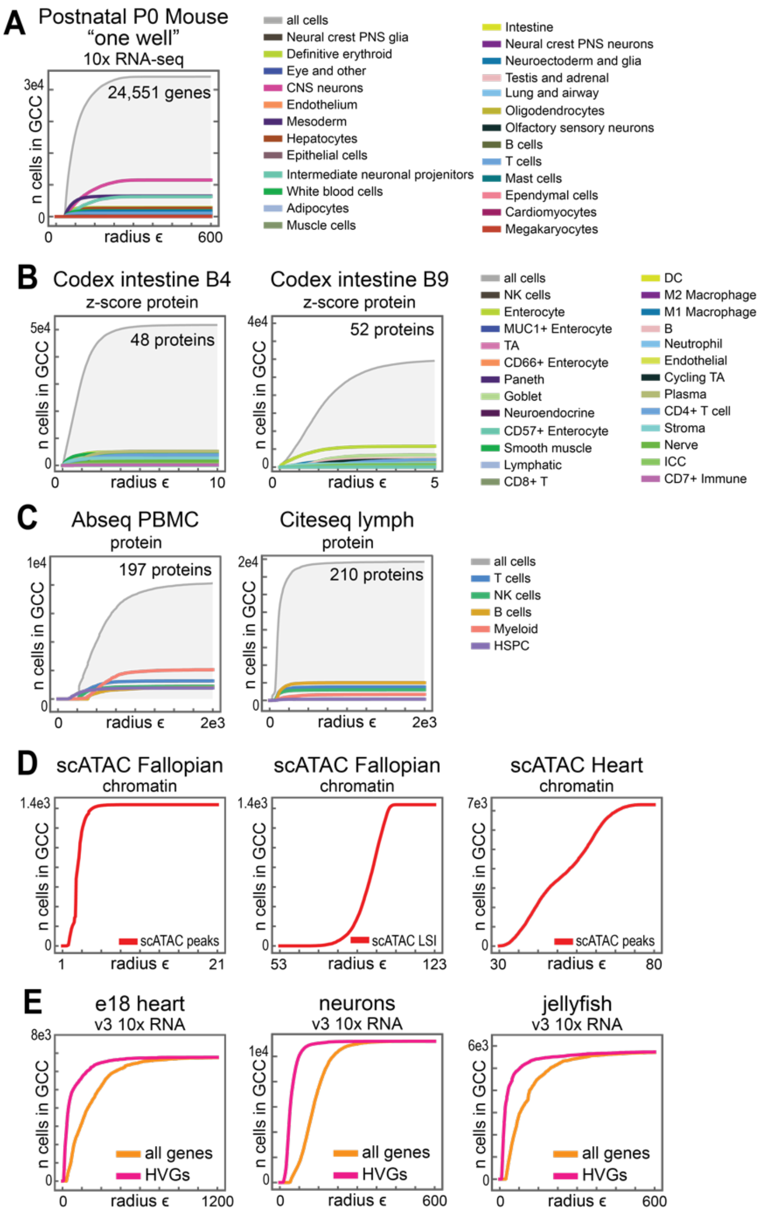


**Figure S4.2.** $\epsilon$ network analysis of data from different single-cell measurement platforms. **A-D)** Size of the giant component as a function of $\epsilon$ for data generated using **A)** one well of one P0 mouse, **B)** genome-wide single-cell proteomics data from two different intestine samples, **C)** targeted proteomics platforms, **D)** single-cell chromatin accessibility platforms, **E)** single-cell mRNA-seq data generated using the V3 Chemistry protocol of 10x Genomics. Additional details of each dataset are available in Table S1.

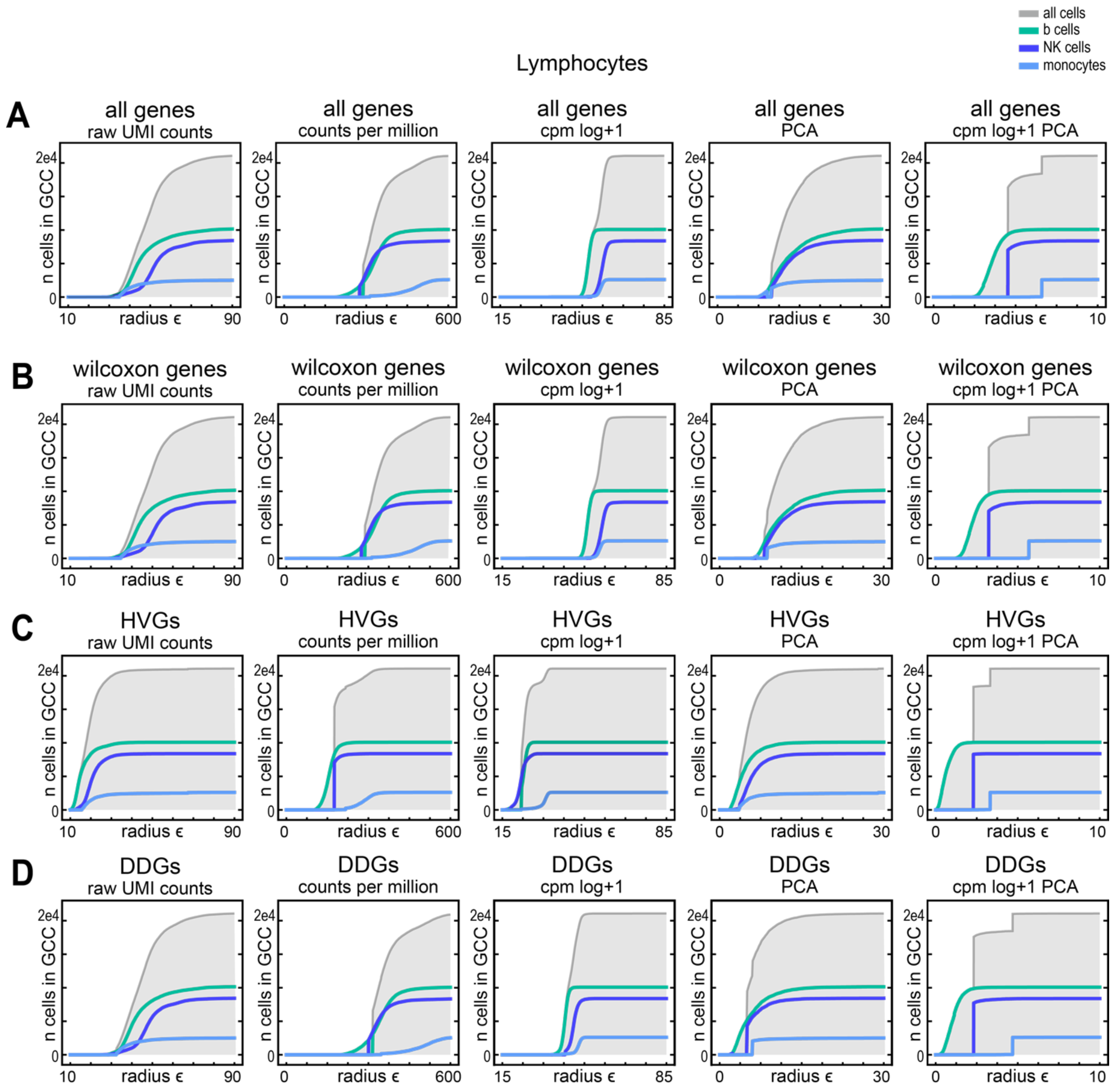


**Figure S5.1.** $\epsilon$ network analysis of feature-selected lymphocytes after common non-linear transformations. For each panel $\epsilon$ networks were constructed for the FACS-separated lymphocytes after no transformation, counts per million normalization (CPM), CPM log+1 transformation, Principal Component Analysis (PCA), or a combination of CPM, then log+1 then PCA. Then, the size of the giant component was plotted over increasing radius $\epsilon$, when the $\epsilon$ networks were constructed after the indicated transformations on either the set of **A)** all genes, **B)** Wilcoxon rank sum genes, **C)** Highly variable genes (HVGs), or **D)** Differentially distributed genes (DDGs).

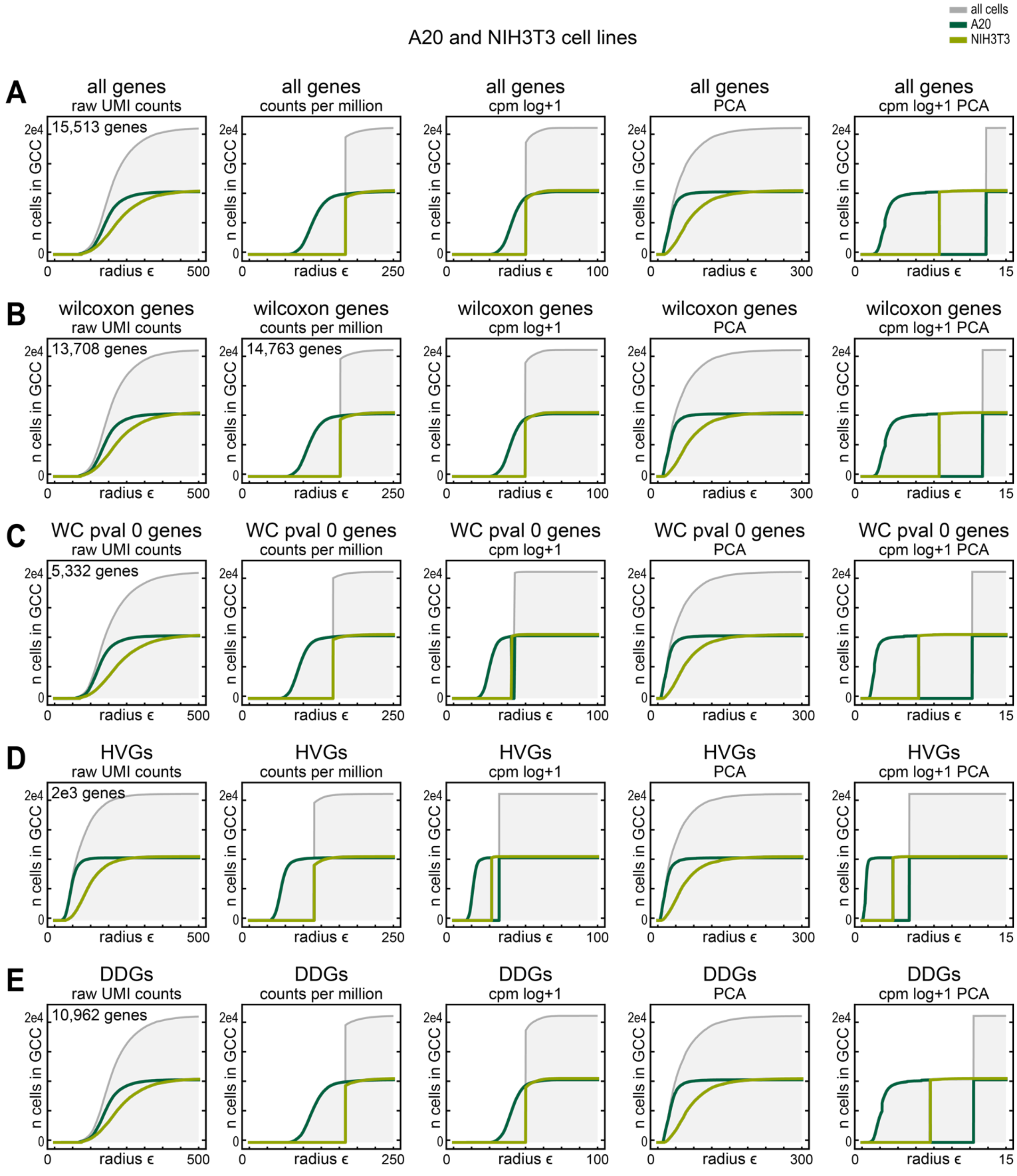


**Figure S5.2.** $\epsilon$ network analysis of feature-selected cell line data after common non-linear transformations. For each panel $\epsilon$ networks were constructed for the multiplexed A20 & NIH3T3 cell lines after no transformation, counts per million normalization (CPM), CPM log+1 transformation, Principal Component Analysis (PCA), or a combination of CPM, then log+1 then PCA. Then, the size of the giant component was plotted over increasing radius $\epsilon$, when the $\epsilon$ networks were constructed after the indicated transformations on either the set of **A)** all genes, **B)** Wilcoxon rank sum genes, **C)** the set of Wilcoxon rank sum genes with pvalues = 0, **D)** Highly variable genes (HVGs), or **E)** Differentially distributed genes (DDGs).

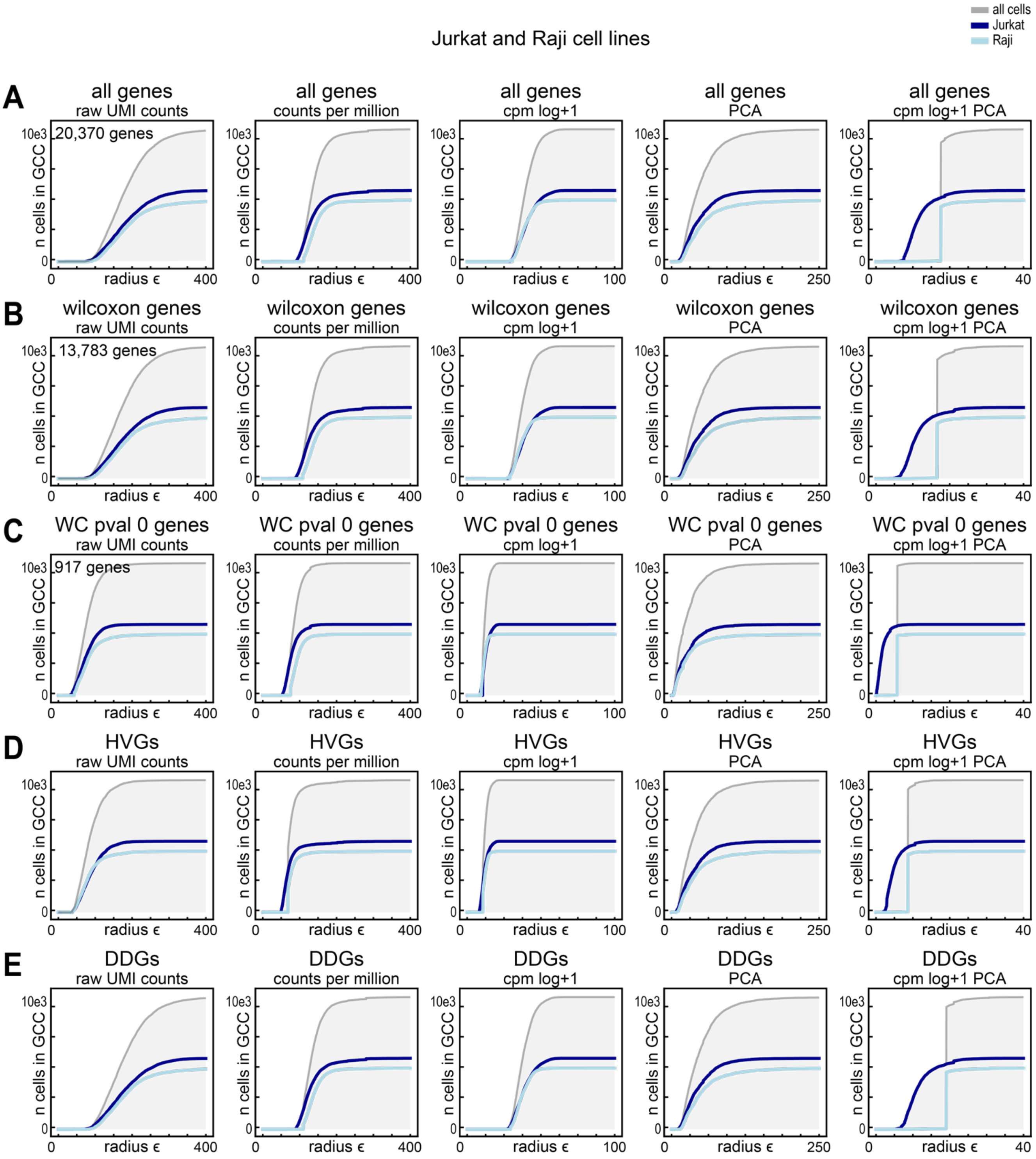


**Figure S5.3.** $\epsilon$ network analysis of feature-selected cell line data after common non-linear transformations. For each panel $\epsilon$ networks were constructed for the multiplexed Jurkat and Raji cell lines after no transformation, counts per million normalization (CPM), CPM log+1 transformation, Principal Component Analysis (PCA), or a combination of CPM, then log+1 then PCA. Then, the size of the giant component was plotted over increasing radius $\epsilon$, when the $\epsilon$ networks were constructed after the indicated transformations on either the set of **A)** all genes, **B)** Wilcoxon rank sum genes, **C)** the set of Wilcoxon rank sum genes with p-values = 0, **D)** Highly variable genes (HVGs), or **E)** Differentially distributed genes (DDGs).

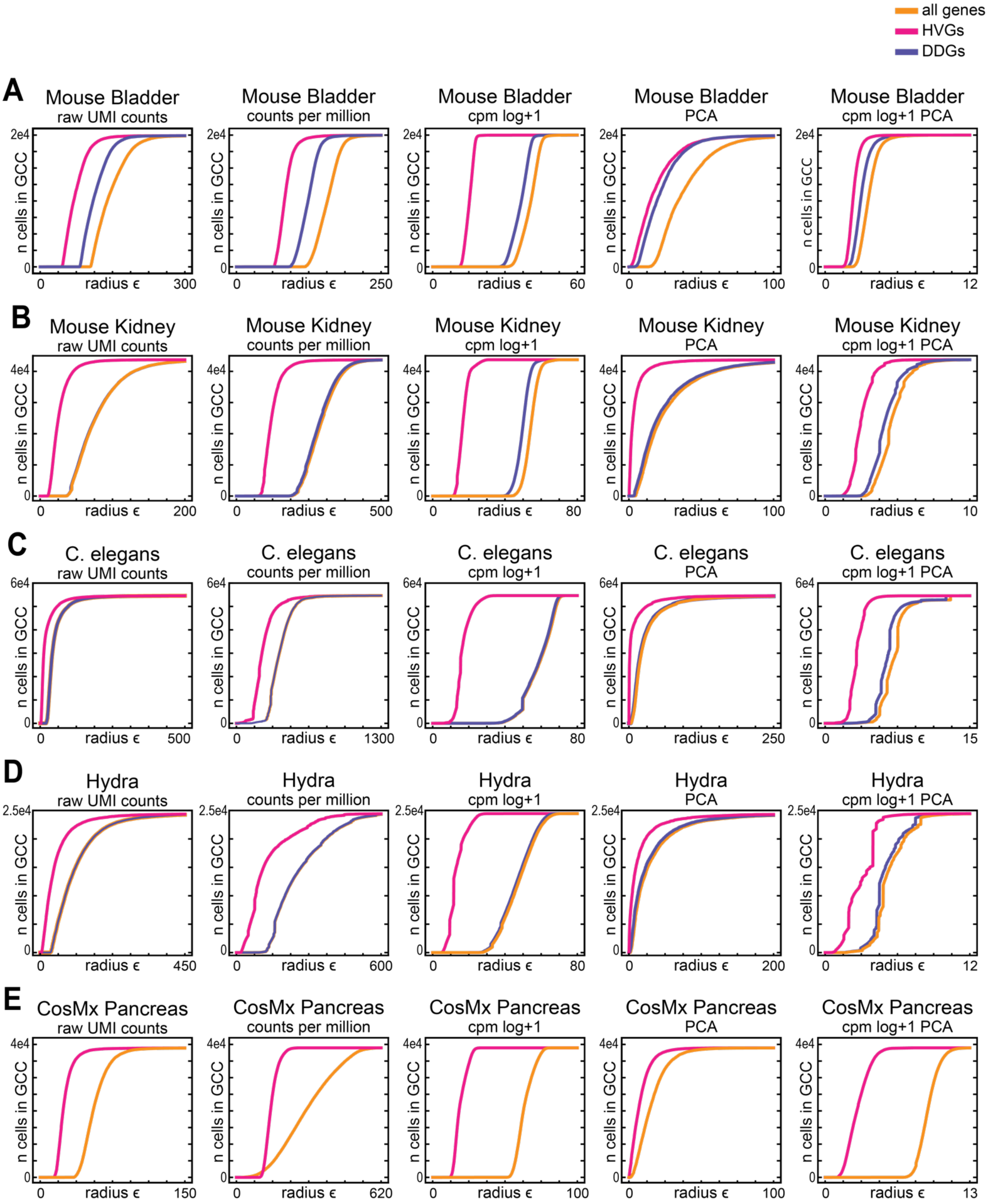


**Figure S5.4.** $\epsilon$ network analysis of feature-selected 10x Genomics scRNA-seq data after common non-linear transformations. For each panel $\epsilon$ networks were constructed after no transformation, counts per million normalization (CPM), CPM log+1 transformation, Principal Component Analysis (PCA), or a combination of CPM then log+1 then PCA. Then, the size of the giant component was plotted over increasing radius $\epsilon$, when the $\epsilon$ networks were constructed after the indicated transformations on either the set all genes (orange), Highly variable genes (HVGs) (pink), or Differentially Distributed Genes (DDGs) (purple), from the **A)** Mouse Bladder, **B)** Mouse Kidney, **C)** C. elegans, **D)** Hydra scRNA-seq data, or **E)** Pancreas CosMx data.

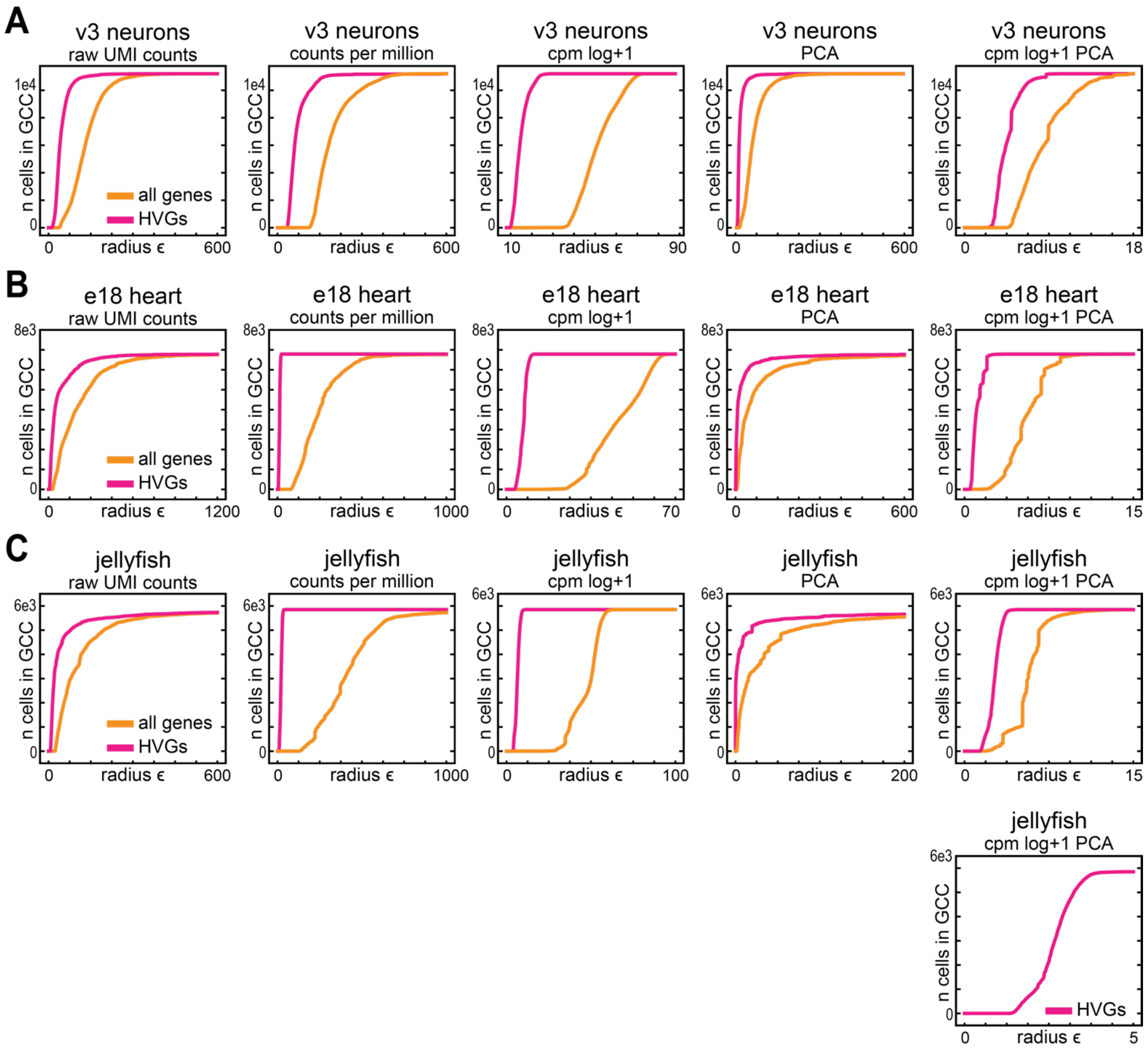


**Figure S5.5.** $\epsilon$ network analysis of feature-selected v3 Chemistry 10x Genomics scRNA-seq data after common non-linear transformations. For each panel $\epsilon$ networks were constructed after no transformation, counts per million normalization (CPM), CPM log+1 transformation, Principal Component Analysis (PCA), or a combination of CPM then log+1 then PCA. Then, the size of the giant component was plotted over increasing radius $\epsilon$, when the $\epsilon$ networks were constructed after the indicated transformations on either the set all genes (orange) or Highly variable genes (HVGs) (pink) from **A)** neurons, **B)** e18 heart, and **C)** fed jellyfish cells.

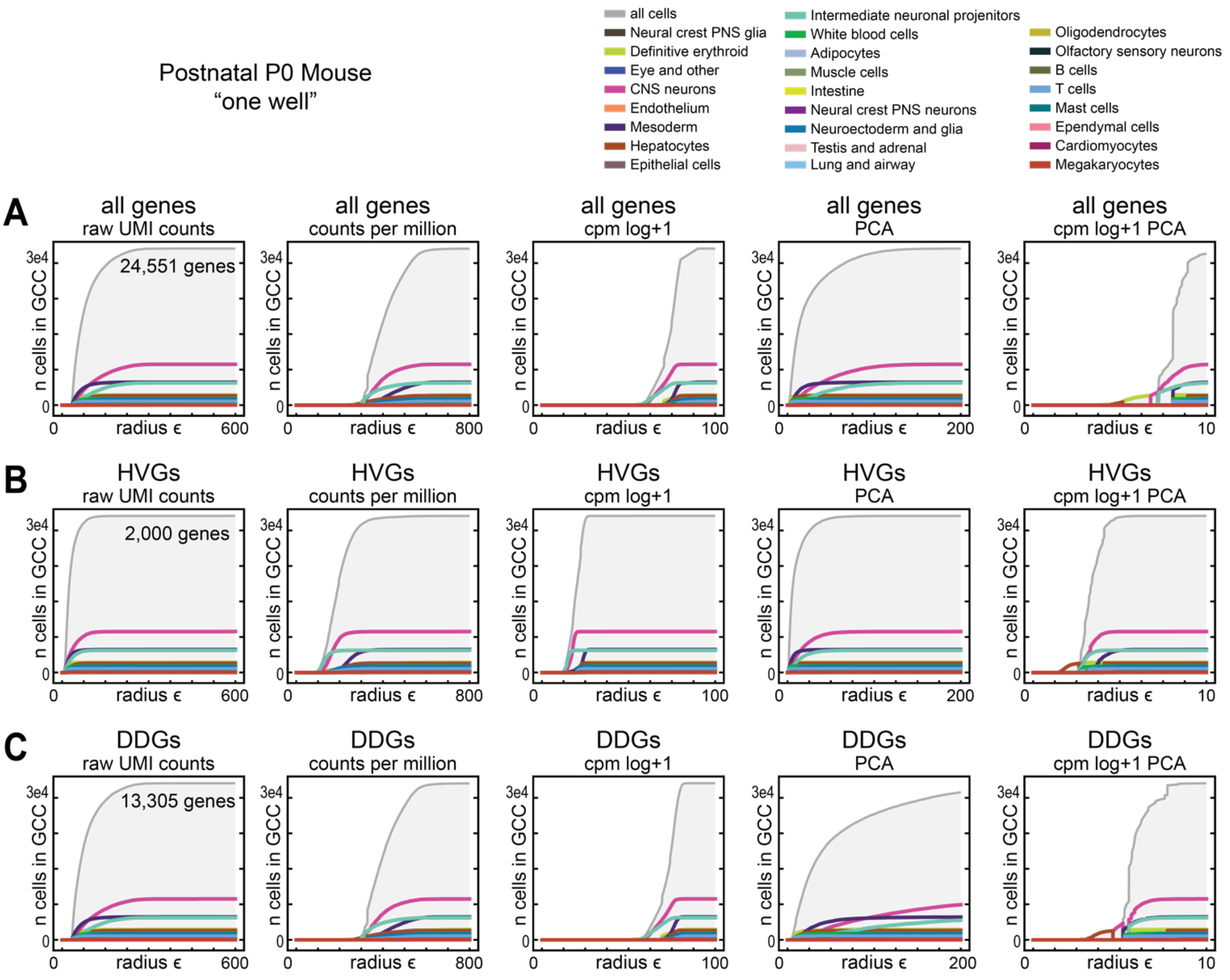


**Figure S5.6.** $\epsilon$ network analysis of feature-selected lymphocytes after common non-linear transformations. For each panel $\epsilon$ networks were constructed for the FACS-separated lymphocytes after no transformation, counts per million normalization (CPM), CPM log+1 transformation, Principal Component Analysis (PCA), or a combination of CPM, then log+1 then PCA. Then, the size of the giant component was plotted over increasing radius $\epsilon$, when the $\epsilon$ networks were constructed after the indicated transformations on either the set of **A)** all genes, **B)** Wilcoxon rank sum genes, **C)** Highly variable genes (HVGs), or **D)** Differentially distributed genes (DDGs).

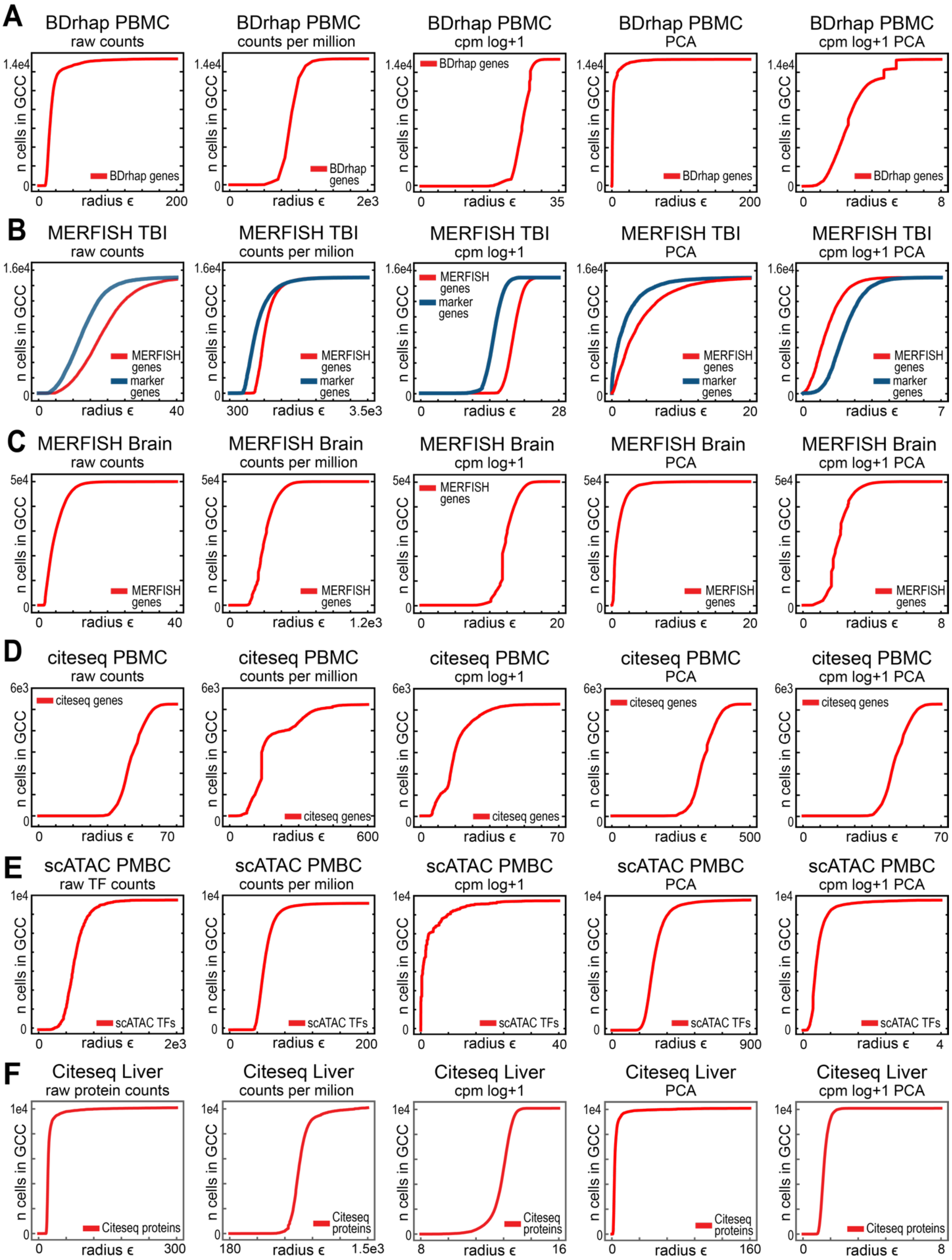


**Figure S5.7.** $\epsilon$ network analysis of different single-cell modalities after common non-linear transformations. For each panel $\epsilon$ networks were constructed after no transformation, counts per million normalization (CPM), CPM log+1 transformation, Principal Component Analysis (PCA), or a combination of CPM then log+1 then PCA. Then, the size of the giant component was plotted over increasing radius $\epsilon$, when the $\epsilon$ networks were constructed after the indicated transformations on **A)** PBMC mRNA data from the BD rhapsody platform, **B)** MERFISH genes or marker genes from the MERFISH Traumatic Brain Injury data, **C)** MERFISH Brain data, **D)** PBMC mRNA data from the CITE-seq platform, and **E)** the Transcription Factor (TF) counts of scATAC data collected from PBMCs.

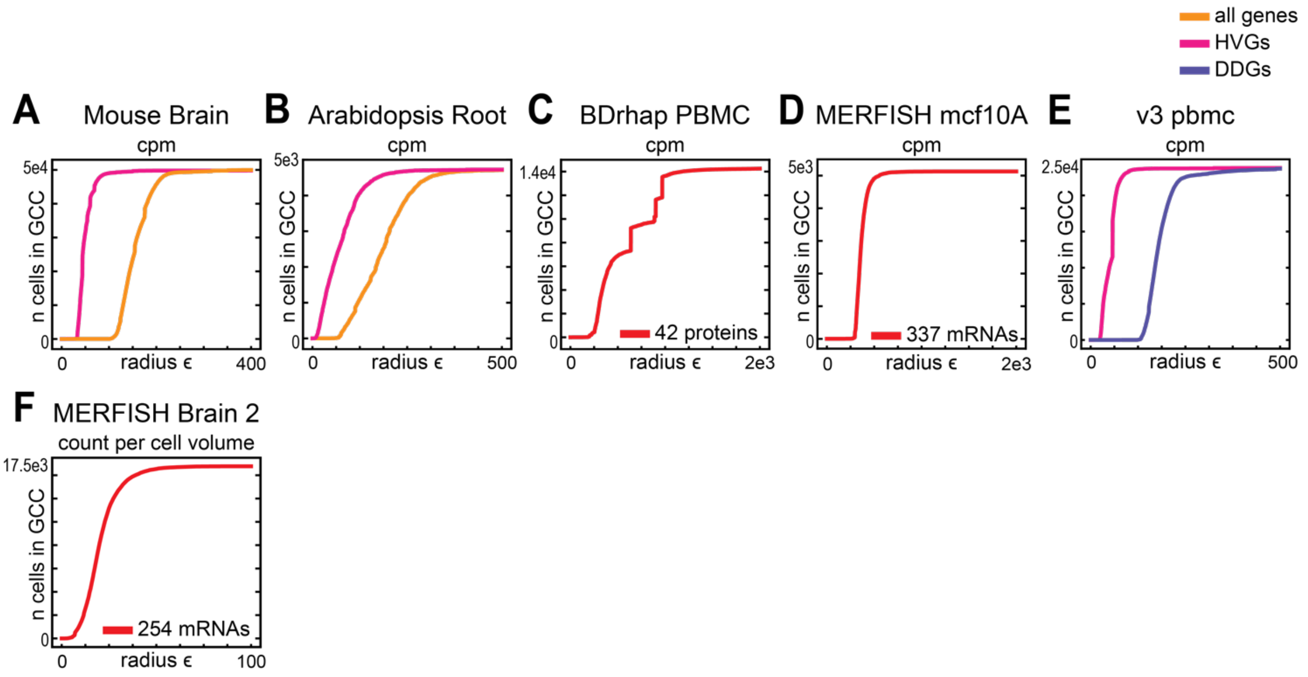


**Figure S5.8.** $\epsilon$ network analysis of different single-cell modalities after normalizing for cell volume. (A-E) For each panel $\epsilon$ networks were constructed after counts per million normalization (CPM), where each UMI count was divided by a size factor determined by the total number of counts in that cell. Then, the size of the giant component was plotted over increasing radius $\epsilon$, when the $\epsilon$ networks were constructed after the indicated transformations on **A)** PBMC mRNA data from the BD rhapsody platform, **B)** MERFISH genes or marker genes from the MERFISH Traumatic Brain Injury data, **C)** MERFISH Brain data, **D)** PBMC mRNA data from the CITE-seq platform, and **E)** the Transcription Factor (TF) counts of scATAC data collected from PBMCs.

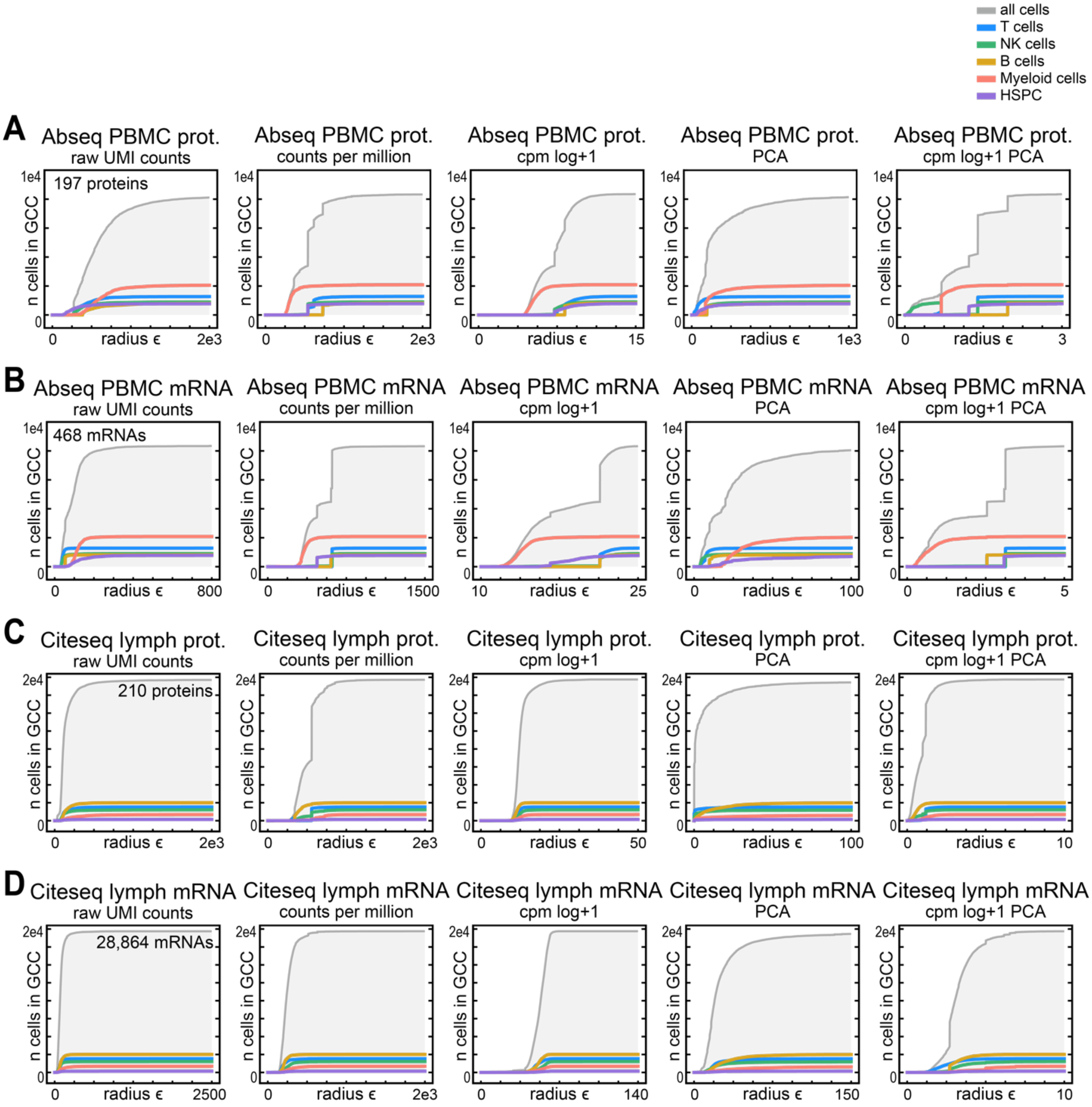


**Figure S5.9.** $\epsilon$ network analysis of different single-cell modalities after common non-linear transformations. For each panel $\epsilon$ networks were constructed after no transformation, counts per million normalization (CPM), CPM log+1 transformation, Principal Component Analysis (PCA), or a combination of CPM then log+1 then PCA. Then, the size of the giant component was plotted over increasing radius $\epsilon$, when the $\epsilon$ networks were constructed after the indicated transformations on **A)** PBMC protein data from the Ab-seq platform, **B)** PBMC mRNA data from the ABseq platform, **C)** Lymph protein data from the CITE-seq platform, and **D)** Lymph mRNA data from the CITE-seq platform.

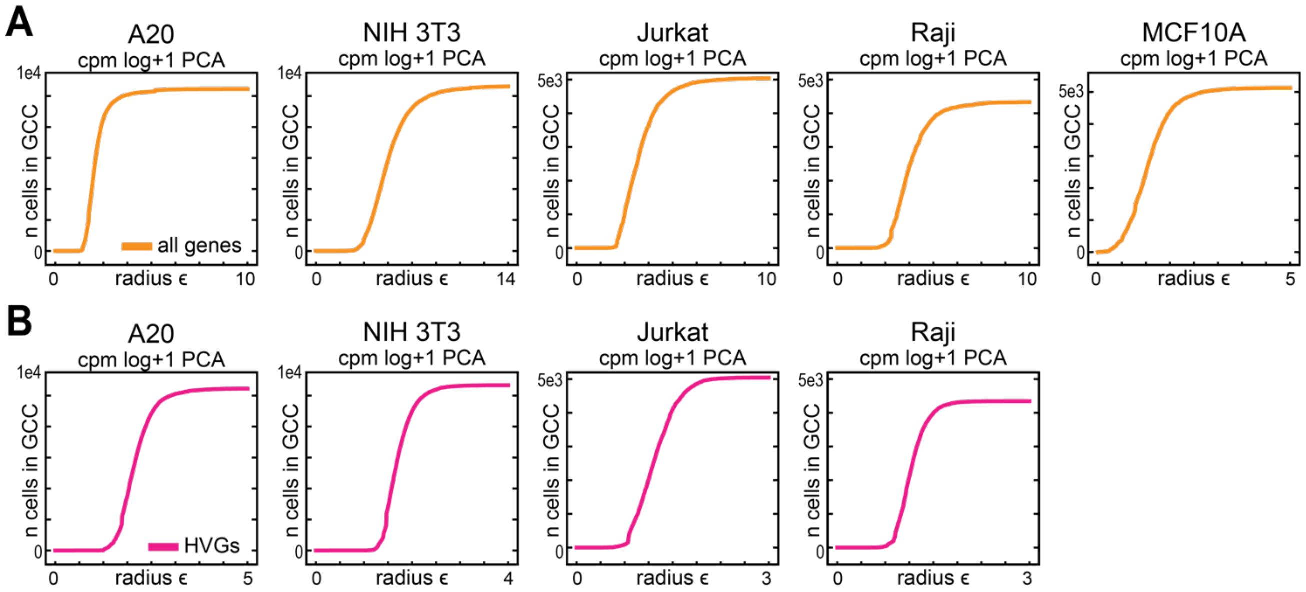


**Figure S5.10.** $\epsilon$ network analysis of data from one cell line after CPM-Log+1-PCA transformation. (For each panel $\epsilon$ networks were constructed after no transformation, counts per million normalization (CPM), CPM log+1 transformation, Principal Component Analysis (PCA), or a combination of CPM then log+1 then PCA. Then, the size of the giant component was plotted over increasing radius $\epsilon$, when the $\epsilon$ networks were constructed after the indicated transformations on **A)** all genes from the 10x Genomics data generated on the A20 cell line, the NIH 3T3 cell line, the Jurkat cell line, the Raji cell line, or the MERFISH MCF10A cell line, or **B)** the HVGs from the 10x Genomics data generated on the A20 cell line, the NIH 3T3 cell line, the Jurkat cell line, or the Raji cell line.

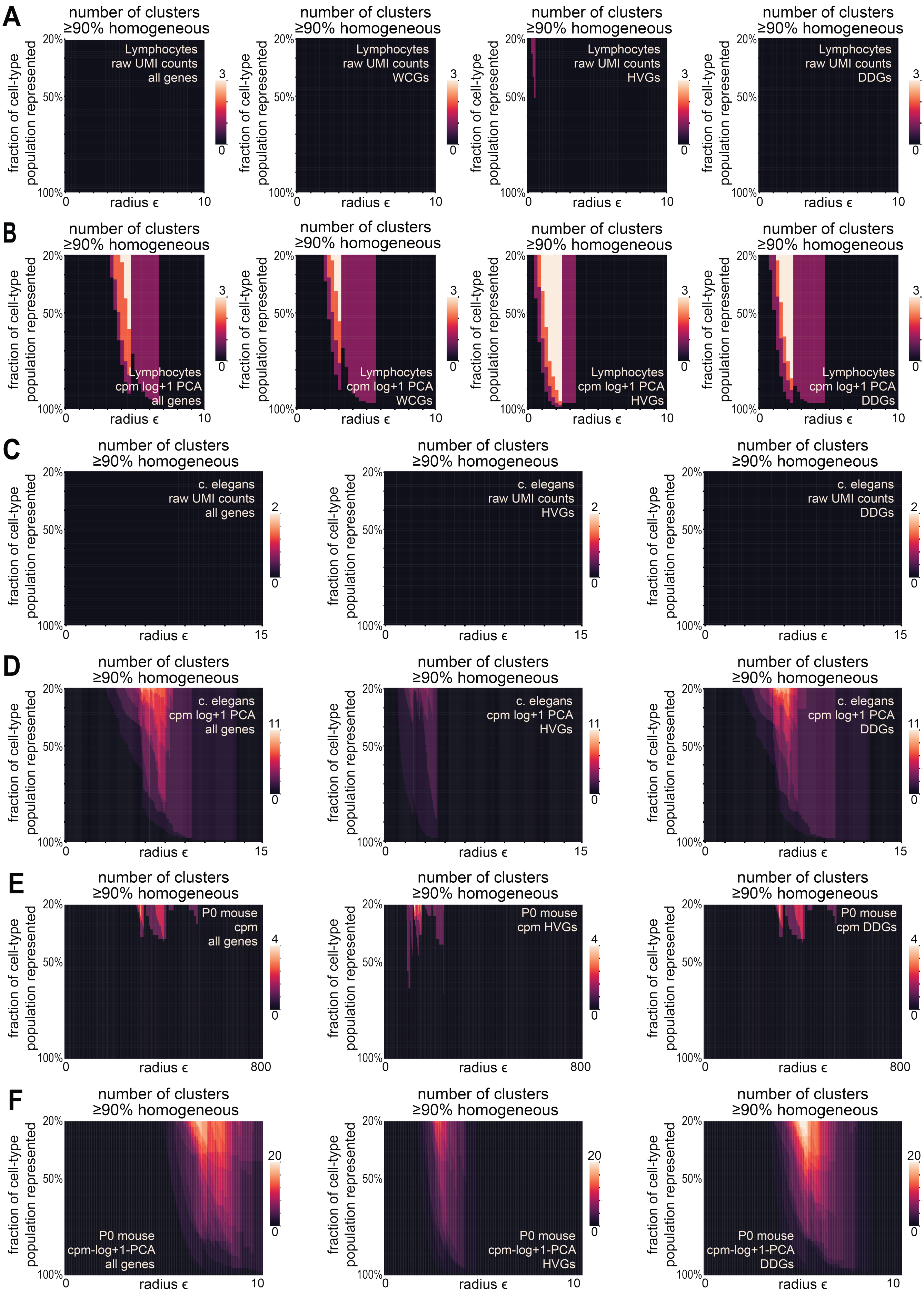
A
number of clusters ≥90% homogeneous
fraction of cell-type population represented
20%
50%
100%
radius ϵ
0
10
3
0
Lymphocytes raw UMI counts all genes
Lymphocytes raw UMI counts WCGs
Lymphocytes raw UMI counts HVGs
Lymphocytes raw UMI counts DDGs
B
Lymphocytes cpm log+1 PCA all genes
Lymphocytes cpm log+1 PCA WCGs
Lymphocytes cpm log+1 PCA HVGs
Lymphocytes cpm log+1 PCA DDGs
C
15
2
c. elegans raw UMI counts all genes
c. elegans raw UMI counts HVGs
c. elegans raw UMI counts DDGs
D
11
c. elegans cpm log+1 PCA all genes
c. elegans cpm log+1 PCA HVGs
c. elegans cpm log+1 PCA DDGs
E
800
4
P0 mouse cpm all genes
P0 mouse cpm HVGs
P0 mouse cpm DDGs
F
20
P0 mouse cpm-log+1-PCA all genes
P0 mouse cpm-log+1-PCA HVGs
P0 mouse cpm-log+1-PCA DDGs

**Figure S5.11.** Analysis of cluster composition for all clusters in the $\epsilon$ networks in raw and transformed data. **A-F)** Heatmap characterizing the size and composition of components in the $\epsilon$ networks for A,B) the FACS sorted lymphocytes, B,C) the *C. elegans* data, and D,E) the P0 mouse data. The x-axis indicates increasing radius $\epsilon$, the y-axis represents the fraction of particular cell types represented in each cluster, relative to the total number of that particular cell type in the data. The heatmap is colored by the number of components meeting that are at least 90% homogenous for a particular cell type. **A)** The raw UMI counts were used to construct $\epsilon$ networks for lymphocyte data, using either all genes, Wilcoxon rank sum genes (WCGs), Highly Variable Genes (HVGs), or Differentially Distributed Genes (DDGs). **B)** The counts per million, log + 1, principal component analysis (PCA) transformed counts were used to construct $\epsilon$ networks in for the lymphocyte data, using either all genes, Wilcoxon rank sum genes (WCGs), Highly Variable Genes (HVGs), or Differentially Distributed Genes (DDGs). **C)** The raw UMI counts were used to construct $\epsilon$ networks for the *C. elegans* data, using either all genes, Highly variable genes (HVGs), or Differentially distributed genes (DDGs). **D)** The counts per million, log + 1, principal component analysis (PCA) transformed counts were used to construct $\epsilon$ networks for the *C. elegans* data, using either all genes, Highly Variable Genes (HVGs), or Differentially Distributed Genes (DDGs). **E)** The raw UMI counts were used to construct $\epsilon$ networks for the P0 mouse data, using either all genes, Highly variable genes (HVGs), or Differentially distributed genes (DDGs). **F)** The counts per million, log + 1, principal component analysis (PCA) transformed counts were used to construct $\epsilon$ networks for the P0 mouse data, using either all genes, Highly Variable Genes (HVGs), or Differentially Distributed Genes (DDGs).

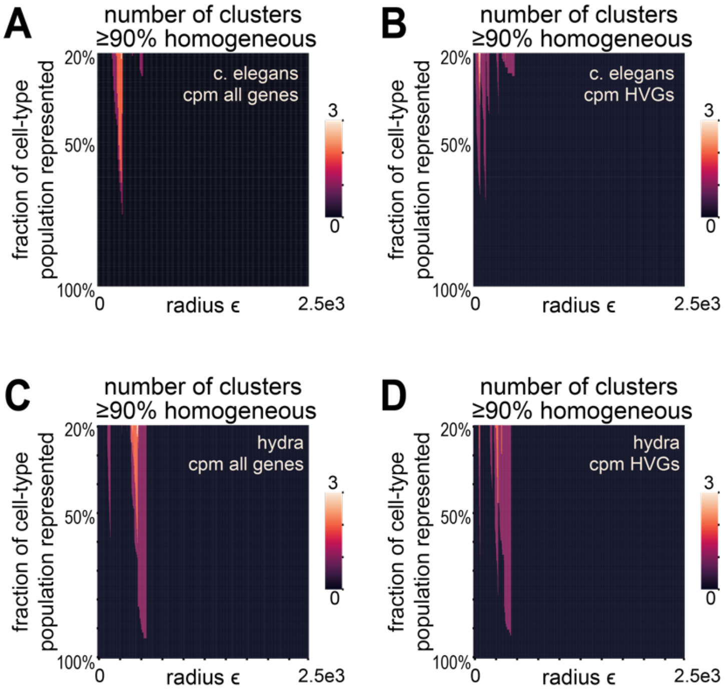


**Figure S5.12.** Analysis of cluster composition for all clusters in the $\epsilon$ networks using raw or counts per million (CPM) transformed data. Heatmap characterizing the size and composition of components in the $\epsilon$ networks for A-B) the *C. elegans* data and in C-D) the *Hydra vulgaris* data. The x-axis indicates increasing radius $\epsilon$, the y-axis represents the fraction of particular cell types represented in each cluster, relative to the total number of that particular cell type identified in the data. Here, each cell's identity was defined using the results of the clustering analysis reported in the manuscripts accompanying these published data. The heat map is colored by the number of components that are at least 90% homogenous for a particular cell type. $\epsilon$ networks were constructed using **A)** raw UMI counts for *C. elegans,* **B)** CPM transformed counts for *C. elegans,* **C)** raw UMI counts for *Hydra vulgaris,* or **D)** CPM transformed counts for *Hydra vulgaris.*

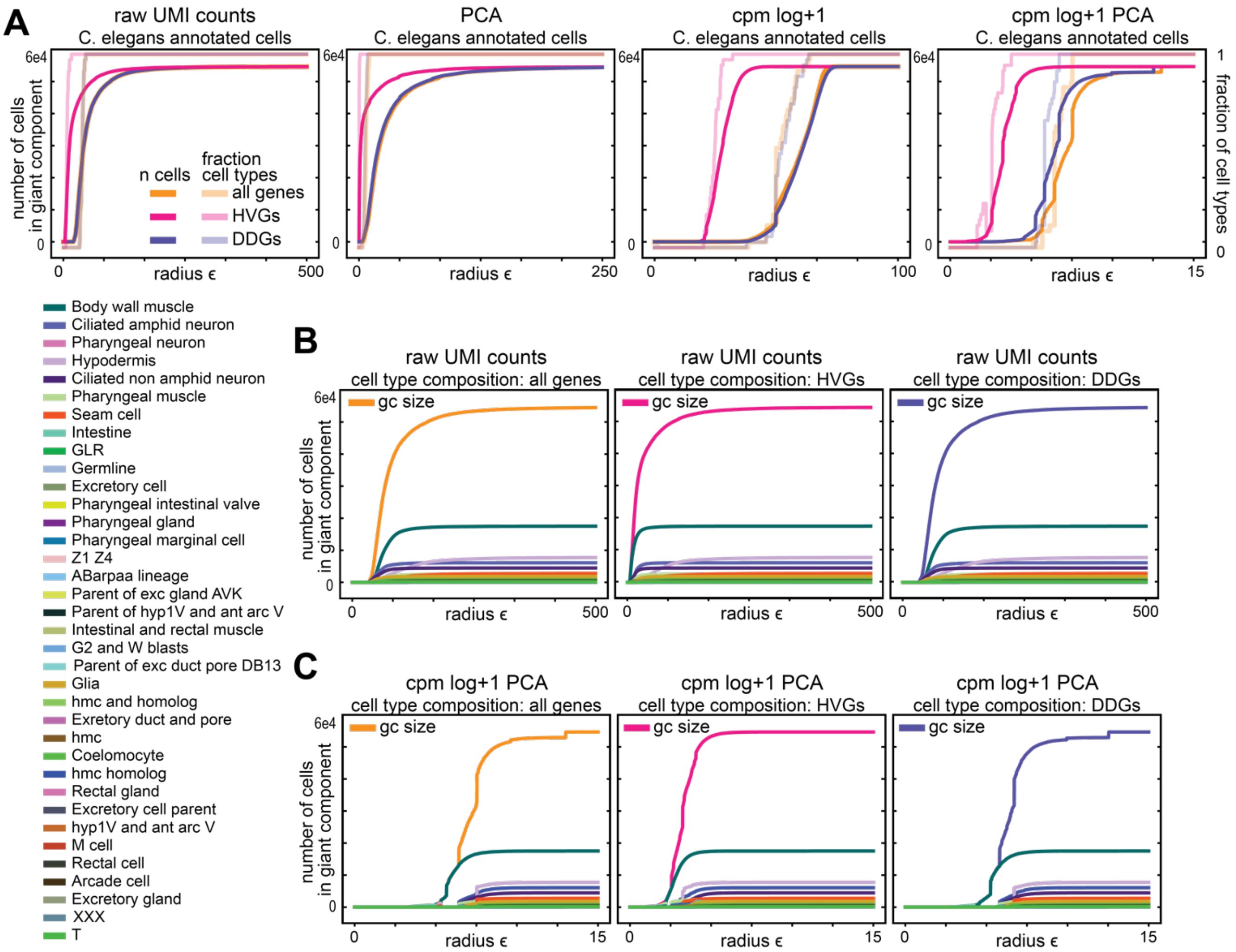


**Figure S5.13.** $\epsilon$ network analysis of annotated cell types from *C. elegans* data. In **A)** $\epsilon$ networks were constructed for scRNA-seq data after no transformation, principal component analysis (PCA), counts per million normalization (CPM) then log + 1 transformation, or CPM, then log+1, then PCA on either all genes, Highly Variable Genes (HVGs), or Differentially Distributed Genes (DDGs). In each panel, the number of cells in the giant component as well as the fraction of all cell types represented in the giant component are plotted over increasing radius $\epsilon$. **B)** For the raw UMI count data, the number of cells in the giant component for all cells and for cells from each annotated cell type were plotted as a function of radius $\epsilon$. In each panel, this analysis was performed on either all genes, HVGs, or DDGs. **C)** For the cpm log+1 PCA transformed data, the number of cells in the giant component for all cells and for cells from each annotated cell type were plotted as a function of radius $\epsilon$. In each panel, this analysis was performed on either all genes, HVGs, or DDGs. In all plots from this figure, cell type annotations were determined using the operationally standard approach described by the authors of the *C. elegans* study, and “unannotated” cells were excluded from the analysis.

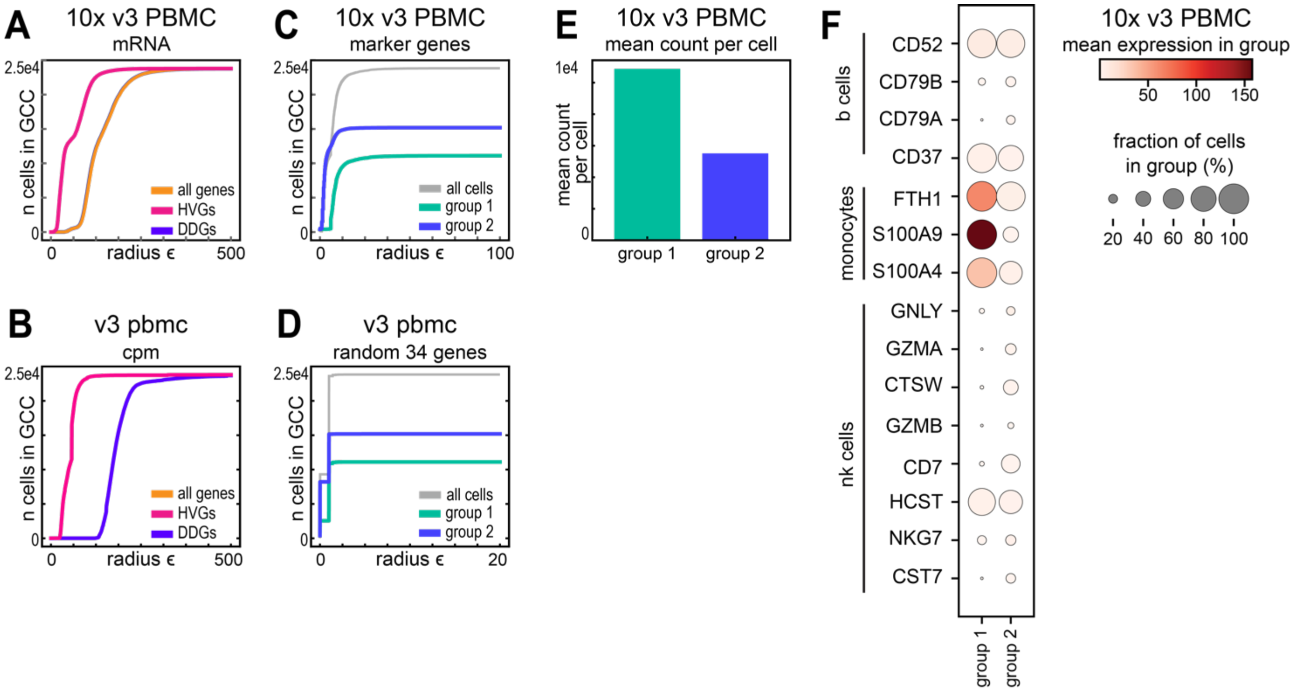


**Figure S5.14.** Two groups of cells in HVG-selected version 3 scRNA-seq data of PBMC cells are not marker gene specific. **A-B)** The size of the giant component was plotted over increasing radius $\epsilon$, when the $\epsilon$ networks were constructed using the 10x Genomics scRNA-seq with version 3 chemistry from PBMCs for all genes, Highly Variable Genes (HVGs), or Differentially Distributed Genes (DDGs), on **A)** raw UMI counts, or **B)** counts per million normalized counts. **C-D)** The size of the giant component was plotted over increasing radius $\epsilon$, when the $\epsilon$ networks were constructed on data from 10x Genomics scRNA-seq with version 3 chemistry from PBMCs genes using **C)** only 34 marker genes for B cells, monocytes and natural killer (NK) cells or **D)** a random selection of 34 genes. **E)** Mean UMI count per cell for the two groups of cells in HVG-selected version 3 scRNA-seq data of PBMC cells. **F)** Dot plot for the two groups showing mean marker gene expression and fraction of cells each marker gene is expressed in.

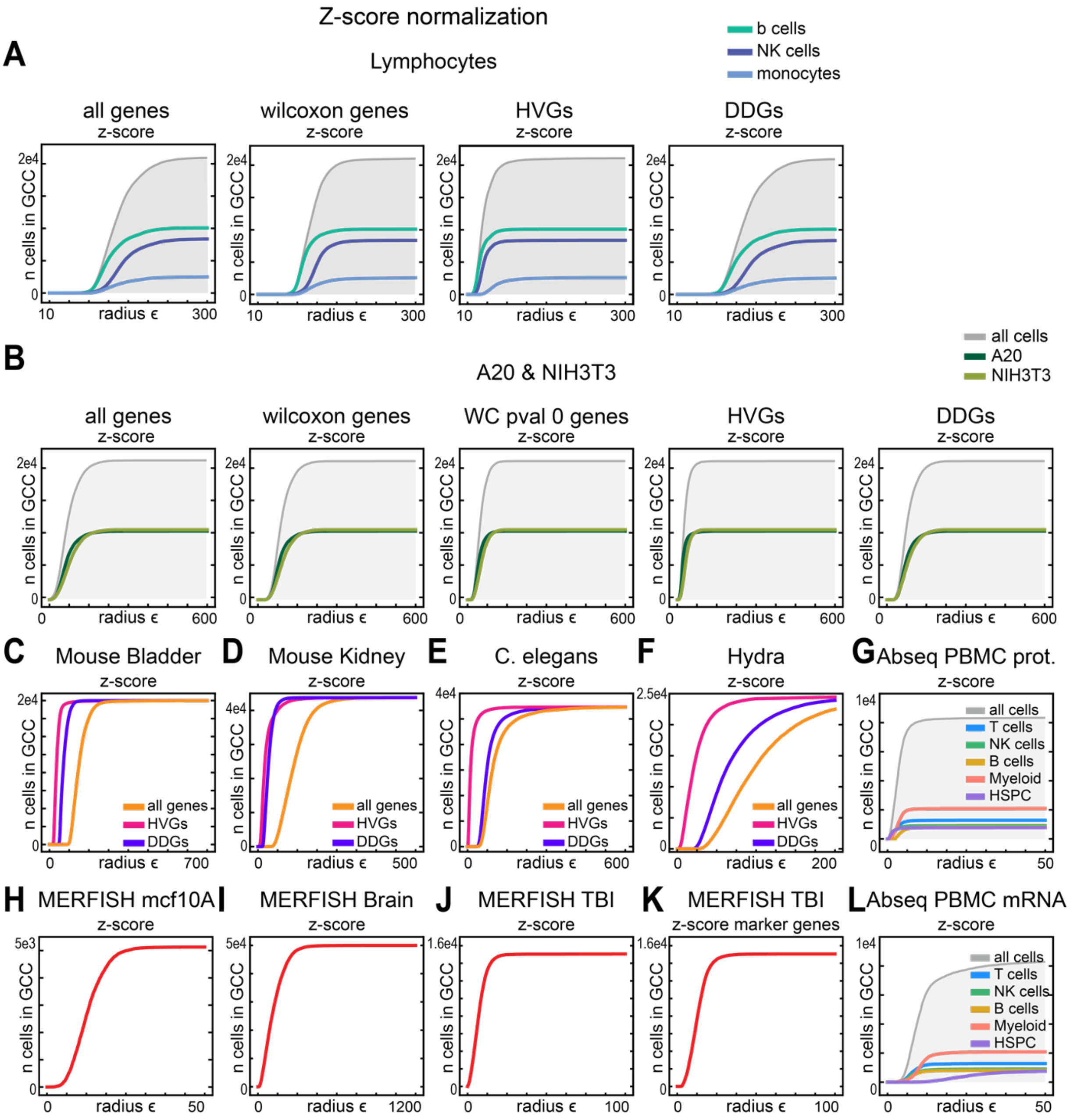


**Figure S5.15.** $\epsilon$ network analysis of z-score normalized data. For each panel $\epsilon$ networks were constructed after z-score normalization. The size of the giant component was plotted over increasing radius $\epsilon$, when the $\epsilon$ networks were constructed after z-score transformation of all genes, Highly Variable Genes (HVGs), Wilcoxon rank sum genes, or Differentially Distributed Genes (DDGs) from 10x Genomics scRNA-seq data from **A)** the FACs sorted lymphocytes, **B)** A20 & NIH3T3 cell lines, **C)** the Mouse bladder, **D)** the Mouse kidney, **E)** C. elegans embryo, or **F)** the Hydra. Giant component analysis was performed on z-score normalized data from **G)** ABseq PBMC proteins, the MERFISH genes from **H)** mcf10A cells, **I)** Mouse brain, **J)** Mouse brain traumatic brain injury data, **K)** marker genes from MERFISH traumatic brain injury data, or **L)** ABseq PBMC mRNA.

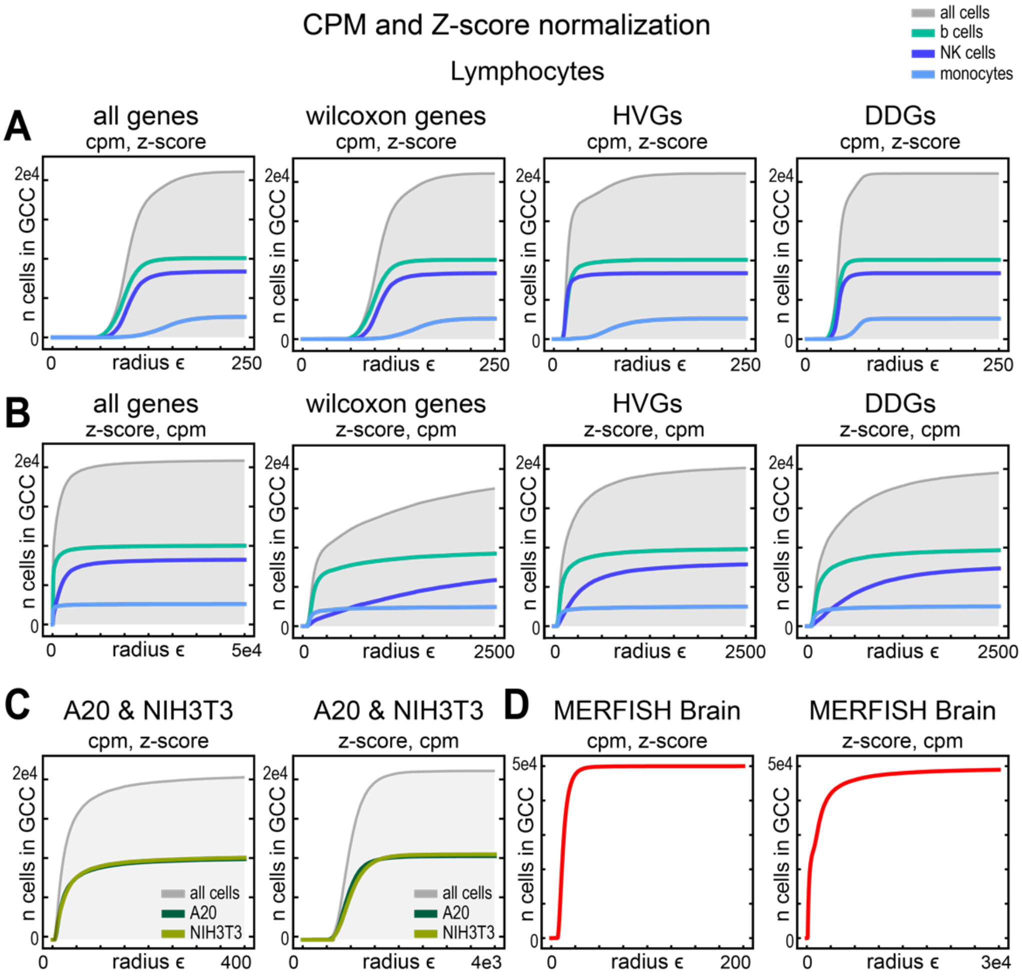


**Figure S5.16.** $\epsilon$ network analysis of data transformed with a combination of counts per million and z-score normalization. In **A)** Counts per million (cpm) then z-score normalization was performed on FACs sorted lymphocyte data using either all genes, Wilcoxon genes, Highly Variable Genes (HVGs), or Differentially Distributed Genes (DDGs). Then, the size of the giant component was plotted over increasing radius $\epsilon$. In **B),** Z-score then counts per million (cpm) normalization was performed on FACS purified lymphocyte data using either all genes, Wilcoxon genes, HVGs, or DDGs. Then, the size of the giant component was plotted over increasing radius $\epsilon$. In **C)**, in the left panel counts per million (cpm) then z-score normalization was performed on 10x A20 and NIH3T3 cell line data. On the right panel z-score normalization then counts per million normalization was performed on 10x A20 and NIH3T3 cell line. Then, the size of the giant component was plotted over increasing radius $\epsilon$. In **D)**, in the left panel counts per million (cpm) then z-score normalization was performed on MERFISH Mouse brain data. On the right panel z-score normalization then counts per million normalization was performed on MERFISH Mouse brain data. Then, the size of the giant component was plotted over increasing radius $\epsilon$.

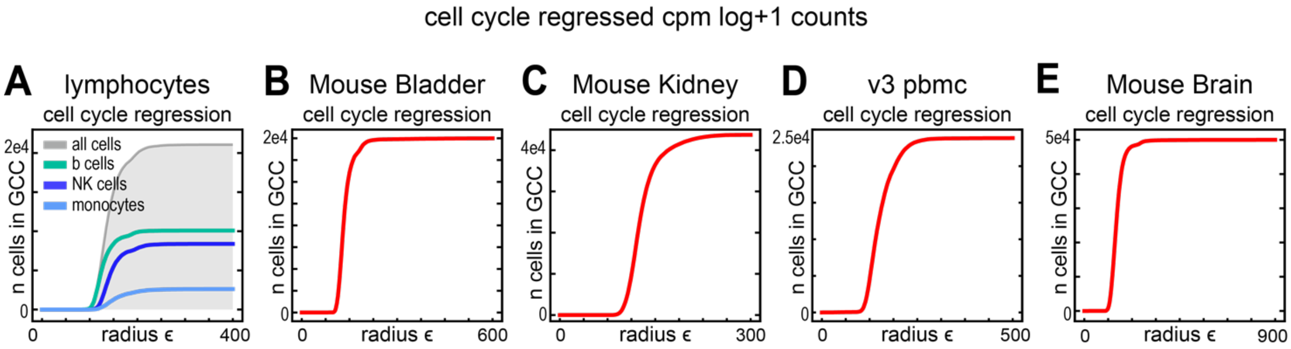


**Figure S5.17.** $\epsilon$ network analysis of data normalized using Scanpy's cell cycle regression function. For each panel $\epsilon$ networks were constructed after cell cycle regression was applied to cpm log+1 normalized data. Then, the size of the giant component was plotted over increasing radius $\epsilon$, when the $\epsilon$ networks were constructed cell cycle regression was applied to **A)** 10x Genomics scRNA-seq data from the FACS purified lymphocytes, or 10x Genomics scRNA-seq data from **B)** the Mouse bladder, **C)** the Mouse kidney, **D)** PBMC data collected with version 3 chemistry, **E)** data from the Mouse brain.

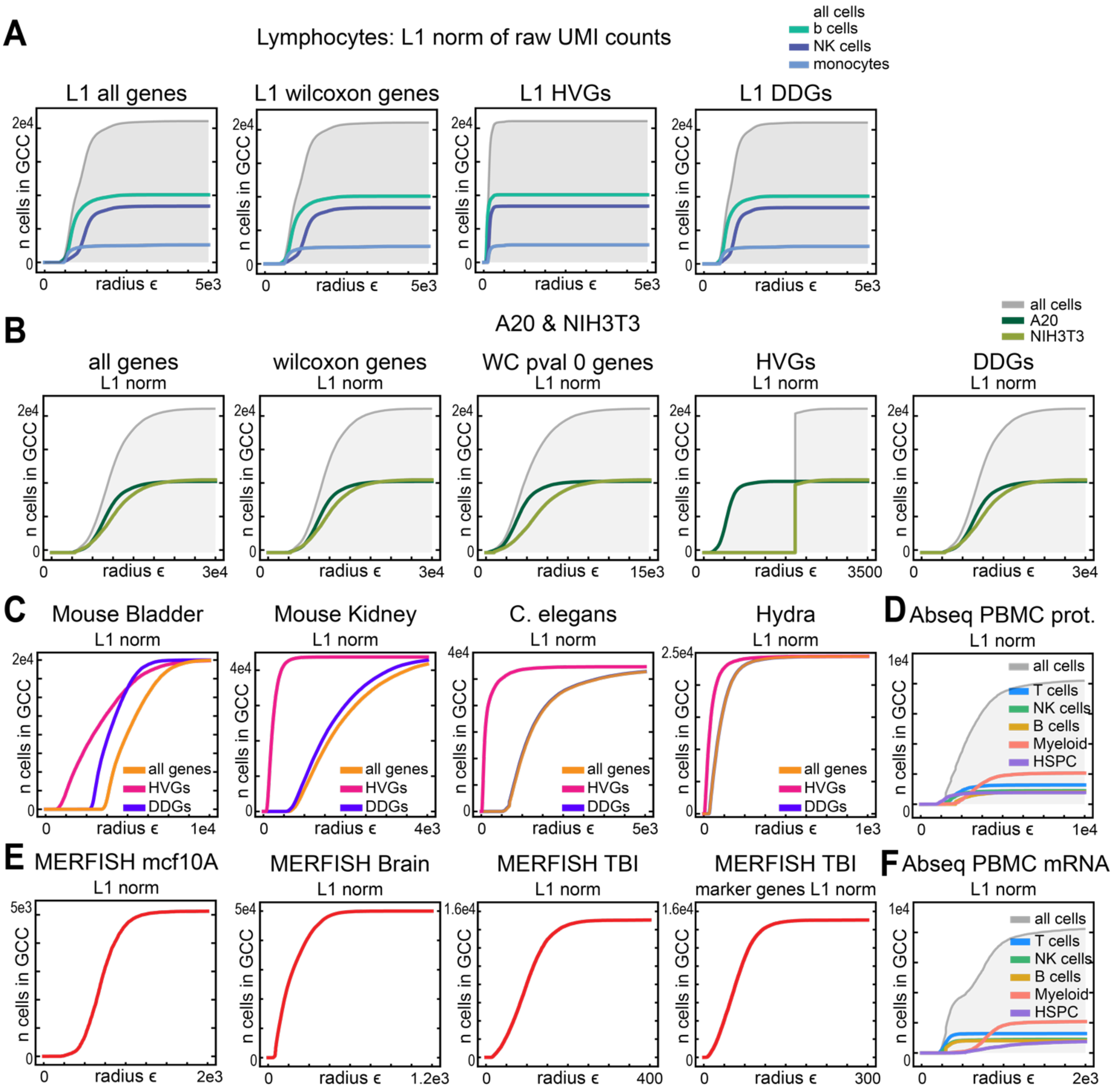


**Figure S5.18.** $\epsilon$ network analysis of data using the $\ell^1$ norm. For each panel $\epsilon$ networks were constructed using the $\ell^1$ measure of distance. Then, the size of the giant component was plotted over increasing radius $\epsilon$, when the $\epsilon$ networks were constructed using either all genes, Wilcoxon genes, Highly variable genes (HVGs), or Differentially distributed genes (DDGs) using the 10x Genomics scRNA-seq data from **A)** the FACs sorted lymphocytes, **B)** A20 & NIH3T3 cell lines, **C)** the Mouse bladder, **D)** the Mouse kidney, **E)** C. elegans embryo, or **F)** the Hydra. Giant component analysis was performed on $\epsilon$ networks constructed using the $\ell^1$ norm using **G)** ABseq PBMC protein counts, the MERFISH genes from **H)** mcf10A cells, **I)** Mouse brain, **J)** Mouse brain traumatic brain injury data, **K)** marker genes from MERFISH traumatic brain injury data, or **L)** ABseq PBMC mRNA counts.

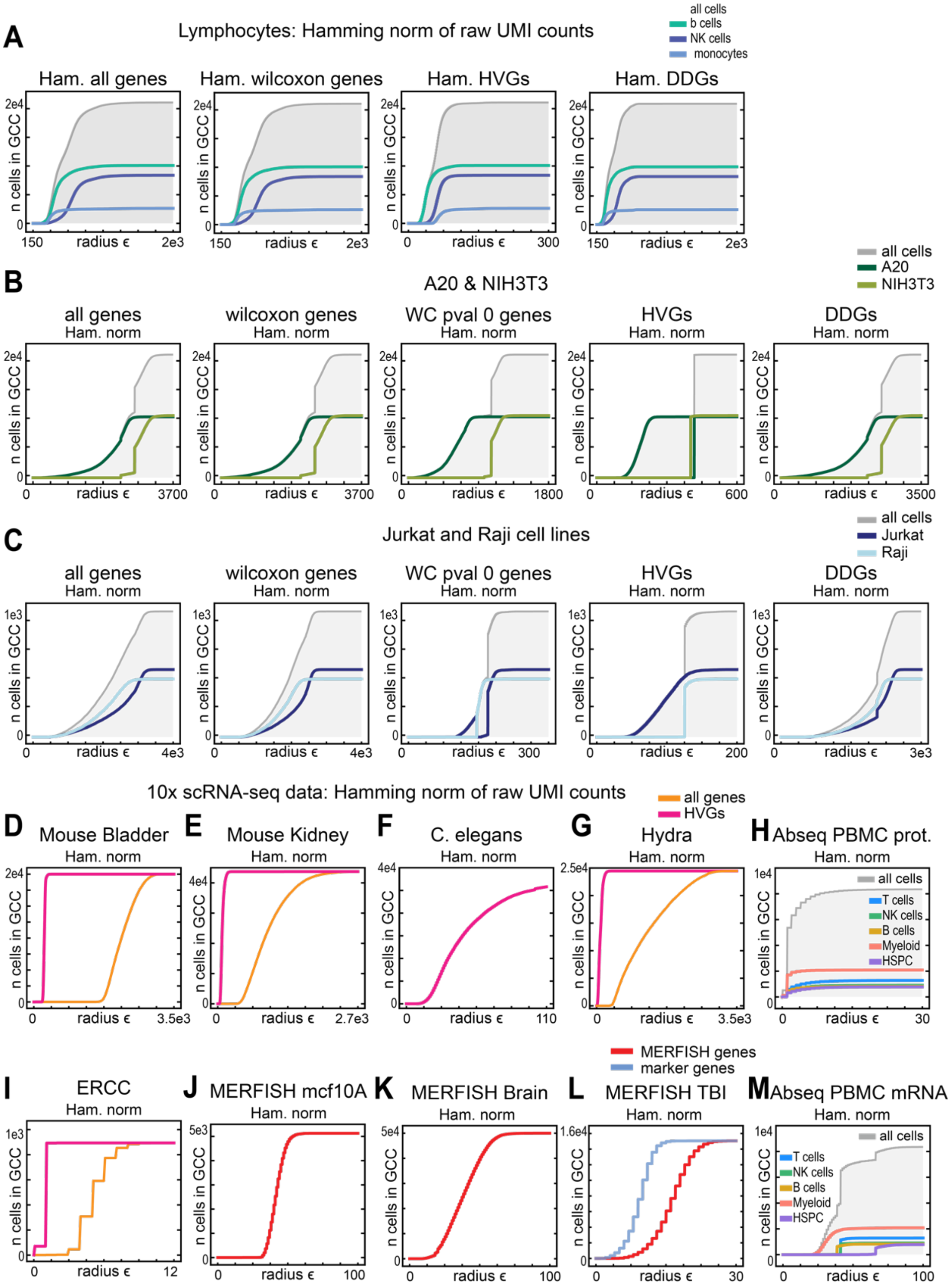


**Figure S5.19.** $\epsilon$ network analysis of data using the hamming norm. For each panel $\epsilon$ networks were constructed using the Hamming notion of distance. To do this, every cell's gene expression feature vector was converted into a vector of 0s or 1s: a gene is "0" if that gene is not expressed in that cell, or "1" if it is expressed in that cell. The Hamming distance is the $\ell^1$ distance on these resulting 0/1 vectors, and simply quantifies the number of differences between them (i.e. the number of times one cell has a "1" and the other a "0" for a gene or *vice versa*). The size of the giant component was plotted over increasing radius $\epsilon$, where the $\epsilon$ networks were constructed using either all genes, Wilcoxon genes, Highly Variable Benes (HVGs), or Differentially Distributed Genes (DDGs) using the 10x Genomics scRNA-seq data from **A)** the FACs sorted lymphocytes, **B)** A20 & NIH3T3 cell lines, **C)** Jurkat & Raji cell lines, **D)** the Mouse bladder, **E)** the Mouse kidney, **F)** C. elegans embryo, or **G)** the Hydra. Giant component analysis was performed on $\epsilon$ networks constructed using the $\ell^1$ norm using **H)** ABseq PBMC protein counts, the MERFISH genes from **I)** mcf10A cells, **J)** Mouse brain, **K)** Mouse brain traumatic brain injury data, **L)** marker genes from MERFISH traumatic brain injury data, or **M)** ABseq PBMC mRNA counts. Note that, due to its definition, the Hamming distance can only take certain discrete values (0 genes differ between cells, 1 gene differs, 2 genes, etc.). This is the reason for the regular step-like behaviors observed in panels (F) and (L) (and to a lesser extent in panels H and M), and is not indicative of separation between cell type groups.

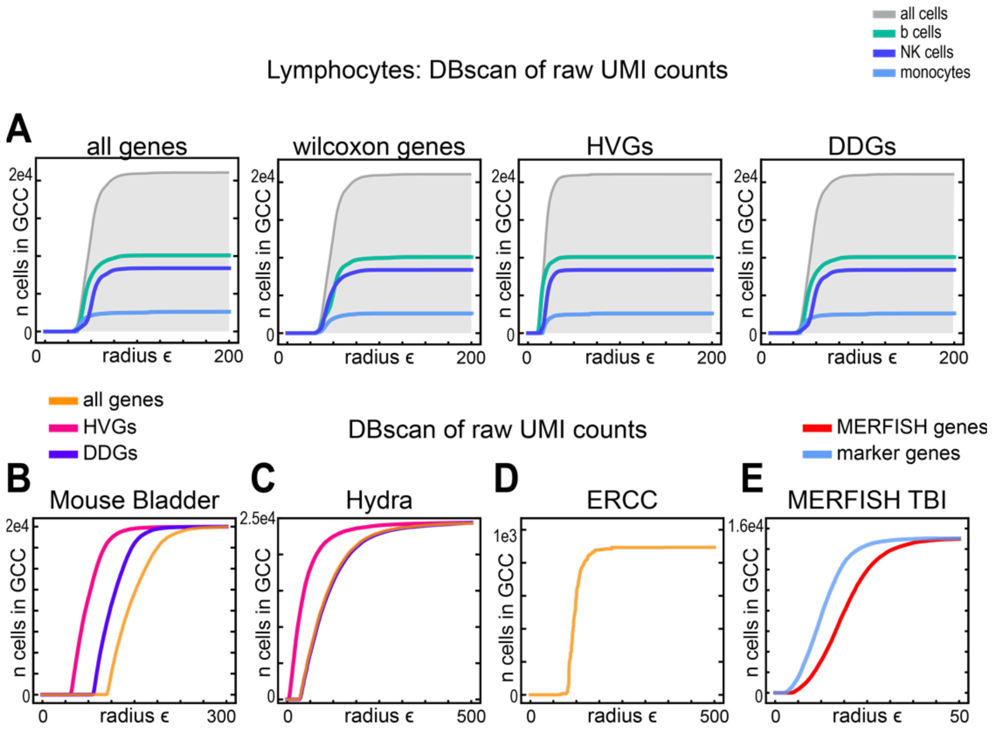


**Figure S5.20.** Giant component analysis of data using the DB scan clustering method. For each panel $\epsilon$ networks were constructed using a non-single linkage method of clustering, namely DB scan. Then, the size of the giant component was plotted over increasing radius $\epsilon$, when the $\epsilon$ networks were constructed using either all genes, Wilcoxon genes, Highly Variable Genes (HVGs), or Differentially Distributed Genes (DDGs) for the **A)** 10x Genomics scRNA-seq data from the FACs sorted lymphocytes, or 10x Genomics scRNA-seq data from **B)** the Mouse bladder, **C)** Hydra, **F)** ERCC spike in data, or **E)** MERFISH data from mouse brain traumatic injury (TBI) data using all MERFISH genes or only marker genes.

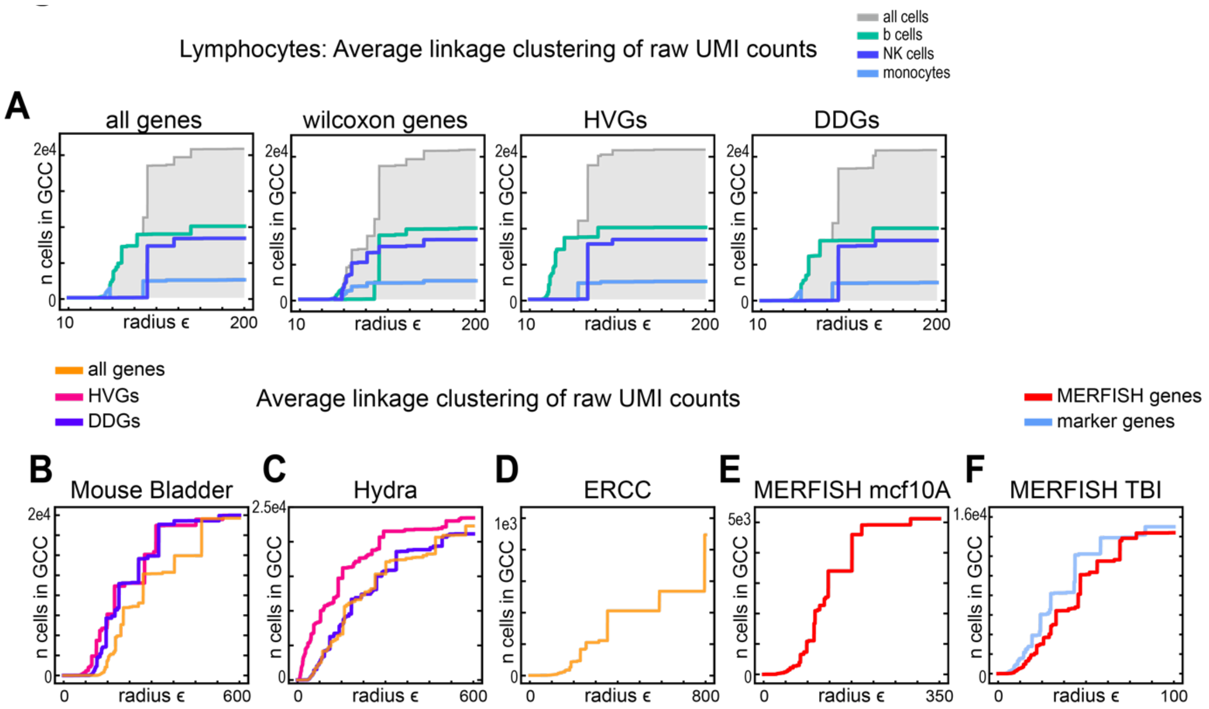


**Figure S5.21.** Giant component analysis of data using the average linkage clustering method. For each panel $\epsilon$ networks were constructed using a non-single linkage method of construction, where the average linkage was considered for each node. Then, the size of the giant component was plotted over increasing radius $\epsilon$, when the $\epsilon$ networks were constructed using either all genes, Wilcoxon genes, Highly variable genes (HVGs), or Differentially distributed genes (DDGs) for the **A)** 10x Genomics scRNA-seq data from the FACs sorted lymphocytes, or 10x Genomics scRNA-seq data from **B)** the Mouse bladder, **C)** Hydra, **F)** ERCC spike in data, or MERFISH data from **E)** the mcf10A cell line, or **F)** mouse brain traumatic injury (TBI) data using all MERFISH genes or only marker genes.

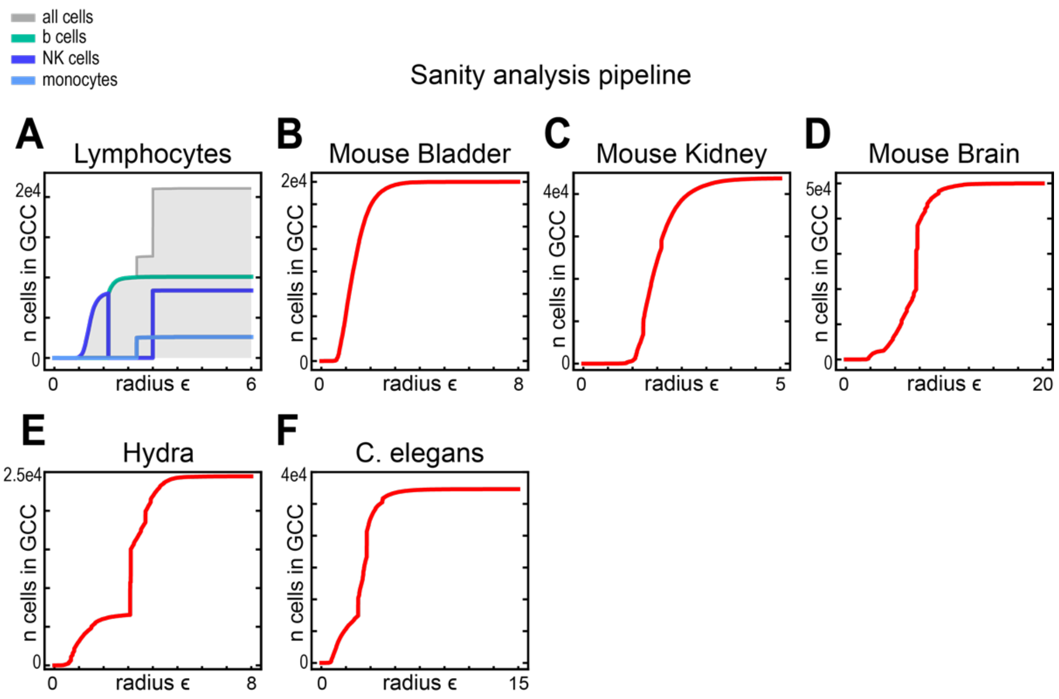


**Figure S5.22.** $\epsilon$ network analysis of data normalized using the Sanity analysis pipeline. For each panel $\epsilon$ networks were constructed using the distance matrix produced by Sanity. Then, the size of the giant component was plotted over increasing radius $\epsilon$, when the $\epsilon$ networks were constructed using **A)** 10x Genomics scRNA-seq data from the FACS purified lymphocytes, or 10x Genomics scRNA-seq data from **B)** the Mouse bladder, **C)** the Mouse kidney, **D)** the Mouse Brain, **E)** the Hydra, or **F)** C. elegans embryos.

### Figure S6.1

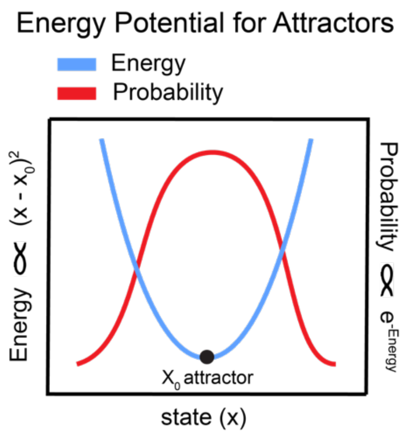


**Figure S6.1.** Simple schematic for the relationship between probability density and the shape of a "potential well" near an attractor. Here we have a potential well with a quadratic deviation around the center of the attractor, which should lead to a well-localized distribution of points near the center of the attractor.

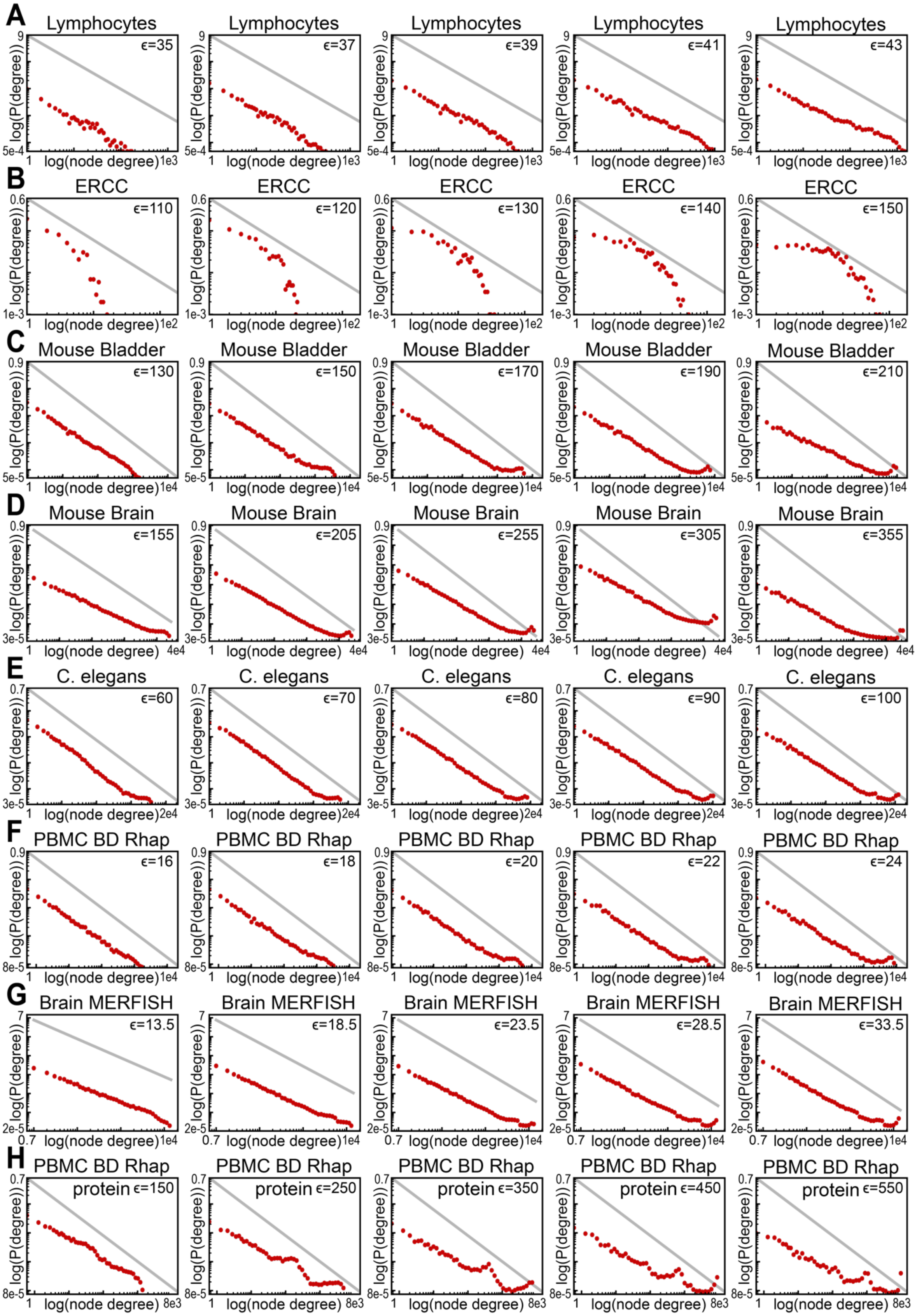


**Figure S6.2.** Density distributions of single-cell datasets are consistent across values of $\epsilon$. **A-H)** Degree Distribution for the $\epsilon$-network of various datasets at the indicated $\epsilon$: **A)** FACS-purified Lymphocytes **B)** ERCC control data **C)** Mouse Bladder, **D)** Mouse Brain, **E)** *C. elegans,* **F)** BD Rhapsody PBMC genes, **G)** MERFHISH Brain, **H)** BD Rhapsody PBMC protein. These data were collected on the 10x platform, unless indicated otherwise. The gray line represents a power-law with an exponent of -1 for reference.

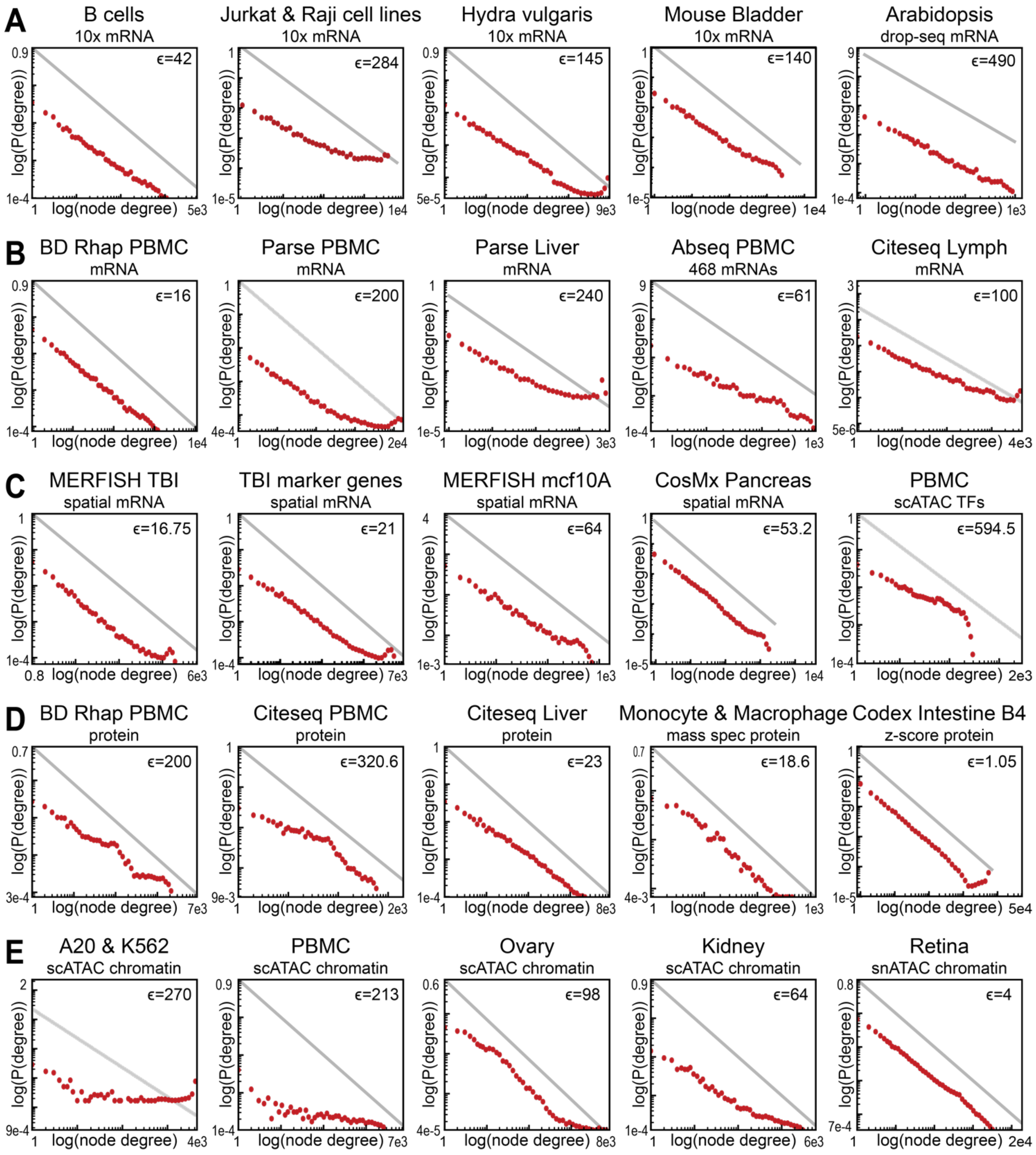


**Figure S6.3.** Density distributions from different single-cell measurement platforms indicate the absence of attractor structures. **A-E)** Degree Distribution for the $\epsilon$-network of various datasets at the indicated $\epsilon$ for data generated using targeted and genome **A)** droplet-based scRNA-seq platforms (10x Genomics and drop-seq), **B)** other RNA-seq platforms, **C)** spatial transcriptomics platforms, **D)** single-cell proteomics platforms, and **E)** single-cell chromatin accessibility platforms. The gray line represents a power-law with an exponent of -1 for reference. Additional details of each dataset are available in Table S1.

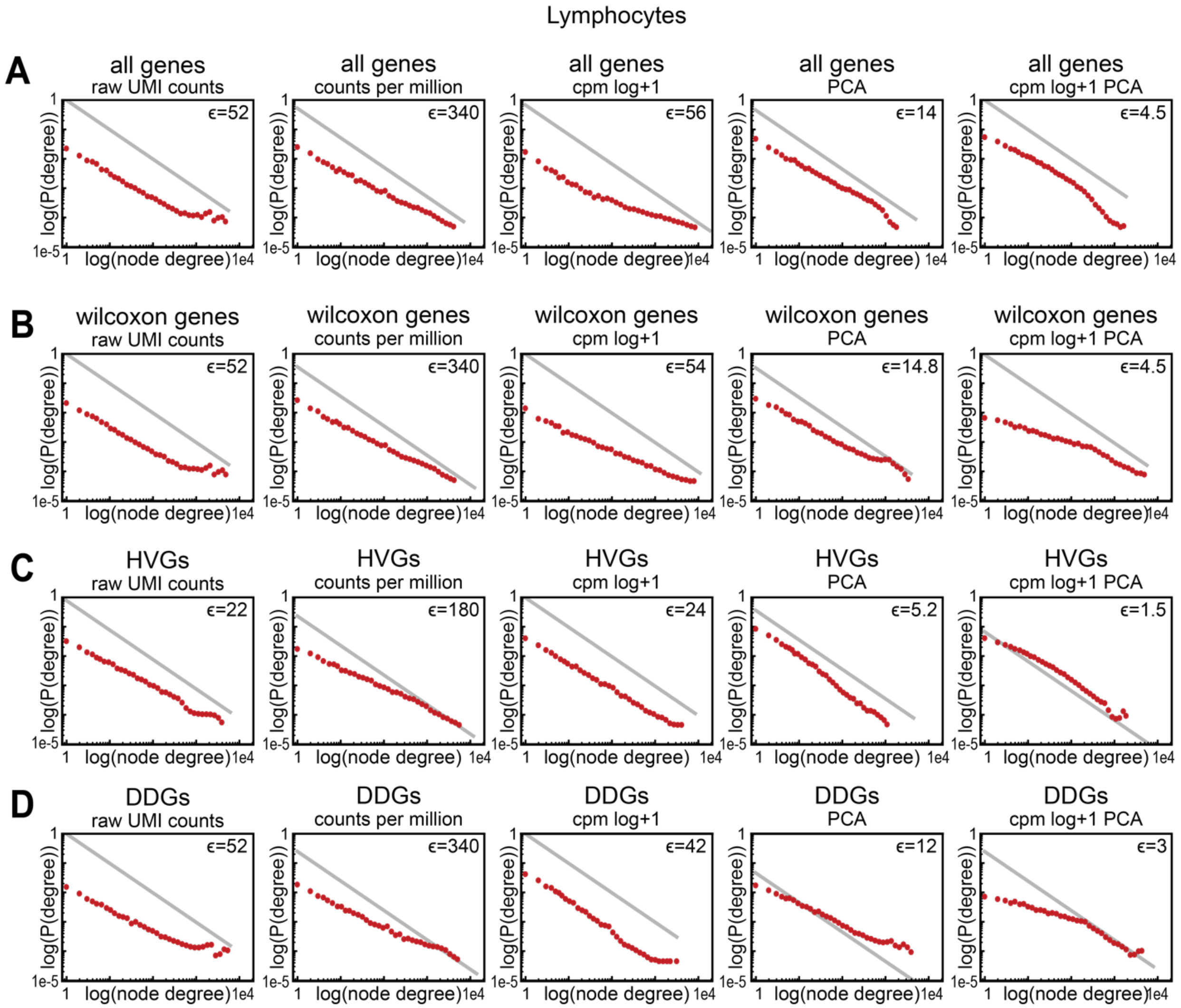


**Figure S6.4.** Nonlinear transformations from the scRNA-seq analysis pipeline do not produce gaussian-like density distributions in lymphocyte data. **A-D)** Degree distributions for the $\epsilon$ networks of the 10x Genomics FACS-purified lymphocytes after feature selection and various nonlinear transformations. Panels from left to right show degree distributions at the indicated $\epsilon$ after no transformation, counts per million (CPM) normalization, counts per million normalization then log+1 transformation, principal component analysis (PCA), or CPM, then log+1, then PCA transformed data. These transformations were performed on the lymphocyte data using either **A)** all genes, **B)** Wilcoxon rank sum genes, **C)** Highly Variable Genes (HVGs), or **D)** Differentially Distributed Genes (DDGs). The gray line represents a power-law with an exponent of -1 for reference.

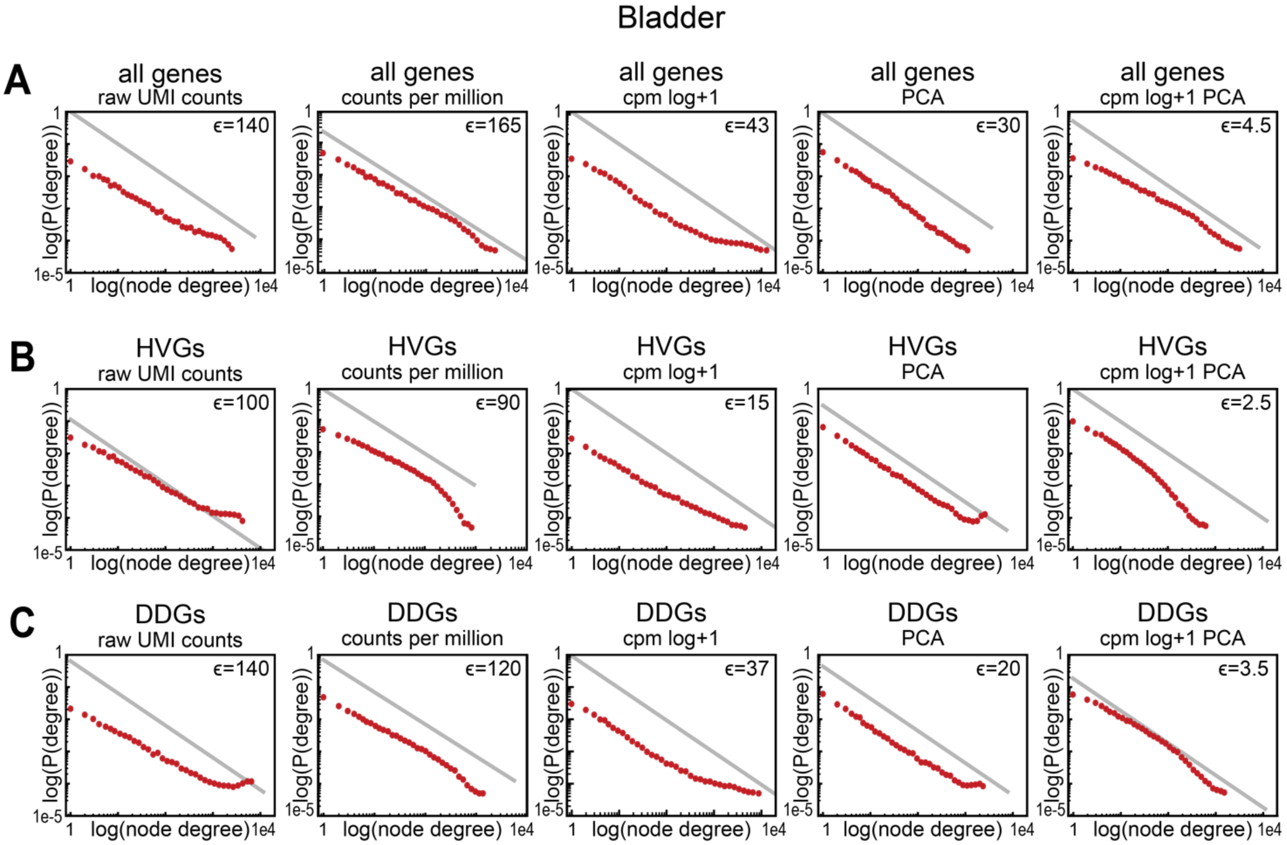


**Figure S6.5.** Nonlinear transformations from the scRNA-seq analysis pipeline do not produce gaussian-like density distributions in Bladder data. **A-C)** Degree distributions for the $\epsilon$ networks of 10x Genomics scRNA-seq data from the mouse bladder after feature selection and various nonlinear transformations. Panels from left to right show degree distributions at the indicated $\epsilon$ after no transformation, counts per million (CPM) normalization, counts per million normalization then log+1 transformation, principal component analysis (PCA), or CPM, then log+1, then PCA transformed data. These transformations were performed on the lymphocyte data using either **A)** all genes, **B)** Highly Variable Genes (HVGs), or **C)** Differentially Distributed Genes (DDGs). The gray line represents a power-law with an exponent of -1 for reference.

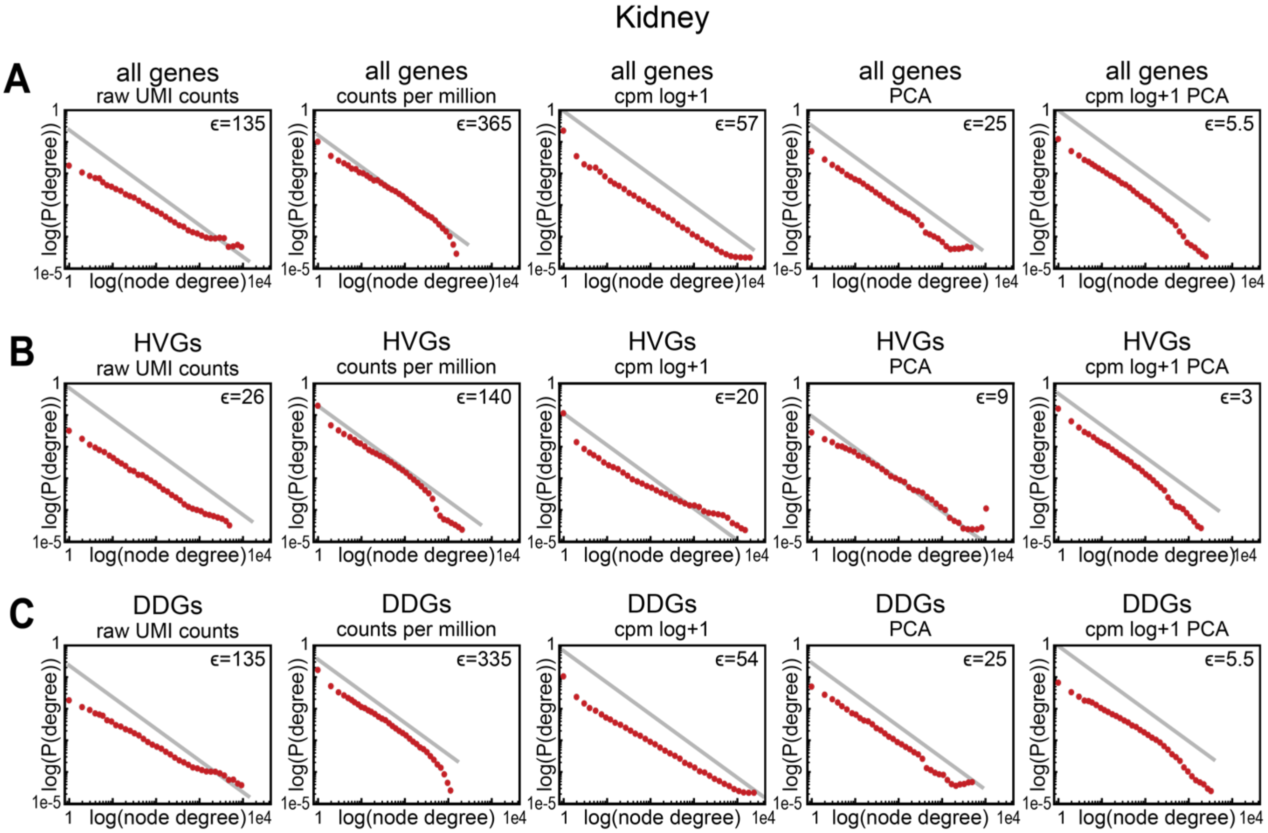


**Figure S6.6.** Nonlinear transformations from the scRNA-seq analysis pipeline do not produce gaussian-like density distributions in Kidney data. **A-C)** Degree distributions for the $\epsilon$ networks of 10x Genomics scRNA-seq data from the mouse kidney after feature selection and various nonlinear transformations. Panels from left to right show degree distributions at the indicated $\epsilon$ after no transformation, counts per million (CPM) normalization, counts per million normalization then log+1 transformation, principal component analysis (PCA), or CPM, then log+1, then PCA transformed data. These transformations were performed on the lymphocyte data using either **A)** all genes, **B)** Highly Variable Genes (HVGs), or **C)** Differentially Distributed Genes (DDGs). The gray line represents a power-law with an exponent of -1 for reference.

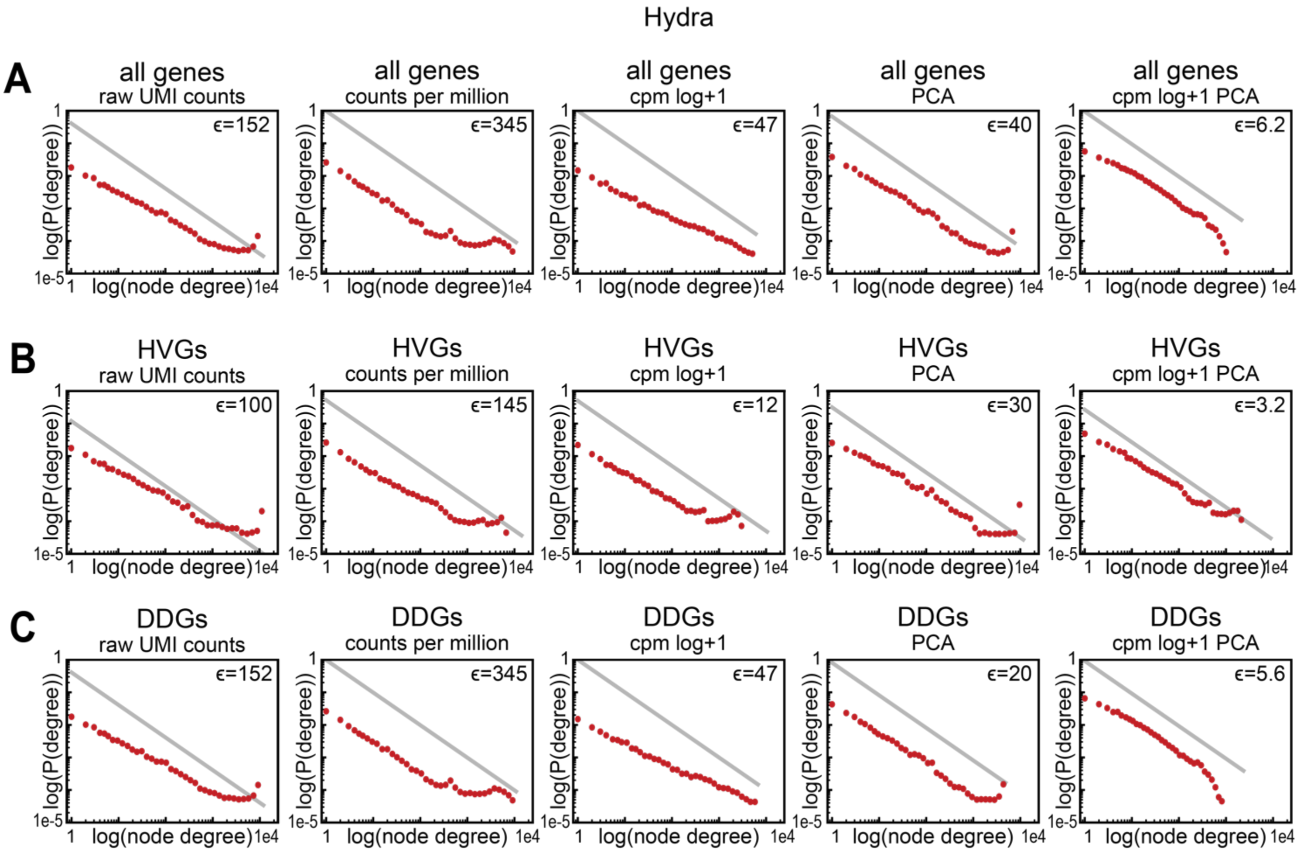


**Figure S6.7.** Nonlinear transformations from the scRNA-seq analysis pipeline do not produce gaussian-like density distributions in Hydra data. **A-C)** Degree distributions for the $\epsilon$ networks of 10x Genomics scRNA-seq data from *Hydra vulgaris* after feature selection and various nonlinear transformations. Panels from left to right show degree distributions at the indicated $\epsilon$ after no transformation, counts per million (CPM) normalization, counts per million normalization then log+1 transformation, principal component analysis (PCA), or CPM, then log+1, then PCA transformed data. These transformations were performed on the lymphocyte data using either **A)** all genes, **B)** Highly Variable Genes (HVGs), or **C)** Differentially Distributed Genes (DDGs). The gray line represents a power-law with an exponent of -1 for reference.

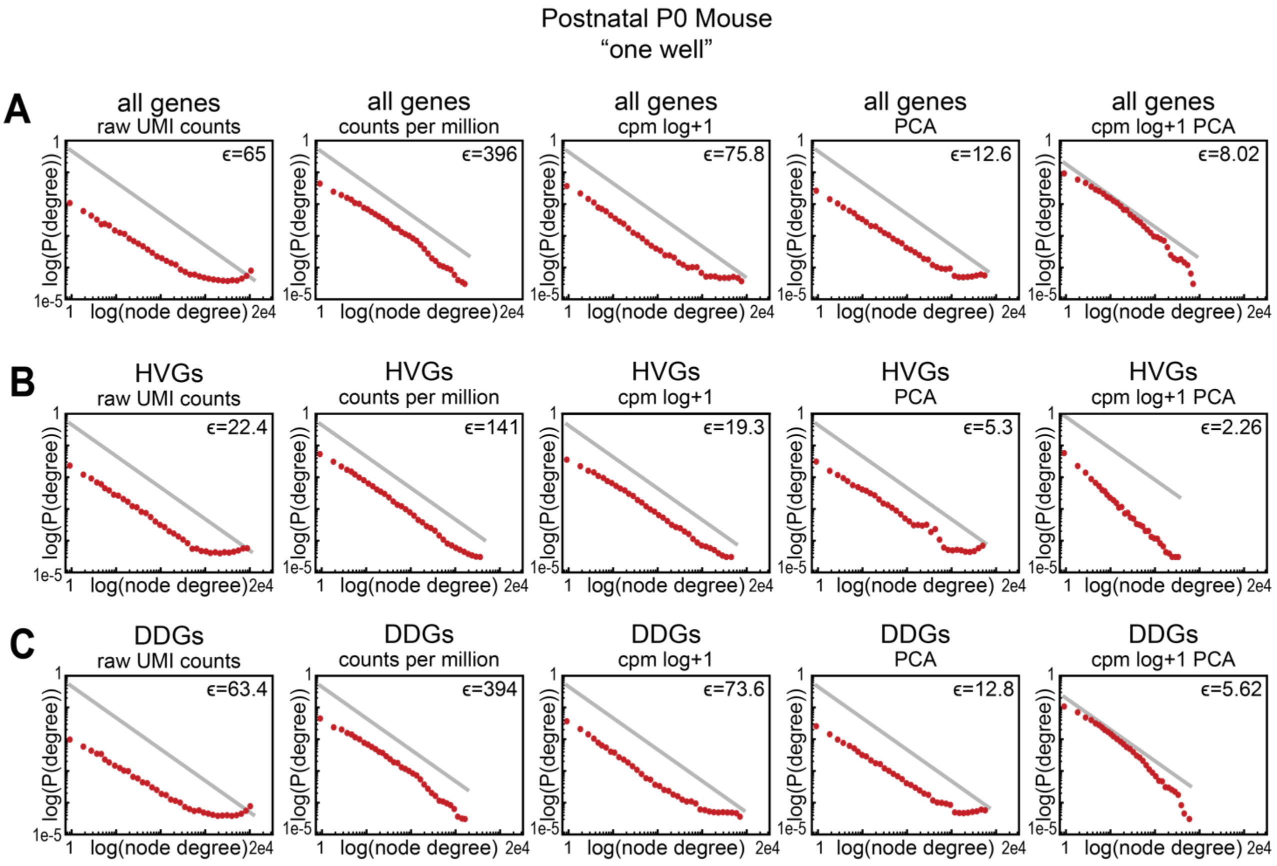


**Figure S6.8.** Nonlinear transformations from the scRNA-seq analysis pipeline do not produce gaussian-like density distributions in Postnatal P0 Mouse data. **A-C)** Degree distributions for the $\epsilon$ networks of 10x Genomics scRNA-seq data from one well and one embryo of the Mouse atlas data, after feature selection and various nonlinear transformations. Panels from left to right show degree distributions at the indicated $\epsilon$ after no transformation, counts per million (CPM) normalization, counts per million normalization then log+1 transformation, principal component analysis (PCA), or CPM, then log+1, then PCA transformed data. These transformations were performed on the lymphocyte data using either **A)** all genes, **B)** Highly Variable Genes (HVGs), or **C)** Differentially Distributed Genes (DDGs). The gray line represents a power-law with an exponent of -1 for reference.

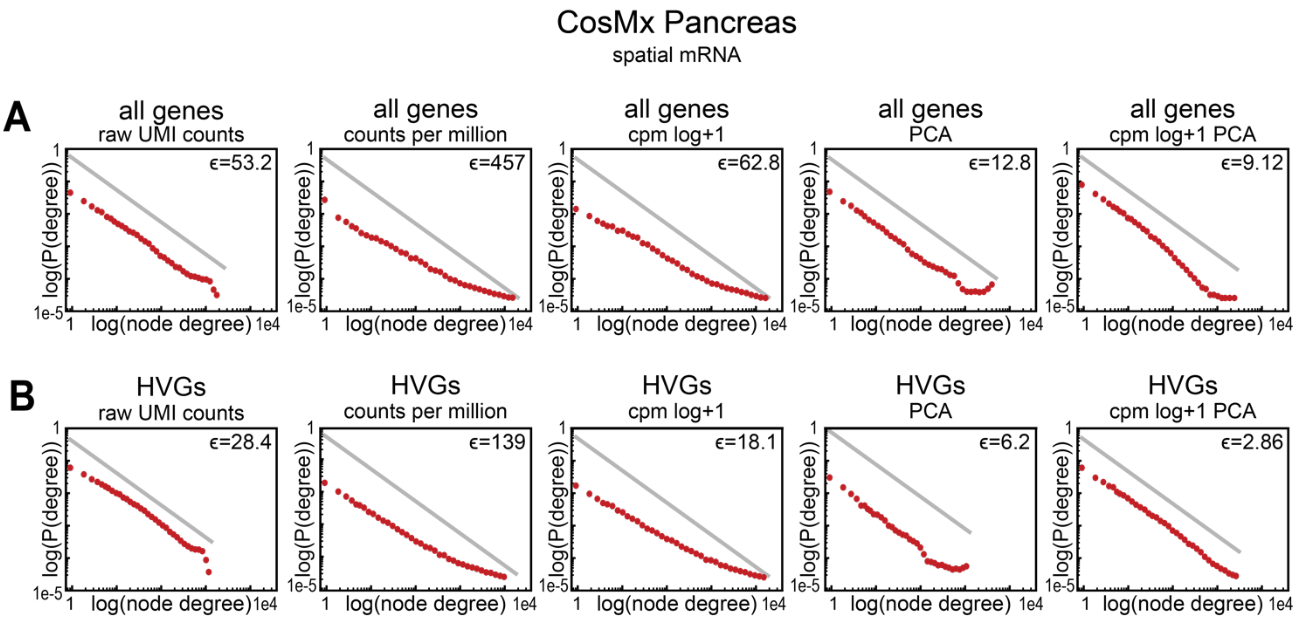


**Figure S6.9.** Nonlinear transformations from the scRNA-seq analysis pipeline do not produce gaussian-like density distributions in CosMx Pancreas data. **A-C)** Degree distributions for the $\epsilon$ networks of spatial scRNA data from CosMx Pancreas data after feature selection and various nonlinear transformations. Panels from left to right show degree distributions at the indicated $\epsilon$ after no transformation, counts per million (CPM) normalization, counts per million normalization then log+1 transformation, principal component analysis (PCA), or CPM, then log+1, then PCA transformed data. These transformations were performed on the lymphocyte data using either **A)** all genes, **B)** Highly Variable Genes (HVGs), or **C)** Differentially Distributed Genes (DDGs). The gray line represents a power-law with an exponent of -1 for reference.

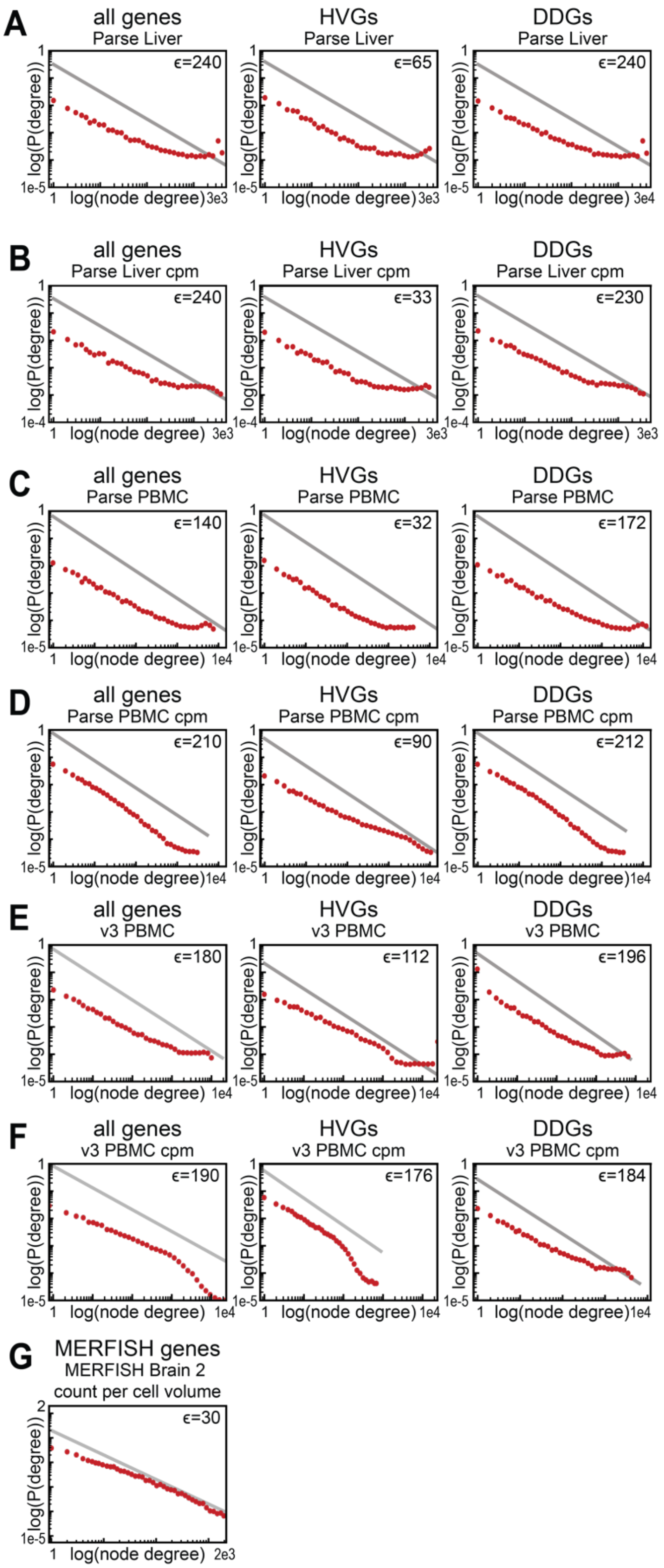


**Figure S6.10.** Feature selection and cell volume normalization do not produce gaussian-like density distributions in data from different single-cell measurement platforms. **A-F)** Degree distributions at the indicated $\epsilon$ for the $\epsilon$ networks of different feature-selected scRNA-seq data. The leftmost panel shows degree distributions from $\epsilon$ networks constructed using all genes, the middle panel shows degree distributions from the subset of Highly Variable Genes (HVGs), and the rightmost panel shows degree distributions from the subset of Differentially Distributed Genes (DDGs). **A)** Non-transformed and **B)** counts per million (CPM) transformed scRNA-seq data collected from mouse liver using the Parse Bioscience platform. **C)** Non-transformed and **D)** counts per million (CPM) transformed scRNA-seq data collected from mouse PBMCs using the Parse Bioscience platform. **E)** Non-transformed and **F)** counts per million (CPM) transformed scRNA-seq data collected from mouse PBMCs using the 10x Genomics version 3 chemistry. **G)** Degree distributions at the indicated $\epsilon$ for $\epsilon$ networks constructed from normalized MERFISH Brain data. Here, each gene was counted by MERFISH then divided by the volume of the cell. (A-G) The gray line represents a power-law with an exponent of -1 for reference.

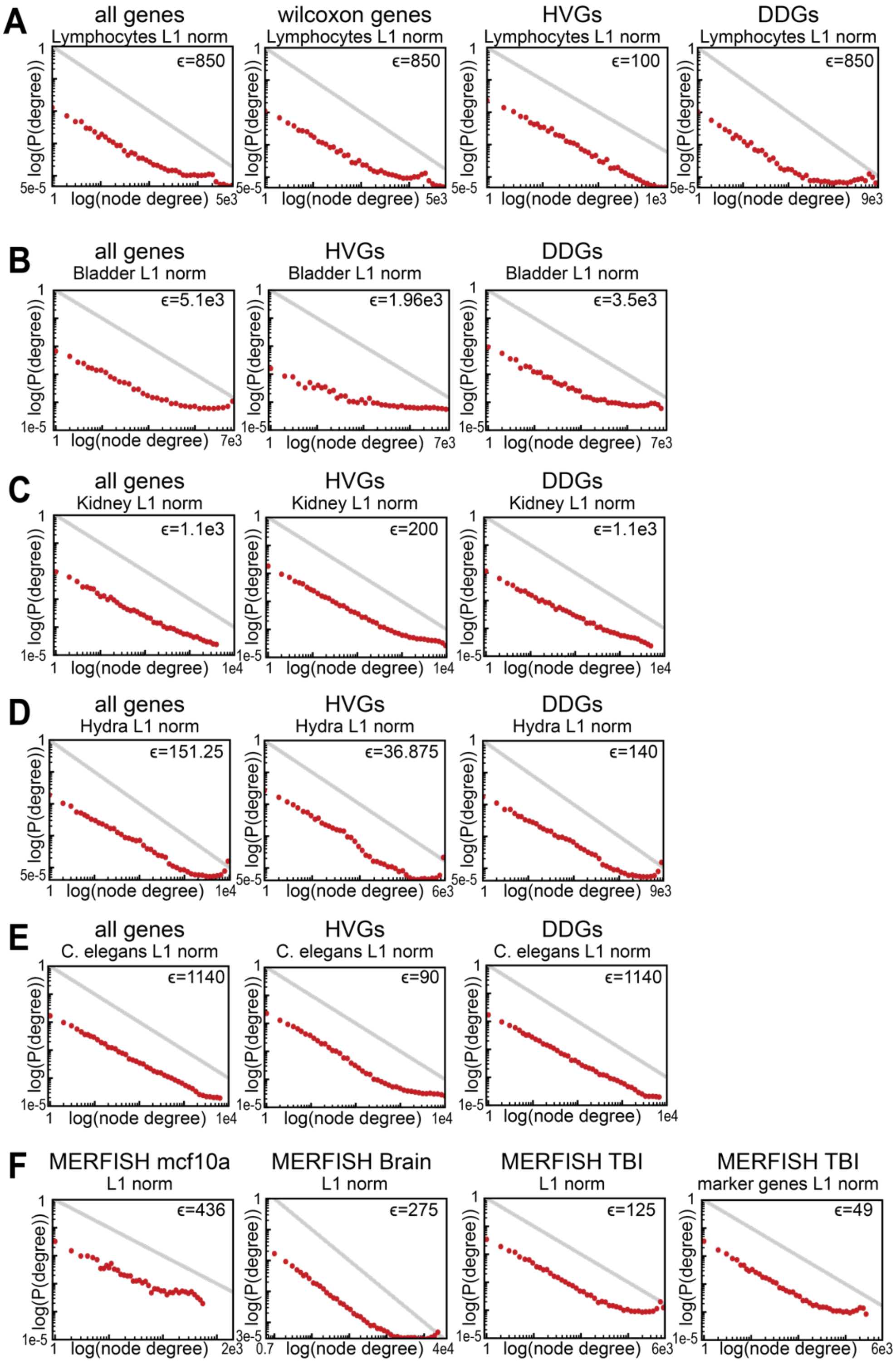


**Figure S6.11.** The $\ell^1$ norm does not produce gaussian-like density distributions in data from different single-cell measurement platforms. **A-E)** Degree distributions at the indicated $\epsilon$ for the $\epsilon$ networks constructed using the $\ell^1$ norm of 10x Genomics scRNA-seq data from **A)** FACS-purified lymphocytes, **B)** mouse bladder, **C)** mouse kidney, **D)** *Hydra vulgaris,* and **E)** *C. elegans*. The leftmost panel shows degree distributions from $\epsilon$ networks constructed using all genes, the middle panel shows degree distributions from the subset of Highly Variable Genes (HVGs), and the rightmost panel shows degree distributions from the subset of Differentially Distributed Genes (DDGs). **F)** Degree distributions at the indicated $\epsilon$ for the $\epsilon$ networks constructed using the $\ell^1$ norm of MERFISH data from mcf10a cells, mouse brain, traumatic brain injury data for all MERFISH genes, or traumatic brain injury data for only marker genes. The gray line represents a power-law with an exponent of -1 for reference.

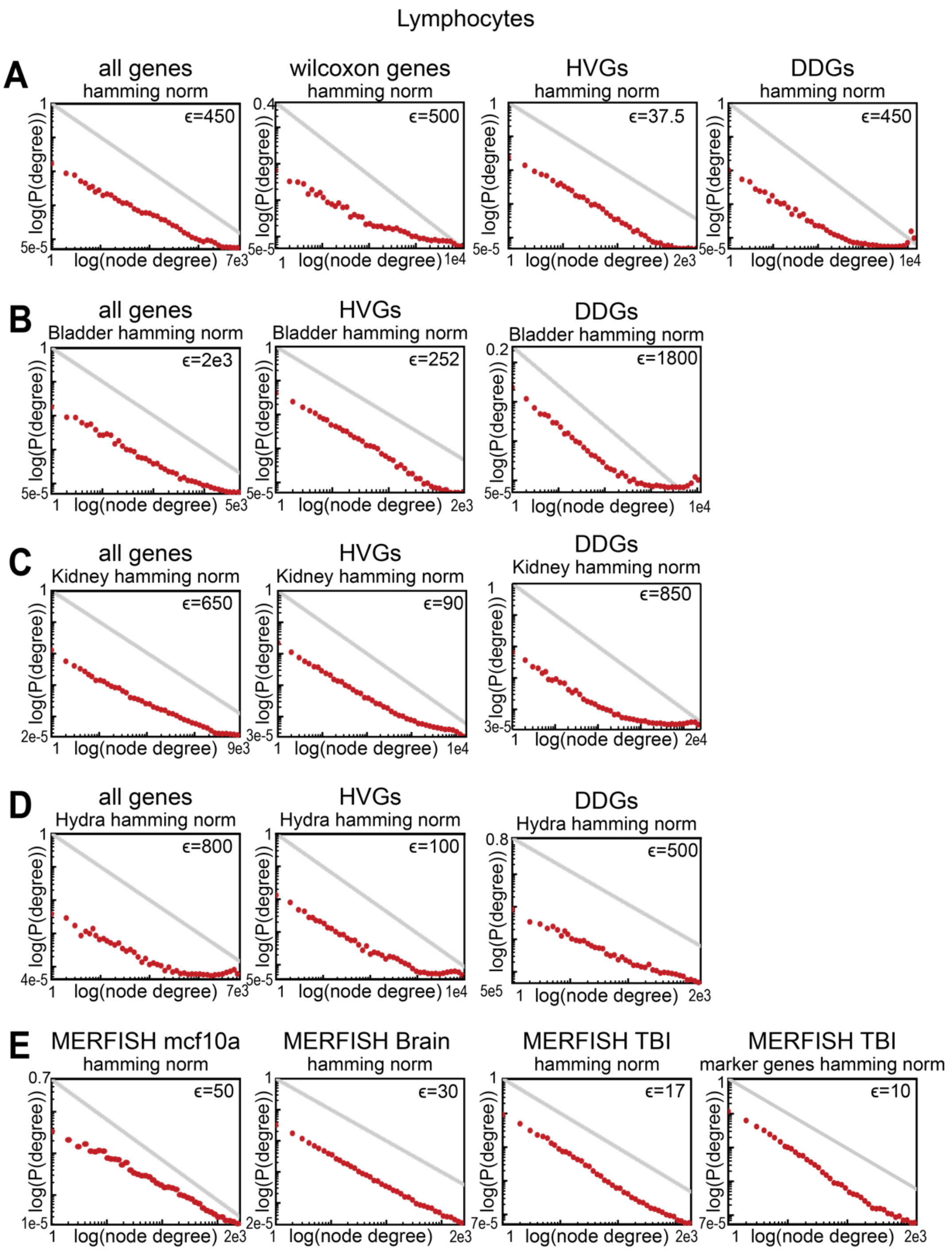


**Figure S6.12.** The Hamming norm does not produce gaussian-like density distributions in data from different single-cell measurement platforms. In this case all Hamming distances were calculated as described in Fig. S5.13. **A-E)** Degree distributions at the indicated $\epsilon$ for the $\epsilon$ networks constructed using the Hamming norm of 10x Genomics scRNA-seq data from **A)** FACS-purified lymphocytes, **B)** mouse bladder, **C)** mouse kidney, **D)** *Hydra vulgaris,* and **E)** *C. elegans*. The leftmost panel shows degree distributions from $\epsilon$ networks constructed using all genes, the middle panel shows degree distributions from the subset of Highly Variable Genes (HVGs), and the rightmost panel shows degree distributions from the subset of Differentially Distributed Genes (DDGs). **F)** Degree distributions at the indicated $\epsilon$ for the $\epsilon$ networks constructed using the Hamming norm of MERFISH data from mcf10a cells, mouse brain, traumatic brain injury data for all MERFISH genes, or traumatic brain injury data for only marker genes. The gray line represents a power-law with an exponent of -1 for reference.

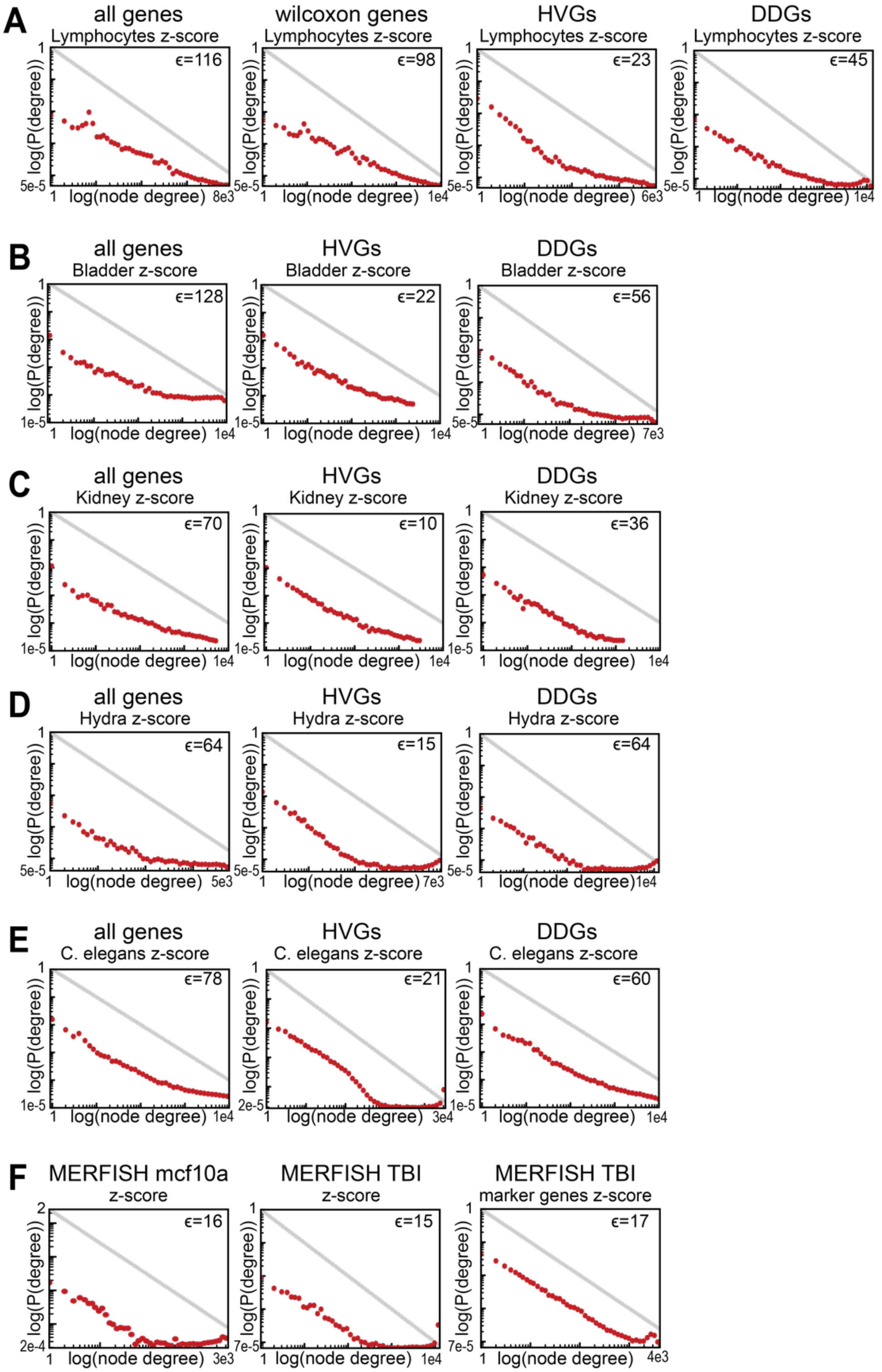


**Figure S6.13.** Z-score transformation does not produce gaussian-like density distributions in data from different single-cell measurement platforms. **A-E)** Degree distributions at the indicated $\epsilon$ for the $\epsilon$ networks of feature selected and z-score transformed data for 10x Genomics scRNA-seq data from **A)** FACS-purified lymphocytes, **B)** mouse bladder, **C)** mouse kidney, **D)** *Hydra vulgaris,* and **E)** *C. elegans*. The leftmost panel shows degree distributions from $\epsilon$ networks constructed using all genes, the middle panel shows degree distributions from the subset of Highly Variable Genes (HVGs), and the rightmost panel shows degree distributions from the subset of Differentially Distributed Genes (DDGs). **F)** Degree distributions at the indicated $\epsilon$ for the $\epsilon$ networks of z-score transformed MERFISH data from mcf10a cells, traumatic brain injury data for all MERFISH genes, or traumatic brain injury data for only marker genes. The gray line represents a power-law with an exponent of -1 for reference.
