## Supplemental Text for "A lack of distinct cell identities in single-cell measurements: revisiting Waddington’s landscape"

**<u>Section 1: Titrating cell types from the mouse atlas</u>**

As discussed in the text, we demonstrate that our $\epsilon$ networks can separate groups of cells in extremely high-dimensional and noisy data generated from biological samples. Interestingly, we are only able to observe this separation in a small number of datasets when a non-linear, counts-per-million transformation is applied to the raw count data. Nonetheless, the fact that we only observe this cell-type separation in data generated from cell lines or data that has been FACS sorted for cell types, prompts the question as to why there exists an absence of separation in the vast majority of biological data characterized from tissues, organs, and organisms. We hypothesized that the absence of separation in more complex biological samples arises in part due to the representation of the full biological heterogeneity that exists across cell types within a tissue. We reasoned that this could be explained by a scenario where tissue samples contain cells of distinct types that exist within slightly overlapping regions of gene expression space, or a scenario where tissue samples contain cells of different types that occupy completely enmeshed regions of gene expression space.

To address this, we designed an experiment to characterize the structure of epigenetic heterogeneity in complex multicellular samples using data from the Mouse Cell Atlases from the Sanger Institute. We first isolated data from a single embryo of the Mouse atlas data, selecting the latest stage in development available, which was 0 days after birth or "P0". Next, to avoid artefacts potentially introduced by batch effects we used the cell-barcode prefixes to select ~30,000 cells from the same individual embryo with the same set of cell-barcode prefixes. All subsequent experiments described below were performed on data selected from this single P0 embryo, from "one well" of the Mouse atlas data.

We next used the available cell-type metadata from the atlas database to select specific subsets of cells. The cell-type annotations were generated by the Sanger Institute after batch correction across all of the developmental samples collected, followed by the standard set of scRNA-seq transformations, two rounds of clustering, and then marker gene annotation to assign cell-type classes to the clusters. We first isolated the distinct lineages of fibroblasts and GABAergic neurons. When we applied our $\epsilon$ network characterization to the raw UMI counts, we observe no separation between the two distinct cell types. When we normalize the data using a counts per million transformation, we are able to observe that about 2/3 of the GABAergic neurons exist in a distinct region from fibroblast cells (Fig S3.4A). However, about 1/3 of GABAergic neurons are as close to fibroblasts as they are to other GABAergic neurons, suggesting that these two cell-type groups don't exist in robustly separable groups even at this level.

We next tested whether adding back cell types representing more closely related populations would reduce the ability to detect some degree of separation between the GABAergic neuron and fibroblast subsets. Since cell-types in the atlas data were hierarchically defined using two rounds of clustering, we selected the larger superset or the more coarse-grained cell type cluster in which these specific cell types have membership. That is, GABAergic neurons belong to a larger superset of cell types labeled "CNS neurons", while fibroblasts belong to a larger superset labeled "Mesoderm" cells. We applied the counts-per-million normalization to the "CNS neuron" and "Mesoderm" supersets then performed our $\epsilon$ network characterization. We find that, when we include the more closely related cell-types in the graph, only about $1/10^{th}$ of "CNS neurons" exist in a region of gene expression space that is separate from "Mesoderm" cell types (Fig S3.4A). When we apply the $\epsilon$ network characterization to all cell type classes from the P0 embryo, we fail to observe any separability between GABAergic neurons and fibroblasts (Fig S3.4A, last panel). These results confirm that if we subset the data into two more distinct regions of gene expression space, while we may be able to observe partial separation between two distantly related groups, the partial separation disappears when a full representation of different cell types from an organism are included in the analysis.

We next characterized the density of neighborhood sizes for each of set of cells described above. Across all four samples, we observe fractal-like density distributions, rather than the gaussian-like distributions that are detectable in our high-dimensional models of attractor structure (Fig S3.4B). This observation combined with the lack of distinct separability between the GABAergic neurons and fibroblasts, suggests that the gene expression patterns of these groups of cells are not generated by an attractor structure.

To address whether this continuum is present due to incremental overlap across groups of cells, or the presence of “bridge cells” joining other separable groups of cells, we performed three additional experiments. First, we titrated back different numbers of randomly sampled cell types from the same well and embryo, to the fibroblast and GABAergic subset. When we apply our $\epsilon$ network characterization to the counts-per-million transformed graph, we observe an incremental decrease in the proportion of GABAergic neurons that are separable from the fibroblast and other cell types (Fig S3.4C). While the proportion of GABAergic neurons that are separable decreases, some separation remains detectable even when 10,000 or about 10x the amount of non-GABAergic neurons are present in the data. The separation only disappears once the additional 30,000 cells from the embryo are added back to the experiment (Fig S3.4A).

We next tested the possibility that a combination of biological variation and noise induces the occurrence of “bridge cells” that connect groups of cells that are actually separate in reality. To do this, we calculated the betweenness centrality across all of our cell-type titration experiments at the $\epsilon$ value at which 50% of the cells are connected. If a cell is acting as a bridge between cell-type groups, we would expect to observe cells with few neighbors, but a high betweenness. Interestingly, we observe a small number of cells with an intermediate number of neighbors and betweenness in the graph generated from the raw UMI counts of fibroblasts and GABAergic cells in isolation where no separation is detected using the $\epsilon$ network approach (Fig S3.4A,C). However, when we apply the counts-per-million transformation, and observe partial separation of these cell types no bridge nodes are observed (Fig S3.4C). Similarly, when we add back the full representation of cell types, or just 10,000 randomly selected cells, we do not observe any cells with intermediate betweenness values and simultaneously moderate or low numbers of neighbors (Fig S3.4C). These findings suggest that in complex tissues, bridge nodes are not obscuring the structure of the data by spuriously connecting groups of cells in dense regions of gene expression space.

Finally, to characterize whether cell-type clusters exist in slightly overlapping versus completely enmeshed regions of gene expression space, we quantify the cluster composition of every cluster of all cells from the “P0 one well mouse” sample as a function of $\epsilon$ values. In the counts-per-million transformed P0 mouse data, we see very few components that are both highly homogenous and contain most of the cells of a given type (Fig. 5B, S5.9F). For instance, we do not see more than 7 clusters that contain more than 50% of the cells of any given annotated cell type, regardless of the $\epsilon$ value we consider. Taken together, these observations favor a model of gene expression that is better represented by a diffuse continuum, rather than slightly overlapping cell-type attractors.

### Section 2: Cell-type gating of CITE-seq and Abseq PBMC data

In the positive controls of multiplexed cell line data and FACS-sorted lymphocytes, cell-type identities were defined prior to scRNA-sequencing, using orthogonal measures of cellular identity. These empirical measurements of ground truth provide an ideal control as information on the transcriptomic profile and cell type identity is available for each cell in the data. These paired measurements allow us to evaluate which conditions or sets of transformations allow transcriptomic profiles from distinct cell types to be separable in epigenetic space. Yet, there exists a scarcity of publicly available data where cell type information and transcriptomic measurements are independently established.

To address this gap in control data, we sought to define cell-type identities in multimodal measurements of lymphocytes, using an approach analogous to FACs-sorting of functionally relevant cell surface proteins. We sought to define cell-type identities in two multi-omics datasets that measured both protein and mRNA expression using either Abseq or Citeseq of Peripheral Blood Mononuclear Cells. To do this, we applied expression level thresholds to the protein counts of "marker-genes" in the Abseq and Citeseq data. Based on the expression levels of marker proteins in our *in silico* 'FACs gates', we then annotated the cell-type identity of each cell in the dataset. While Abseq and Citeseq measure both mRNA and protein levels simultaneously, we used only the protein measurements to annotate these data. We reasoned that, likely due to the larger copy numbers, the larger measurement values of contained a larger amount of information on cellular identity, compared to the mRNA measurements.

We annotated cells from both the mRNA and protein measurements generated using Abseq on PBMC cells using the following gates:

**Tcell:** CD3+, CD56-, CD16-, CD21-, CD11c-, CD34-
**NKcell:** CD3-, CD56+, CD21-, cd11b-, CD11c-, CD34-
**Bcell:** CD3-, CD56-, CD16-, CD21+, CD11c-, CD34-
**Myeloid:** CD3-, CD56-, CD16-, CD21-, CD11c+, CD34-
**HSPC:** CD3-, CD56-, CD16-, CD21-, CD11c-, CD34+

We annotated cells from both the mRNA and protein measurements generated using Citeseq on cells collected from the Spleen and Lymph using the following gates:

**T cell:** CD3+, CD16-, CD21-, CD11c-, CD34-
**NK cell:** CD3-, CD16+, CD94+, CD21-, CD34-
**B cell:** CD3-, CD16-, CD21+, CD11c-, CD34-
**Myeloid:** CD3-, CD16-, CD21-, CD11c+, CD34-
**HSPC**: CD3-, CD16-, CD21-, CD11c-, CD34+

We then applied the standard set of transformations to the Citeseq and Abseq protein and mRNA data, as well as Z-score normalization, the L1 norm, the Hamming norm; and performed our $\epsilon$ network characterization of cell-type separability in epigenetic space.

Using the Abseq measurements of 197 proteins in PBMC data, we are again able to observe separation of the expected cell types when suitable transformations were applied (in this case, CPM, CPM-log+1, PCA, and CPM-log+1-PCA all generated separation) (Fig 1A, and S5.7). These findings demonstrate that our method is sensitive enough to detect cell-type separation in biological data, despite numerous sources of measurement noise. Further, this observation suggests it is possible to develop effective procedures for clustering cells into operationally useful cell types. While these common transformations produce an appropriate mapping for these 197 protein measurements of PBMC data, the same set of transformations were not able to recover the same groups in the same sample, when using the 468 mRNA's that were measured instead of the proteins (Fig 1A). These results are concordant with the observation that the current standard scRNA-seq analysis pipeline fails to produce robust cell-type separation. Additionally, we did not observe any separation of cell type groups in five additional PBMC datasets generated using either BDrhapsody, CITE-seq, or scATAC-seq, including the annotated CITE-seq data discussed above. (Figs. 1, S5.6A,D,E, and 5.7C,D).

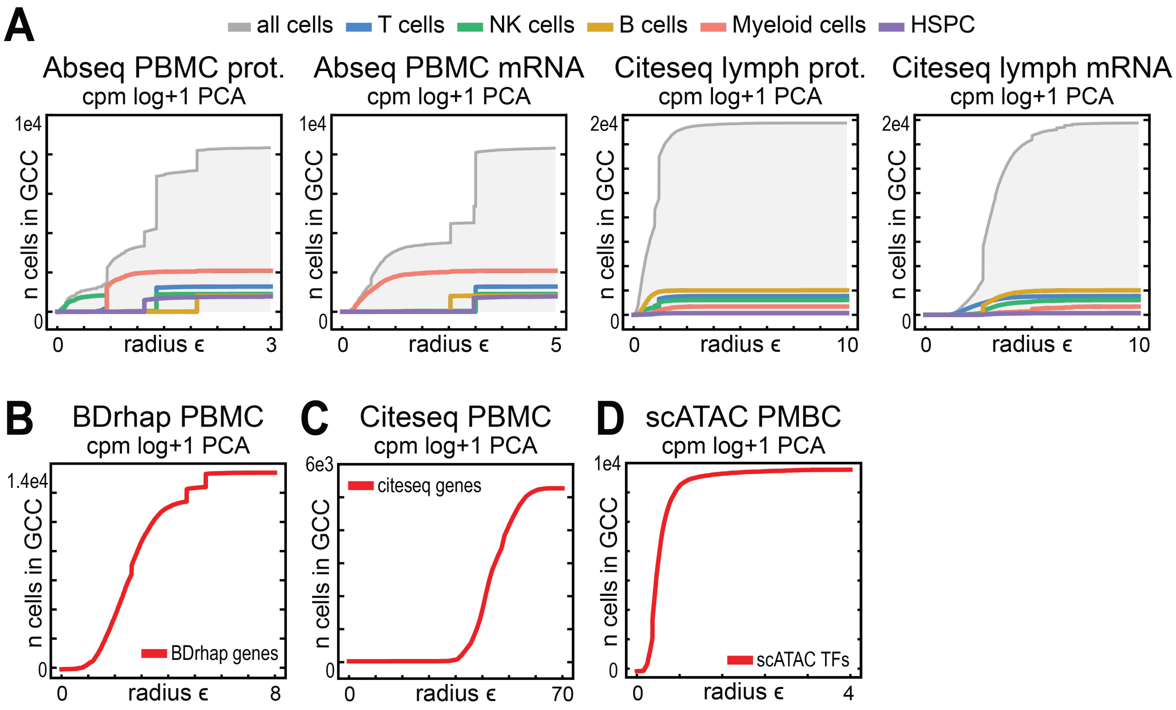

***Figure 1:*** $\epsilon$ network analysis of epigenetic data generated from PBMC cells, after applying the standard set of transformations used in popular scRNA-seq analysis pipelines. (A-D) $\epsilon$ network analysis was performed on transformed count matrices of PBMC cells generated using various sequencing technologies. Plots displaying the number of cells in the giant connected component (GCC) versus the radius of $\epsilon$ used to construct the graph. **A)** Data generated using either protein or mRNA counts from Abseq PBMC cells or Citeseq Lymph cells. The UMI counts were transformed using CPM, then log+1, then PCA. Cell types were annotated using a 'marker gene' expression strategy analogous to FACS. (B,C,D) PBMC count data generated using **B)** the BDrhapsody platform, **C)** the Citeseq platform, or **D)** scATAC-seq. Count data was transformed using CPM, then log+1, then PCA.

**Section 3: The curse of dimensionality**

One form of the "curse of dimensionality" is the fact that, for some types of distributions, distances tend to get more similar as the dimensionality increases. Specifically, over 20 years ago Beyer and co-workers proved that, for some distributions of points in a metric space, the minimum and maximum distances between points sampled from those distributions become more similar as the dimensionality increases, and eventually become indistinguishable as the dimensionality tends to infinity[1]. In other words, if we call the maximum distance in a dataset $D_{max}$ and the minimum distance $D_{min}$, then $\left(\frac{D_{max}}{D_{min}}\right)_n \to_p 1$ where $n$ is the dimensionality of the space and $\to_p$ means "converges in probability," which is defined in more detail below. They argued that this would make notions of "nearest neighbors" in datasets meaningless, although that is not necessarily what their result should be taken to mean (see below). Nonetheless, for certain distributions, it is a fact that increasing dimensionality tends to make all the distances more similar to one another. The Beyer paper and others have argued that this effect can be a problem at even relatively small numbers of dimensions, say 15 or higher, which is obviously much smaller than the dimensionality typically encountered in single-cell genomics datasets (~20,000 or more)[1,2].

Naively, one might expect that this result would imply that the $\epsilon$ networks described in this work would struggle to separate groups of cells. To understand why this might be a problem, say we have a group of cells, and this group of cells is drawn from a distribution that has the $\frac{D_{max}}{D_{min}} \to 1$ property described above. Since all the cells become equidistant, it would be impossible to define a value of the $\epsilon$ distance cutoff that could separate them into multiple groups. The limit described above means that all distances converge to the same value; let's call that value $d_T$ for the "typical" distance. In this scenario, we would go from a completely disconnected graph at $\epsilon < d_T$ to a completely connected graph at $\epsilon > d_T$, meaning the giant component graphs shown in Fig. 2B of the main text would always be guaranteed to be a single step function. In other words, if this version of the curse of dimensionality holds for cell types, then we would be guaranteed *not* to see separation between cell type groups when dimensionality is high, as it typically is for scRNA-seq and related single-cell epigenetics data. This would imply that our findings are an artifact of the dimensionality of the data, rather than a real reflection of a lack of separation between cell types in biological reality.

Interestingly, however, we find empirically that our $\epsilon$ networks ***can*** separate groups of cells in extremely high-dimensional datasets. Perhaps most convincingly, as shown in Fig. 3D, CPM transformation of data from two different cell lines (NIH 3T3 cells and A20 cells) results in clear separation between the two cell type groups even in more than 15,000 dimensions. This demonstrates that it is possible for $\epsilon$ networks to separate cells into different groups, even at dimensionalities that would be very likely to suffer from the above-mentioned curse. We also showed that even simple Gaussian mixture models, or Negative Binomial mixture models, give rise to step-like behavior in the giant component despite extremely high dimensionality (~10,000 dimensions, see Figs. S3.2A,B). Of course, these are not infinite-dimensional examples, but there are cases where the dimensionality is much higher than the 15 or so discussed in the literature[1].

How do we reconcile our findings of separable distributions in real data with these previous theoretical results? The key issue here is that the results in the Beyer paper, and in subsequent works, hold for cases where points are sampled from the *same distribution*. If points are sampled from *different distributions* that are sufficiently separated, then the "curse of dimensionality" actually has the opposite meaning. Below, we prove a corollary to Theorem (1) in Beyer et al[1] that shows that, if two distributions suffer from the curse of dimensionality, then $\epsilon$ networks can separate the two distributions with probability 1 in the limit as $n \to \infty$, provided the distributions are far enough apart. Precise definitions of "far enough apart" are given below, but just ensure that any two points from the two different distributions are closer to points in their own distribution than they are to any arbitrary point drawn from the other distribution.

Preliminaries

Here, we (mostly) follow the notation of Beyer and co-workers, to make the corollary below easier to follow based on their results[1]. We will concern ourselves with real-valued vector spaces of dimensionality $n$ and will keep the dimensionality implicit, so $\boldsymbol{x} \in \mathbb{R}^{\boldsymbol{n}}$. The probability of some event $e$ will be denoted $P[e]$. For a random variable $X$, we denote the expected value of that variable $\mathrm{E}[X]$ and its variance $\mathrm{Var}[X] \equiv \mathrm{E}[X^2] - (\mathrm{E}[X])^2$. Vector-valued random variables are also denoted in bold face, i.e. $\boldsymbol{X}$. We will consider a finite sample of $N$ points from the vector space.

Although Beyer et al.[1] does not explicitly define what a "distance" is, and instead proves the theorem in a rather general setting, we will use the following standard definition of the p-norm:

**Definition 1** A p-norm is a function that maps a pair of vectors $\mathbf{x}$ and $\mathbf{y}$ to a non-negative real number. So we have:

$$\mathrm{d_p}: \mathbb{R}^\mathrm{n} \times \mathbb{R}^\mathrm{n} \to \mathbb{R}_+.$$

The function is defined:

$$\mathrm{d_p}(\mathbf{x}, \mathbf{y}) = \left( \sum_{\mathrm{i}=1}^{\mathrm{n}} |\mathrm{x_i} - \mathrm{y_i}|^\mathrm{p} \right)^{\frac{1}{\mathrm{p}}}$$

where the $\mathrm{x_i}$'s and $\mathrm{y_i}$'s are the coordinates of the vectors $\mathbf{x}$ and $\mathbf{y}$. Note we will take $\mathrm{p}$ to be a natural number (i.e. $\mathrm{p} \in \{1,2,3,\ldots\}$), and so the function $\mathrm{d_p}$ defined above is obviously a metric. As a result, $\mathbb{R}^\mathrm{n}$ endowed with $\mathrm{d_p}$ is a metric space. Note that the definition depends on the dimensionality of the vectors but that dependence is kept implicit for notational convenience.

It is also helpful to define the following form of convergence:

**Definition 2** A sequence of random, vector-valued variables $\mathbf{X_1}, \mathbf{X_2}, \ldots$ of constant dimensionality $\mathrm{n}$ **converges in probability** to a constant vector $\mathbf{x}$ if, for all $\delta \in \mathbb{R},\ \delta > 0$, the probability of the (2-norm) distance between $\mathbf{X_m}$ and $\mathbf{x}$ being less than $\delta$ is 1 in the limit as $\mathrm{m}$ goes to infinity. In symbols, we have:

$$\forall \delta > 0,\ \lim_{\mathrm{m} \to \infty} \mathrm{P}[\mathrm{d_2}(\mathbf{X_m}, \mathbf{x}) \le \delta] = 1.$$

If this is the case, we write $\mathbf{X_m} \to_\mathrm{p} \mathbf{x}$.

These two definitions out of the way, we can develop the final preliminaries. For our purpose, we will forgo the distinction between "data" and "query" distributions made in Beyer et al[1]. This is immaterial for our purposes, as nothing in the proof requires these distributions to be different, and this more accurately represents the case in which we are interested, namely, the distances between points in a given dataset. We will have a sequence of "data distributions," $F_1, F_2, \dots, F_n, \dots$, which are just probability distributions on vectors in $\mathbb{R}, \mathbb{R}^2, \dots, \mathbb{R}^n, \dots$; we will leave the notion of a "probability distribution" implicit. When a random variable takes on values drawn from such a distribution, we write $\boldsymbol{X} \sim F_i$.

Finally, say we have a finite sample of points $X_n = \{\boldsymbol{x}_1, \boldsymbol{x}_2, \dots, \boldsymbol{x}_N\}$, each representing a realization of the random variable $\boldsymbol{X} \sim F_n$; note each vector thus has dimensionality $n$. We choose an arbitrary point from this set $\boldsymbol{x}_i$ and define $D_{\max,p}(\boldsymbol{x}_i, X_n) \equiv \max\{d_p(\boldsymbol{x}_i, \boldsymbol{x}_j) \mid \boldsymbol{x}_j \in X_n\}$ which is just the max distance between all the points in $X_n$ and the chosen point $\boldsymbol{x}_i$. We can similarly define $D_{\min,p}(\boldsymbol{x}_i, X_n)$. This leads to the following theorem from Beyer et al[1]:

**Theorem 1** Given a family of distributions $F_1, F_2, \dots, F_n, \dots$, and, for each distribution, fix a sample $X_n$ and a point from that sample $\boldsymbol{x}_i$. If the following criterion is met:

$$\lim_{n \to \infty} \mathrm{Var}\left[\frac{d_p(\boldsymbol{x}_i, \boldsymbol{x}_j)}{\mathrm{E}[d_p(\boldsymbol{x}_i, \boldsymbol{x}_j)]}\right] = 0$$

(where $\mathrm{E}[d_p(\boldsymbol{x}_i, \boldsymbol{x}_j)]$ is the expected value of the distance between our chosen point and all other points in the sample), then we have:

$$\frac{D_{\max,p}(\boldsymbol{x}_i, X_n)}{D_{\min,p}(\boldsymbol{x}_i, X_n)} \to_{\mathrm{p}} 1.$$

We provide this result here without proof; the proof is fairly straightforward and is given in Beyer et al[1]. We actually don't need to require that the metric in question be a p-norm; the theorem will hold for any metric with the requisite variance property. Here we focus on the p-norm simply since it is the most common example of a family of metrics used to analyze single-cell data.

This theorem indicates that, if the variance in the distances between an arbitrary point and the other points in the sample, relative to the average of that distance, limits to 0 as the number of dimensions tends to infinity, then the ratio between the maximum and minimum distance in the distribution will tend to 1. In other words, the max and min distances become the same. Since all the points are drawn from the same distribution, and we just chose one such point arbitrarily, it follows that the theorem holds for *all* the points in the distribution. This means that, in the limit of infinite dimensionality ($n \to \infty$), *all the points become equidistant from each other.*

As a result, we will say that any distribution with the property:

$$\lim_{n \to \infty} \mathrm{Var}\left[\frac{d_p(\boldsymbol{x}_i, \boldsymbol{x}_j)}{\mathrm{E}[d_p(\boldsymbol{x}_i, \boldsymbol{x}_j)]}\right] = 0$$

is an ***asymptotically equidistant*** distribution under the given p-norm.

As mentioned above, this generates an apparent problem for our analysis, since it would seem that these distributions would be impossible to separate using an $\epsilon$ network. Note, however, that the above theorem works with a sample from a *single distribution*. We are generally concerned, however, with cases where points are sampled from *different distributions*; notions of attractors aside, the simplest interpretation of the Waddington's landscape picture is that different cell types will be sampled from different regions of epigenetic space.

So, say that $\mathrm{X}_1, \mathrm{X}_2, \ldots, \mathrm{X}_n, \ldots$ and $\mathrm{Y}_1, \mathrm{Y}_2, \ldots, \mathrm{Y}_n, \ldots$ are two families of distributions of increasing dimensionality that are both individually asymptotically equidistant under a given p-norm. Now take a sample from each of these distributions at a given dimensionality $n$: $A_n = \{\boldsymbol{a}_1, \boldsymbol{a}_{2,} \ldots, \boldsymbol{a}_N\}$ and $B_n = \{\boldsymbol{b}_1, \boldsymbol{b}_{2,} \ldots, \boldsymbol{b}_M\}$ with $A_n$ being a sample from $\boldsymbol{X} \sim \mathrm{X}_n$ and $B_n$ being a sample from $\boldsymbol{Y} \sim \mathrm{Y}_n$.

Because both distributions are asymptotically equidistant under the given p-norm, for that norm we can define $D_A \equiv D_{\min,p}(\boldsymbol{a}_i, A_n)$ and $D_B \equiv D_{\min,p}(\boldsymbol{b}_j, B_n)$. Note that these are the minimum distances for a pair of arbitrary points $\boldsymbol{a}_i$ and $\boldsymbol{b}_j$. Since all the points become equidistant in the limit, the specific point chosen for this definition will not matter, but it is convenient to choose a specific pair of points as described below.

**Corollary 1** Say $\mathrm{X}_1, \mathrm{X}_2, \ldots, \mathrm{X}_n, \ldots$ and $\mathrm{Y}_1, \mathrm{Y}_2, \ldots, \mathrm{Y}_n, \ldots$ are two families of distributions of increasing dimensionality that are both individually asymptotically equidistant under a given p-norm. Take samples $A_n$ and $B_n$ as described above. Then, if we can choose two points from the two samples $\boldsymbol{a}_i$ and $\boldsymbol{b}_j$ such that:

$$d_p(\boldsymbol{a}_i, \boldsymbol{b}_j) > D_A + D_B + \max\{D_A, D_B\},$$

then there exists some $\epsilon \in \mathbb{R}_+$ such that the resulting $\epsilon$ network separates the two groups into two components, each consisting purely of points from $A_n$ and $B_n$ respectively, with probability 1 as $n \to \infty$.

*Proof* First we establish that all points in $A_n$ will be connected with probability 1 in the limit. Put $\epsilon = \max\{D_A, D_B\}$. Then note that, by Theorem 1, we have:

$$\frac{D_{\max,p}(\boldsymbol{a}_i, A_n)}{D_{\min,p}(\boldsymbol{a}_i, A_n)} \to_{\mathrm{p}} 1$$

as $n \to \infty$ because $A_n$ belongs to an asymptotically equidistant family of distributions. This directly implies $P[d_p(\boldsymbol{a}_i, \boldsymbol{a}_j) \leq \epsilon] \to 1 \; \forall \, \boldsymbol{a}_j \in A_n$ as $n \to \infty$. Note that, if $d_p(\boldsymbol{a}_i, \boldsymbol{a}_j) \leq \epsilon \; \forall \, \boldsymbol{a}_j \in A_n$, then all the points in that set will be connected to the single point $\boldsymbol{a}_i$ at that value of $\epsilon$, and, as such, they will trivially all be in the same connected component of the graph.

The argument for $B_n$ is identical to the one above. So all points from $A_n$ will be in a connected component with all other points from $A_n$, and all points from $B_n$ will be in a connected component with all other points from $B_n$.

All that remains is to show that no points from $A_n$ will be connected to any point from $B_n$. Recall that we have at least one $\boldsymbol{a}_i$ and $\boldsymbol{b}_j$ such that $d_p(\boldsymbol{a}_i, \boldsymbol{b}_j) > D_A + D_B + \max\{D_A, D_B\}$, and that we

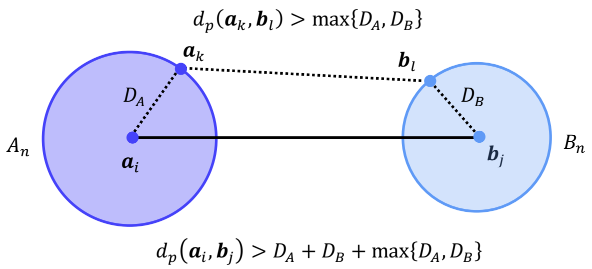

**Fig. 2** Schematic of the scenario in corollary 1. We have two distributions of points, $A_n$ and $B_n$, and two arbitrary points $\boldsymbol{a}_i$ and $\boldsymbol{b}_j$, with the condition $d_p(\boldsymbol{a}_i, \boldsymbol{b}_j) > D_A + D_B + \max\{D_A, D_B\}$. Note that $D_A$ and $D_B$ are the "limiting distances" of the two distributions. The triangle inequality guarantees that the distance between any two additional arbitrary points $\boldsymbol{a}_k$ and $\boldsymbol{b}_l$ will be larger than $\max\{D_A, D_B\}$. So we can thus set $\epsilon = \max\{D_A, D_B\}$ to separate all points from $A_n$ and $B_n$ into two different connected components.

have put $\epsilon = \max\{D_A, D_B\}$. Clearly $\boldsymbol{a}_i$ and $\boldsymbol{b}_j$ will not be connected with each other at this value of $\epsilon$. Now posit that there exists some pair of points, $\boldsymbol{a}_k \in A_n$ and $\boldsymbol{b}_l \in B_n$, such that $d_p(\boldsymbol{a}_k, \boldsymbol{b}_l) \leq \epsilon$ (which would be required for these two points to be connected in the graph). Because $d_p$ is a metric, it obeys the triangle inequality, and so:

$$d_p(\boldsymbol{a}_i, \boldsymbol{b}_j) \leq d_p(\boldsymbol{a}_i, \boldsymbol{a}_k) + d_p(\boldsymbol{a}_k, \boldsymbol{b}_l) + d_p(\boldsymbol{b}_l, \boldsymbol{b}_j),$$

(see Fig. 2 for a schematic). Because of the asymptotic equidistance property of $A_n$ and $B_n$, we have that $d_p(\boldsymbol{a}_i, \boldsymbol{a}_k) \to_{\mathrm{p}} D_A$ and $d_p(\boldsymbol{b}_l, \boldsymbol{b}_j) \to_{\mathrm{p}} D_B$ as $n \to \infty$ by Theorem 1[1]. Given this convergence, we can re-write the above inequality as:

$$d_p(\boldsymbol{a}_i, \boldsymbol{b}_j) \leq D_A + D_B + d_p(\boldsymbol{a}_k, \boldsymbol{b}_l).$$

Say we then have $d_p(\boldsymbol{a}_k, \boldsymbol{b}_l) \leq \epsilon = \max\{D_A, D_B\}$. This gives:

$$d_p(\boldsymbol{a}_i, \boldsymbol{b}_j) \leq D_A + D_B + \max\{D_A, D_B\},$$

which obviously directly contradicts the condition of the corollary, namely:

$$d_p(\boldsymbol{a}_i, \boldsymbol{b}_j) > D_A + D_B + \max\{D_A, D_B\}.$$

So this means that, if we set $\epsilon = \max\{D_A, D_B\}$, the triangle inequality guarantees that $d_p(\boldsymbol{a}_k, \boldsymbol{b}_l) > \epsilon \; \forall \; \boldsymbol{a}_k \in A_n, \boldsymbol{b}_j \in B_n$ with probability 1 as $n \to \infty$. In other words, we cannot have any point from $A_n$ connected to any point from $B_n$.

Combining the two arguments above, at $\epsilon = \max\{D_A, D_B\}$, we have all of the points from $A_n$ in a single connected component, and similarly all the points from $B_n$ in a single connected component. But no points from $A_n$ and $B_n$ can be connected to each other. Thus there must be two separate connected components that consist only of points form the corresponding distributions. So, with probability 1 as $n \to \infty$, the $\epsilon$ network approach separates the two sets into two distinct connected components. ■

If we have more than two distributions, then obviously the above corollary will guarantee that they will all be separable, so long as the pairwise distance between each of them is large enough. So this means an arbitrary collection of asymptotically equidistant distributions will be separable in the $\epsilon$ network sense so long as the condition $d_p(\boldsymbol{a}_i, \boldsymbol{b}_j) > D_A + D_B + \max\{D_A, D_B\}$ holds for some arbitrary pair of points drawn from every pair of distributions.

This result is relatively straightforward, but may at first seem to contradict the result from Theorem 1. In particular, imagine we combine $X_1, X_2, \ldots, X_n, \ldots$ and $Y_1, Y_2, \ldots, Y_n, \ldots$ into a single family $F_1, F_2, \ldots, F_n, \ldots$; why would Theorem 1 not hold for that family? Note that the family $F_1, F_2, \ldots, F_n, \ldots$ would be a family of *mixture* distributions. This would involve constructing $F_n$ by having some probability of sampling from $X_n$ and some probability of sampling from $Y_n$. Speaking loosely, we could write $F_n = P[x]X_n + P[y]Y_n$. How does Theorem 1 not apply to this case?

Note that the key condition of Theorem 1 is that the variance in the relative distance distribution (i.e. the distances between all the points, relative to the mean) has to go to zero. The condition of the corollary, however, requires that points from these two families of distributions, namely $X_n$ and $Y_n$, have to be *sufficiently separated* from one another at every value of $n$. Thus, while the relative variance *within* each distribution will go to 0, the variance *between* each distribution (and thus the variance of the mixture distribution $F_n$) cannot go to 0. In other words, if the data has sufficient structure, then the requirements for Theorem 1 will not hold for that data in the limit. Indeed, the "curse of dimensionality" in this context *guarantees* separability, rather than suggesting the two distributions will become confused as the dimensionality goes to infinity.

### Section 4: Interpretations of Waddington's landscape and the sensitivity of $\epsilon$ networks

One of the critical questions about the $\epsilon$ network approach we apply here is: how far apart do groups of cells need to be in order for this approach to reliably detect them? In other words, what has to be true for us to conclude that the data is consistent with Waddington-like cell type groups in any given epigenetic space?

To start with, it is perhaps instructive to revisit Waddington's landscape and the predictions that it makes. Here, we will focus purely on the picture itself and how it relates to the distributions of cells in epigenetic space:

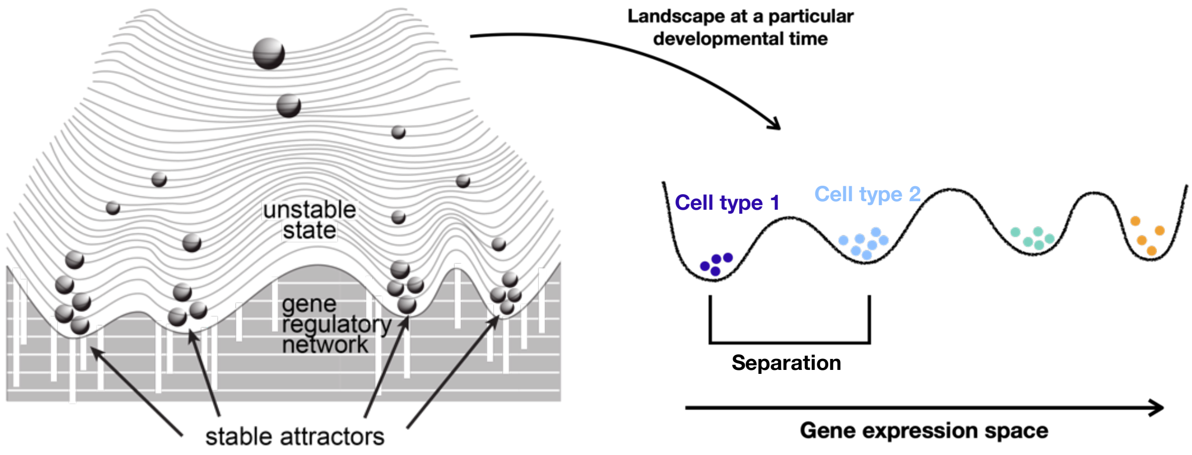

**Figure 3.** Predictions of Waddington's Epigenetic Landscape. To the right, we have the standard picture of the epigenetic landscape, reproduced from Fig. 1 in the main text. If we choose one particular developmental time, say the final "fully differentiated" state, this corresponds to a single horizontal "slice" through the landscape, shown to the left. At that time, we have several different valleys on the landscape. The x-axis of this landscape is a 1-dimensional schematic of a (potentially) high-dimensional epigenetic space, which is usually taken to be gene expression space (but could be a transformed version of that space or even a different epigenetic space entirely). Just from a purely visual perspective, there is separation between the bottoms of the "valleys" in the landscape in that epigenetic space. In the modern mathematical treatment of the landscape, the bottoms of the valleys correspond to point attractors of the underlying gene regulatory network[10-14], with separatrices between the corresponding basins of attraction (i.e. the "peaks" or "hills" on the landscape). Given that cells are not exactly identical in their epigenetic space, due to gene expression noise or other factors, the landscape is typically drawn with groups of cells at the bottom or each valley. While we have drawn this picture ourselves, similar representations may be found in refs[10-14].

If we take a given time point during development (say, for simplicity, the "last" time point where the cells have adopted terminally differentiated cell fates), the prediction made by the landscape is clear: cells should be found in the "bottoms" of the valleys, which are visually separated from one another in the picture (Fig. 3, right). Unfortunately, the landscape itself is just a picture, and as such makes no precise mathematical statement regarding the distributions of cells in epigenetic space. But the version shown to the right in Fig. 3 schematizes every version of the landscape in which the cell types are represented by populations (i.e. more than just a single cell at the bottom of any given valley) of which we are aware.

By far and away the most common mathematical interpretation of this landscape describes the bottoms of the valley as point attractors of the dynamical system that regulates the epigenetic state of the cell. The nature of this theory, and why it entails a focus on macromolecular concentrations as the corresponding epigenetic state space, is described in detail in the next section. That perspective on the Waddington picture relies on deterministic dynamical systems, which do not admit variation in epigenetic state in these attractors. In other words, the attractor states are single points, and cell within the basin of attraction will asymptotically approach these points with time. While some work has been done to explore these dynamics in stochastic models of gene regulation[3]. the variation around these attractor states is often not explicitly considered in modeling studies. As such, we imagine that some degree of heterogeneity in gene expression state will characterize cells in the neighborhood of the attractor, which is represented visually as populations of cells near, but not exactly at, the bottom of the attractor well in the landscape picture (Fig. 3).

The question then becomes, how do we interpret this landscape in terms of the predictions it makes for single-cell data? Since there is heterogeneity in this data, we cannot expect all cells of a given time to be exactly "on top" of the attractor state in the landscape. As such, we need to develop an expectation for how the data should be distributed in epigenetic space if Waddington's landscape holds and further demonstrate that the $\epsilon$ network approach will be able to detect that structure if it exists.

To formalize this mathematically, we start by positing that every cell has a definitive "cell type" class to which it naturally belongs. We will call the set of possible cell types $T$, and since this will evidently be a finite set, we can say $T = \{1, 2, \ldots, n\}$, where $n$ is the total number of cell types in our data, without loss of generality. So we can call these "cell type 1," "cell type 2," etc. In addition to this, we have a set of cells, and each cell is associated with some epigenetic state. We will call our set of cells $C$ and our epigenetic space $E$; note that $C$ can be thought of as a finite subset of $E$. We take $E$ to be a finite-dimensional vector space endowed with some metric $d$, and require that the distance so defined is finite between every pair of cells in $C$ (in other words, no cell is infinitely far away from any other cell, which is certainly the case for any practical dataset we have considered in this work). Finally, each cell has a definitive type, meaning we have a function $f: C \to T$ that assigns each cell to its corresponding cell type. For convenience we will define $C_1 \subset C$ as the subset of cells of "type 1", meaning $C_1 = \{c | c \in C, f(c) = 1\}$. We will assume that there are at least two non-empty sets of cell types comprising $C$ (hence $C_1$ being defined as a strict subset above).

Note that the above may be a bit abstract, but corresponds to the cases typically considered in single-cell epigenetic studies. In such studies, we always have a finite number of cells, and each cell is represented by a point in some epigenetic space. That space could be a space of UMI counts, or a transformed version of that like log CPM+1-HVG-PCA space; regardless, we always represent each cell as point in a vector space. On top of this, we are assuming that every cell has a "true" cell type; in practice we might not *know* the cell type for each cell, but every cell has a definitive cell type, so the function $f$ exists. Ultimately the argument below does not depend on whether we can determine the cell type for each cell empirically; it just has to exist in theory.

To progress further, we will assign to each cell type one of the valleys on the landscape. We will posit that each valley has a characteristic location in the epigenetic space, which we will call $a_i \in E$ with $i \in T$. This is just the "bottom" of the valley on the landscape corresponding to that cell type (Fig. 3 right); in the dynamical systems perspective this is just the point attractor for that basin of attraction. So every cell type has one such characteristic, or archetypal, position on the epigenetic landscape. Again, we might not be able to calculate the value of this position exactly from data, but that is not important for the argument below.

Now, for every cell $j$ that belongs to some cell type $i$, we can calculate the distance between that cell and this characteristic location, $d(c_j, a_i)$, where $c_j \in C_i$. We define $C_i^*$ as the set of cells at the minimum distance from $a_i$. As long as every distance to this location is unique, this set will just consist of the cell that is closest to the bottom of the basin for that cell type on the landscape, but in general there could be "ties" so we consider this to be a set with potentially more than one member.

Using this framework, we now consider building an $\epsilon$ network. As a reminder, in this context an $\epsilon$ network is just an undirected graph where the set of nodes is $C$ and two nodes/cells $c_j$, $c_k \in C$ have an edge if $d(c_j, c_k) \leq \epsilon$. At any given value of $\epsilon$, this graph will consist of a set of connected components, where a connected component is just a subset of nodes where every member of that subset can be connected to every other member of that subset by some path in the graph.

We say that a cell $c_j$ is topologically connected to the basin for a cell type $i$ at $\epsilon$ if $c_j$ and a member of $C_i^*$ are part of the same connected component. In other words, this just means that, if we construct a series of neighborhoods of radius $\epsilon$ around the points in our dataset, then there is a path in our dataset through such neighborhoods from that cell to the bottom of the basin for cell type $i$, or, more precisely, at least one of the closest representatives of that point on the landscape that we have available in our dataset.

For every cell type $i$, we can define a "special" value of $\epsilon$ as the minimum $\epsilon$ such that there exists a connected component in the graph that has the following properties:

1. More than 50% of the cells of that type are in that component
2. At least one member of $C_i^*$ is in that component

To be slightly more formal, call the set of values of $\epsilon$ where there is at least one connected component that satisfies properties 1 and 2 above $\{\epsilon_{50}\}_i$. We then define $\epsilon_i^* = \inf \{\epsilon_{50}\}_i$. This is just the minimum value of $\epsilon$ that we need to get more than 50% of the cells of that type into the same connected component with a cell that is closest to the bottom of the basin for that cell type. Note that we have chosen this value of 50% arbitrarily; one could choose any fraction of cells of the given type that one likes (the first criterion would then read "More than $q$ percent of the cells of that type are in the component"). We should also note that some cells of the other types could be part of that component—there is no restriction here regarding that.

We define the set of nodes in the component that contains more than 50% of cells of type $i$ and a member of $C_i^*$ at this minimal value of $\epsilon$ as $B_{50,i}^*$. We can consider this set an estimate of the cells that belong to the "basin" of cell type $i$. Note that by construction $B_{50}^*$ is *unique*; there cannot be two separate components of the graph that both contain more than 50% of cells of type $i$. So, in a sense this is the smallest possible unique characterization of the basin, subject to the idea that at least 50% of cells of type $i$ need to be topologically connected to the "bottom" of the basin.

Take two cell types $i$ and $k$; without loss of generality say $\epsilon_i^* \geq \epsilon_k^*$. In other words, choose $i$ to be the cell type with the larger value of $\epsilon$ required to connect more than 50% of those cells with at least one cell of that type that is closest to the characteristic location of that cell type on the landscape. There are two possible scenarios for this situation: at that value of $\epsilon$, either a cell from $C_k^*$ is part of $B_{50,i}^*$, or not. In other words, in one scenario, $B_{50,i}^*$ *also contains* a cell closest to the bottom of the valley for a different cell type $k$, and in the other scenario, it does not.

Consider the case where $B_{50,i}^*$ <u>does not</u> contain a cell from $C_k^*$; this is the case where the two "basins" of the two cell types are not topologically connected at $\epsilon_i^*$. We can then prove the following simple statement:

**Proposition 1** If $B_{50,i}^*$ does not contain a cell from $C_k^*$, then the giant component must have a discrete jump in size corresponding to at least 50% of cells of type $i$, or at least 50% of cells of type $k$, as a function of $\epsilon$.

*Proof* There are 2 possible scenarios here. Consider first the case where, at $\epsilon = \epsilon_i^*$, $B_{50,i}^*$ is *not part* of the giant component $G$ of the graph at that value of $\epsilon$, which we will call $G_\epsilon$ for convenience. Since $B_{50,i}^*$ and $G_\epsilon$ are both subsets of the set of all cells $C$, this scenario simply entails $B_{50,i}^* \not\subset G_{\epsilon_i^*}$. Define $\{\epsilon_{\mathrm{ni}}\}$ as the set of all values of $\epsilon$ where no cell from $B_{50,i}^*$ is in the giant component; this is just the set of $\epsilon$'s where $B_{50,i}^* \not\subset G_\epsilon$. Obviously this set is non-empty by construction for this scenario. Put $\epsilon_{\max} = \sup\{\epsilon_{\mathrm{ni}}\}$; since all distances between all cells in our landscape are finite by assumption, $\epsilon_{\max}$ is finite. Slightly less formally, $\epsilon_{\max}$ is just the largest value of $\epsilon$ so that the cells from $B_{50,i}^*$ are not part of the giant component. Now, for any $\epsilon > \epsilon_{\max}$, this means that $B_{50,i}^*$ will be a part of the giant component. So, as $\epsilon$ increases past $\epsilon_{\max}$, the size of the giant component must increase by the number of nodes in the component that contains $B_{50,i}^*$ at $\epsilon_{\max}$. Since $B_{50,i}^*$ by definition contains more than 50% of cells of type $i$, this discrete increase will be at least 50% of the number of cells of that type.

In the second scenario, $B_{50,i}^*$ is in the giant component at $\epsilon_i^*$. Recall that, by construction, no element of $C_k^*$ is in $B_{50,i}^*$, and also that $\epsilon_i^* \geq \epsilon_k^*$. So we have $B_{50,k}^* \not\subset G_{\epsilon_i^*}$; in other words, the giant component *cannot* contain the set of cells in the "basin" for cell type $k$. If it did, then $B_{50,i}^*$ would be in the same component as a cell from $C_k^*$, since by definition there is some cell from that set in $B_{50,k}^*$, which contradicts the condition of the proposition. So cells from $B_{50,k}^*$ cannot be in the giant component at this value of $\epsilon$. We can use an identical argument to the one above to show that there is also an $\epsilon_{\max}$ for this case, and as $\epsilon$ increases past this $\epsilon_{\max}$, the component containing $B_{50,k}^*$ will join the giant component. This will cause a discrete jump in the size of the giant component, corresponding to more than 50% of the cells of type $k$. ■

This simple argument shows that, if there is *no discrete jump* in the size of the giant component, then we can conclude that $B_{50,i}^*$ and $B_{50,k}^*$ must be topologically connected. That means that we cannot find a value of $\epsilon$ such that more than 50% of the cells of cell type $i$ are connected to the characteristic location of that cell type on the landscape, but *not* connected to the characteristic location of another cell type. Note that, while there must be a discrete jump in the size of the giant component if the two basins are separate, the converse is not true; the presence of such a jump does not prove that any two basins (defined as they are above) are separate. So this is a necessary but not sufficient condition. Regardless, if we don't see a jump, then we can conclude that, for the landscape $E$ and metric $d$, there are no separable basins in the sense defined above.

We should note that the idea that these basins should only contain 50% of the cells is quite permissive. Indeed, drawings like the one shown in Fig. 3 always show 100% of the cells of the same type as being in the corresponding basin; indeed, in Waddington's landscape, the fact that a cell is in a basin is what *makes* the cell that cell type. In other words, in the theory of Waddington's landscape, the position of a cell in the basin of attraction of a cell type requires that that cell be of that type (see more on this section 5 below). Interestingly, the above argument can be made for any relevant cutoff $B^*_{q,i}$ where $q$ is whatever percentage of cells one is willing to require that must be topologically connected to an element of $C^*_i$ (i.e. the cell closest to the "bottom" of the basin). Regardless, even with a highly permissive cutoff of 50%, we should still see a jump in the size of the giant component.

How large these jumps should be depends on the relative sizes of the cell type populations in any given dataset. Most scRNA-seq studies contain several "cell type clusters" that consist of at least a few thousand cells, and in general the size and composition of these clusters conform to historical expectations regarding the composition of the tissues/organisms under study[4–7]. In other words, it is clear that the field generally expects to find cell type groups consisting of at least a few thousand cells. In our analysis, we rarely observe discrete jumps of such a size, even in transformed spaces (Figs 5A, and Figs S5.1-S5.8), and, when we do, that represents a clear indication of separability in the corresponding epigenetic space (as for the cell line data shown in Fig. 3D in the main text). In most datasets, however, is extremely rare for jumps to exceed 5% of the dataset even when they are observed. Thus, even in cases where we do observe discrete jumps in the size of the giant component, it is rare for these to be large enough to be reasonably thought to represent the cell type populations that are typically expected. And in many datasets we cannot observe discrete jumps at all, even in highly transformed spaces (Figs 4 and 5, and Figs S4.1,S4.2,S5.1-S5.8, and S5.11-19). This suggests that it is not possible to find a consistent set of transformations that can separate cells into Waddington-like groups given available epigenetic data.

One could argue that the requirement for topological separation used as the basis for the argument above is inappropriate. In other words, one might argue that Waddington's landscape does not actually suggest that $B^*_{50,i}$ and $B^*_{50,k}$ be separate from one another at $\epsilon^*_i$. While every single drawing of the landscape has that property, one might attempt to argue that the Waddington picture is just a schematic of a somehow more complex epigenetic reality.

To asses this perspective, note that we constructed our analysis in Proposition 1 to be fairly general from a mathematical standpoint, which involved considering a number of mathematical technicalities that are not observed in practical analysis of real data. For instance, for the data sets and metrics we considered in this work, essentially all pairwise distances are in practice *unique*, which means edges are added to the $\epsilon$ network one at a time. This doesn't mean that cells have to be added to any component one at a time; obviously, if we add an edge between one cell in one component and another cell in a different component, the two components will combine all at once, resulting in the emergence of a new component that is larger in size. The idea, however, is that *in practice,* we will not observe a component dramatically growing in size at a particular $\epsilon$ because a large number of cells join "independently." Thus, large discrete jumps in the size of a component at a single $\epsilon$ correspond to the merger of two components.

Also, to simplify this discussion of whether our $B_{50,i}^*$ criterion is a reasonable perspective on the Waddington picture, we will ignore cases in the dataset where there are multiple cells of any given cell type with *exactly the same distance* from the characteristic location on the landscape $a_i$. So, here, $C_i^*$ can be thought of as being a single cell, $c_i^*$, which is the "representative cell" that is closest to the bottom of the basin in a given dataset. Whenever cells are actually drawn on pictures of Waddington's landscape, there are always cells at the bottom of the basins, meaning that the picture asserts cells of a given type should be found "near" the bottom. So we will assume that, if Waddington's landscape holds for a given dataset, there is at least some cell in the dataset that is reasonably close to this location on the landscape. In other words, this representative cell should be thought of a as a good representative, sufficiently close to $a_i$ that we can use that location and the location of $c_i^*$ interchangeably in practice.

Given these simplifying observations, what would it mean for $B_{50,i}^*$ to be connected to $B_{50,k}^*$? Simply put, that means that around 50% of cells of a given type are *further away* from cells close to the bottom of the basin than those cells are to cells from a different basin. This might look something like the scenario shown in Fig. 4.

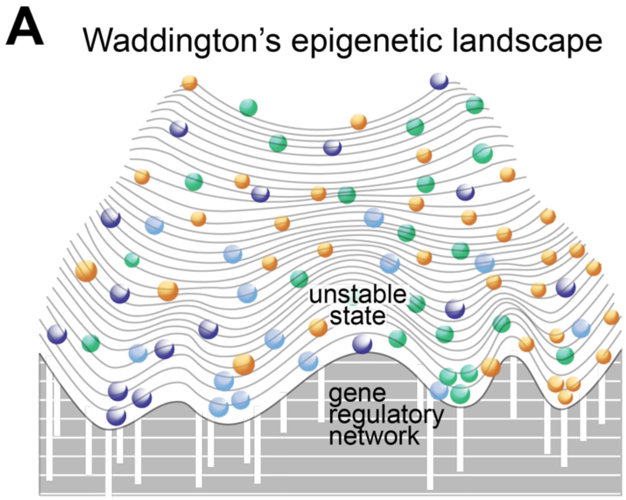

**Figure 4.** Waddington's Epigenetic Landscape as a visual representation of single cell epigenetic data **A.** Schematic of Waddington's landscape, where the epigenetic state of developing cells are spread into different regions of epigenetic space. In this landscape, "basins of attraction" produce a cell type, but whether a cell of a given type expresses the gene expression pattern of its cognate cell type basin is indeterminate. Only 50% of cells of a given type are closer to the bottom of their specific cell type basin than they are to a different cell type basin.

This is clearly not the scenario envisioned by Waddington's landscape. In this scenario, cells of this type are spread into different regions of epigenetic space; if they weren't, then Proposition 1 shows that we would see a discrete jump in the size of the giant component. Indeed, the lack of discrete jumps, particularly in the space of raw UMI counts or perhaps CPM normalized counts (see below) are extremely interesting. In order to avoid having large jumps, cells have to join the giant component in small groups, and the only way to achieve this is to have hierarchically organized data across the space where cells are further and further away from cells in denser regions than they are *from other cells in these less dense regions*. Our density distribution results emphasize this fact (Fig. 6 and S6.2-S6.13).

So, in almost every dataset or transformed dataset we analyze, cells don't mass into groups as suggested by the Waddington picture; instead, cells of different types are all found in regions of hierarchically organized density. This is completely inconsistent with the Waddington picture of "stable groups" canalized into cell fates due to the organization of epigenetic interactions or constraints. In other words, there is no way to imagine a scenario in which a dataset is drawn from a landscape schematized in the Waddington picture that does not have the $B^*_{50,i}$ property that Proposition 1 considers. Note that the above arguments require absolutely no dynamical systems theory as a basis; put simply, Waddington's landscape is a poor visual metaphor for the organization and structure of these data.

**Section 5: Transformations and the biochemical basis of cell fate specification**

The initial analyses we performed focused on the raw UMI counts, which are rarely analyzed directly in the scRNA-seq field. As discussed extensively in the main text, most cell type clustering analyses are performed on highly transformed spaces, including CPM-style normalization, linear and/or non-linear dimensionality reduction, etc. Of course, for most datasets even these transformations don't "work," in the sense that they do not generate Waddington-like distributions on the resulting epigenetic landscape (Figs. 4 and 5, and Figs. S4.1, S4.2, S5.1-5.8, and S5.11-5.19). But why focus so much on the raw UMI counts, rather than just analyzing the data in the typical spaces that are used for the analysis?

The answer to this question lies in the way in which Waddington's landscape is used to provide a *biochemical* explanation for cell fate specification during development/ontogeny. Multicellular organisms, and in particular animals, exhibit cells of incredible phenotypic diversity—compare the ameboid phenotype of a neutrophil to the sessile and highly arborated structure of a neuron. Since all the cells of an adult animal have more-or-less the same genome, the source of these differences cannot be genetic; instead, they must be epigenetic. Also, these cell states are stable: cells don't spontaneously switch from one state to another. In other words, we do not observe neurons suddenly retracting their projections and crawling around.

Waddignton's epigenetic landscape simultaneously explains why these cell states are different and also why they are stable[8–11]. Ultimately, however, we have to ask what the epigenetic state being schematized by the landscape actually is. The current form of the landscape picture universally posits that cells are different because the levels of different macromolecules within the cells are different[11–13]. While these macromolecules could be of many different types, consider the case of protein levels for simplicity. Proteins are understood to be the primary enzymatic and structural components of cells, so it is natural to imagine that, if the levels of different proteins are different, then the resulting phenotypes of the cells would be different. For instance, a neuron will likely express structural proteins that construct and maintain its projections, and of course particular cell-surface receptors and neurotransmitters that allow for neural communication. Instead of expressing these proteins, we would expect a neutrophil to express proteins that allow them to recognize and phagocytose infectious agents.

So, in this perspective, cells of different types are different because they express different macromolecular components at different levels. This allows cell types to have distinct phenotypes and functions, but the question becomes: how are these different levels generated and stably maintained? The modern answer to this question is that some proteins interact with DNA elements like promoters and enhancers, with each other, with the machinery that controls chromatin modification state and accessibility, with DNA elements, etc., to form a complex biochemical network of interactions. This network ultimately controls the flux of RNA, particularly mRNA, production, and translation of these mRNA molecules similarly controls the flux protein production. The modern interpretation of Waddington's landscape holds that these complex networks have a set of stable configurations, corresponding to states with different levels of proteins and other macromolecules in the cell. So, there is one stable state where "neuron expression levels" are maintained, and another stable state where "neutrophils expression levels" are maintained. These different states are the bottoms of the basins on Waddington's landscape, and their stability is visually represented by the hills in between those basins. The idea that a complex regulatory network "underlies" the shape of the landscape has led to many authors drawing a "gene regulatory network" with ropes or ties holding down the basins in the landscape as a mechanism for schematically representing the above notions (see Fig. 1A in the main text, among others)[9,11,14].

The above arguments are compelling, but developing a theory of this kind requires a mathematical framework in which potentially vague ideas like "stability" or "gene regulatory network" can be made precise[15,16]. The most natural such mathematical framework is dynamical systems theory; note that, in the above discussion, we used the term "flux of RNA production." Dynamical systems theory offers a mathematical language for describing how such fluxes depend on the state of the system, and thus is a perfect approach to describing Waddington's landscape more formally. This is typically done using deterministic dynamical systems, although stochastic approaches have also been employed to great success[3]. We focus on the deterministic perspective first.

Applying dynamical systems theory requires a state space of variables whose values will change over time[15–17]. In the case of a biochemical interaction network, the relevant interactions are represented as chemical reactions. For instance, say a protein *T* binds to a promoter for gene *X* and, when bound, changes the rate at which mRNA is transcribed for that gene. We will denote the promoter for gene *X* as *pX* and the mRNA for that gene as *mX*. We would represent this as the following set of simple chemical reactions:

$$T + pX \xrightarrow{k_{on}} TpX$$

$$TpX \xrightarrow{k_{off}} T + pX$$

$$TpX \xrightarrow{k_{tr}} TpX + mX$$

where the symbols on top of each arrow correspond to a "rate constant" for each reaction. Note that focusing on chemical reactions like this is not an arbitrary choice made for convenience; instead, these kinds of reactions are the chemical basis of our understanding of biology.

The next task is to translate a set of reactions like this into a dynamical systems framework. This is done by applying the "Law of Mass Action" (LMA) which is extremely well established and dictates that the flux of a chemical reaction is determined by the "concentrations" of the reactant chemical species[15–21]. The LMA provides a semantics for translating any set of chemical reactions like the ones shown above into a system of Ordinary Differential Equations (ODEs); this is sometimes referred to as "Chemical Reaction Network Theory" (CRNT). For instance, for our reactions above, we have:

$$\frac{d[pX]}{dt} = -k_{on}[T][pX] + k_{off}[TpX]$$

$$\frac{d[TpX]}{dt} = k_{on}[T][pX] - k_{off}[TpX]$$

$$\frac{d[mX]}{dt} = k_{tr}[TpX]$$

where the symbol $[X]$ stands for "concentration of the chemical species X." We will not go into the theory behind the LMA or CRNT, as it is extremely well established, but some references may be found here[15,17]. Mathematically, it is sometimes more natural to talk about systems of ODEs as "vector fields," and we will use the two terms interchangeably[15,16].

It is important to note that the choice of concentrations as the state variables here is not made arbitrarily or for convenience, but rather dictated by the LMA. In other words, it is empirically the case that the flux of chemical reactions is *determined by the concentrations of the reactant species*. So the state space of the system is inherently the space of concentrations of those species. One might note that many models of biochemical reaction networks do not rely purely on the LMA; for instance, it is extremely common to use "Hill Functions" for fluxes of processes like mRNA production from a bound promoter. This might look something like this:

$$\frac{d[mX]}{dt} = k_{tr}\left(\frac{[TpX]^n}{K^n + [TpX]^n}\right)$$

where $k_{tr}$, $K$ and $n$ are constant parameters. While this is not a direct application of LMA, the justification for using these kinds of functions is that they abstract chemical processes, like cooperativity in promoter binding, DNA bending, etc., that one does not wish to model explicitly. Regardless, the relevant state variables here are also molecular concentrations, since the fluxes of the processes that are being abstracted are dictated by those concentrations.

In this framework, we can now be somewhat more precise about the formulation of Waddington's landscape. We can now say that the "gene regulatory network" is a biochemical reaction network of the same type as the one schematized above, though undoubtedly much larger and complex. This directly entails a dynamical system where the variables are the concentrations of chemical species, and the ODEs are generated by the application of the LMA to those reactions. The bottom of each basin then corresponds to a stable equilibrium point of the vector field corresponding to this biochemical reaction network. These equilibrium points may also be referred to as stable "steady states" (to avoid confusion with chemical equilibria, which they are not), "point attractors" (to avoid confusion with limit cycles and strange attractors, which are not single points), or often just "attractors" when the precise application of the term is clear from context.

The biochemical reaction networks underlying cell fate specification thus have multiple possible point attractors; there is a "neuron attractor," a "neutrophil attractor," and so on, each with different levels of macromolecules. The basins on Waddington's landscape correspond to the basin of attraction of the corresponding attractor. Speaking semi-formally, a basin of attraction is a set of points in the state space where, if one "starts" the dynamics from that point, the resulting trajectory will get closer to the governing point attractor over time (formal definitions may be found in a variety dynamical systems texts[15]). The existence of these basins of attraction forms the basis for our understanding of why a neutrophil doesn't suddenly become a neuron, or even a more closely related immune cell type. If a cell in the neutrophil basin experiences some perturbation the concentrations of macromolecules, as long as it remains in that basin, it will naturally "return" towards the macrophage attractor over time.

The above argument is obviously entirely deterministic, and one could question the utility of this framework given that there is obviously a great deal of variation in macromolecular concentrations even between cells of the same type. Interestingly, the LMA is straightforward to formulate in a stochastic context where such variation can be studied in terms of the inherent randomness of chemical reactions at low copy numbers[3,20,22,23]. Ultimately, these models don't generate "point attractors," but rather "steady-state probability distributions" that are multimodal in the underlying space of protein/mRNA levels. Such models can admit spontaneous transitions between cell states due to stochastic fluctuations, so the notion of perfect stability of cell types does not hold for these systems[3]. Nonetheless, the basic idea is the same: the molecular interactions between species generate a landscape in the space of *levels of macromolecules*, and the modes of the probability distribution of cells across that landscape correspond to the different cell types[3]. So, while such stochastic frameworks are more complex, they predict the same basic structure for the data as their deterministic counterparts.

So, given the above, what do we expect in single-cell epigenetic experimental data? The vast majority of available single cell data reports *levels of macromolecules within single cells*. For instance, scRNA-seq generates a "Unique Molecular Identifier (UMI) count matrix," and, in that matrix, if we see an entry of "7" for a gene $j$ in a cell $i$, that means that the experiment detected 7 unique copies of the mRNA corresponding to that gene in that cell. Based on the theory outlined above, we expect that different cell types should have different levels of these macromolecules, at least for a subset of genes that are important for the observed differences in cellular phenotype, either because they have differential functions or because they are involved in the network of cell fate specification. This is particularly true if we consider a dataset that is made up entirely of terminally-differentiated cells, which should be sitting at the "end" of the landscape in their final, stable and fully-specified cell fates (Fig. 3 above). So, it is natural to start by asking whether we can identify attractors in the space of these macromolecular levels. As described in great detail elsewhere in this manuscript, there is no evidence for attractor structure in molecular levels in any of the datasets we have studied.

Careful consideration of the above arguments, however, suggests that considering "raw counts" may be naïve or unfounded. The biochemical reaction network we have described doesn't operate on counts, but rather on *concentrations*, which are counts per unit volume. More specifically, call the number of counts of mRNA for gene $j$ in a cell $i$ "$n_{i,j}$." The attractor in question should really exist in the state space of concentrations of that mRNA. If we call the molar concentration of that mRNA $x_{i,j}$, we are actually interested in:

$$x_{i,j} = \frac{n_{i,j}}{A \cdot V_i}$$

where $A$ is Avogadro's number and $V_i$ is the volume of cell $i$ in liters. Unfortunately, the majority of single-cell technologies are destructive and thus we do not know the volumes of the cells. If the volumes of cells in a population are more-or-less constant, then we can consider $x_{i,j} \propto n_{i,j}$ based on the above equation, and use the count numbers instead.

But cell volume could vary significantly, and since we don't have measurements of cell volume, we cannot say for certain how representative the count data is of actual macromolecular concentrations. We can, however, make a simple approximation, and say that the *total number* *of UMI counts* for a cell is proportional to its volume. In other words, the larger a cell is, the more mRNA counts we will get for that cell in the data. Define the total number of counts in a given cell $i$ as $n_{i,T} \equiv \sum_j n_{i,j}$; this is just the sum of the counts for all genes within that cell. The above assertion then amounts to saying $V_i \propto n_{i,T}$. This gives us:

$$x_{i,j} \propto \frac{n_{i,j}}{n_{i,T}},$$

which suggests that we can use the total number of counts to generate an approximation of concentration.

Interestingly, the above formula is almost identical to traditional CPM normalization, which is:

$$\hat{x}_{i,j} = \left(\frac{n_{i,j}}{n_{i,T}}\right) Q,$$

where $Q$ is a constant that determines the units of the normalized quantity. For instance, if we set $Q = 10^6$, then this is "Counts Per Million." Interestingly, most recent analyses set $Q = 10^4$ despite still referring to this as "CPM" normalization, but ultimately $Q$ is a free parameter set by whomever is analyzing the data.

Since CPM-style normalization could be thought of as proportional to macromolecular concentration, one could reasonably argue that it is the appropriate space for investigating the presence or absence of Waddington-style attractors. Interestingly, for the A20/NIH3T3 cell line data, we see absence of separation in the space of raw UMI counts, but clear, distinct separate groups after CPM normalization. While this might argue for CPM normalized data as the appropriate space in which to conduct all analyses; see below for an analysis of whether this normalization "works" for that data because it is revealing patterns in concentration space that are absent in the space of raw numbers, or if there is some other factor at work.

Aside from those cell line results, CPM normalization essentially never generates distinct groups when the data on macromolecular counts is genome-wide (Figs. 5, S5.1-5.20, 6, S6.2-6.13). We do see some separation in CPM spaces for cases where the molecular count data, particularly protein data, is only available for a few hundred genes, but it is unclear how well the assumption that $V_i \propto n_{i,T}$ holds when that total is calculated over a very small subset of genes (as in the BD Rhapsody protein or Abseq data, Figs. S5.7A, B, S5.8C). Those examples are fairly rare, however; for the vast majority of cases, we don't see separation in either the space of raw levels or the space of CPM normalized data. Interestingly, there is one MERFISH data from the Zhuang group that reported the *volumes* of the cells in addition to the count numbers of the various genes, which is possible because this is a microscopy-based technique[24]. In this case, when we compute the actual concentrations of mRNA in these cells for these data, we see *no separation whatsoever* (Fig. S5.8F).

This brings up the question of the many other transformations that are often applied to the data before analysis. Let's say we start with CPM normalized data and apply log-transformation to it. This is done through the following formula:

$$\hat{t}_{i,j} = \log(\hat{x}_{i,j} + p),$$

where $p$ is a constant "pseudocount" value added to the normalized data because the vast majority of $\hat{x}_{i,j}$ values are $0$, which would clearly cause a problem for this transformation. While $p$ is often set to $1$ (leading to "log CPM+1" transformation), this is another free parameter in the pipeline. As described below, operationally the values of $Q$ and $p$ likely influence the outcome of the analysis by controlling the separation between 0 and non-0 values in the resulting transformed count distributions.

In most datasets we analyzed, regardless of the transformations we applied, we were unable to observe separation, suggesting that there is no consistent space in which such separation can be obtained. For instance, after selecting HVGs, we observe separation in log CPM+1-PCA space for the A20 & NIH3T3 cell lines, Jurkat & Raji cell lines, and the lymphocyte data, but not the mouse bladder, hydra, *C. elegans*, or a host of other datasets (Figs. 5, S5.1-S5.8, S5.11-5.20).

But, let's say through subsequent work we discover some function $g: E \to L$ where $E$ is a space that can be reasonably thought to represent macromolecular levels/concentrations and $L$ is a nonlinearly transformed space where we can consistently observe separation. There are two possible explanations here. On the one hand, it could be that data from $E$ is noisy and, somehow, $g$ "de-noises" the data such that the attractors that are not observable in $E$ because of technical noise can be found in $L$. The only way to assure that such a scenario holds, in the absence of a reliable *a priori* theory of the noise structure of the data in $E$, is to develop more accurate experimental methods that eventually show the attractor structure in $E$.

In the other scenario, we have no attractor structure in $E$, and we actually *generate* it through $g$. Does this provide evidence for Waddington's landscape in the standard biochemical interpretation described in detail above? The answer to that question is no. In order for the biochemical interaction network to be seen as causally generating stable cell types, there *must be* an attractor structure in the space of molecular levels. This is not a matter of choice, but rather an inherent feature of the fact that the theory itself is based on a network of biomolecular interactions generating those stable states. Any attractor in such a system of molecular interactions must exist in the space of molecular concentrations. If such attractors do not exist, then the paradigm describing the biochemical basis of development, however beautiful, is inconsistent with the data and cannot serve as the explanation for the molecular basis of differentiation and development.

### Section 6: The consequences of CPM normalization and log transformation

As mentioned above, we found that, in the A20 & NIH3T3 cell lines, simple CPM normalization of the data generated two clearly different cell type groups without the need to perform any other transformations or feature selection. This raises the question of why such a transformation generates the two groups. Our analysis below follows closely the work of Townes et al[25], who, at least to our knowledge, first explored the consequence of these transformations on the underlying distributions of the data.

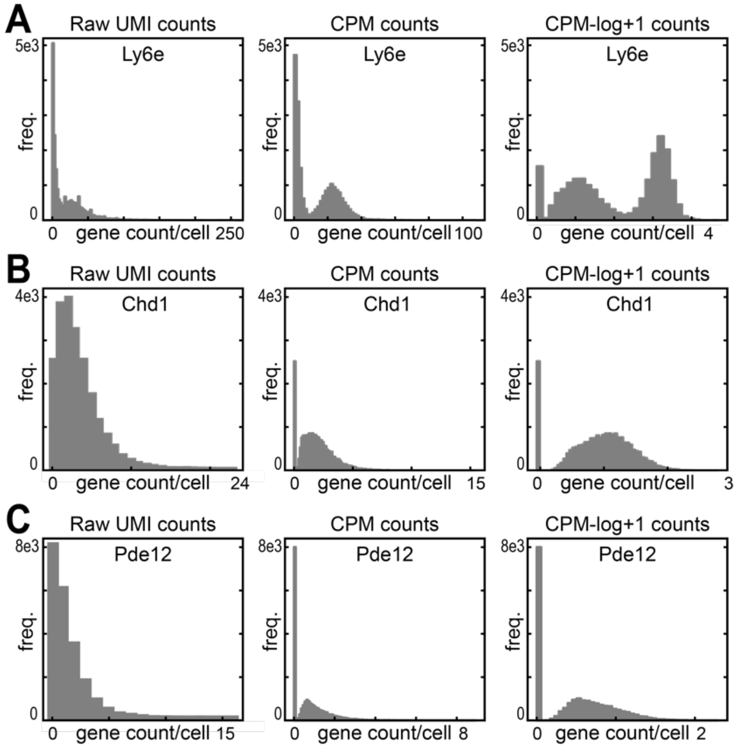

**Figure 5.** The effect of transformations on gene count distributions in scRNA-seq data **A,B)** Histograms illustrating the distribution of gene counts across cells from the multiplexed 10x A20 & NIH3T3 cell line data. In the left-most panel, raw UMI counts are plotted; in the middle panel, counts-per-million transformed counts are plotted; and in the right-most panel, counts-per-million then log+1 transformed counts are plotted for A) Ly6e, B) Chd1, and C) Pde12.

Consider two genes taken from the A20 & NIH3T3 data[26], presented both as raw UMI counts and as CPM normalized versions, with $Q = 10^4$ as is typically done in current analysis pipelines. The results are shown in Fig. 5. This transformation has a very clear effect on the data—it dramatically separates the 0's from the non-0 values. The reason for this is obvious: if $x_{i,j} = 0$ then $\hat{x}_{i,j} = 0$, so this transformation cannot alter the probability of finding a 0 value in the data (i.e. the height of the 0 bin in the histogram has to be the same before and after the transformation). The smallest non-0 value in the transformed space will generally be due to the cell(s) with the largest total number of counts that express just one copy of the gene, and for those cells $\hat{x}_{i,j} = Q/n_{i,T}$. As a consequence of all of this, a clearly unimodal distribution can be converted into a bimodal one; we might be able to model the untransformed distributions as Poisson or Negative Binomial distributions, but we would need a Zero Inflated version of such models for the transformed cases (Fig. 5). Note that the value of $Q$ just sets the distance between 0 and the minimum non-0 value in the dataset, so it controls the degree of separation observed (Fig. 5).

There are two ways in which CPM normalization could be helpful for generating separate cell type groups. For one, it could be that the variation in the raw counts $x_{i,j}$ does not conform to Waddington's landscape because those are not concentrations, but CPM normalization generates a dataset more similar to a concentration space, as discussed in section 5 above. In that scenario, we would expect that the variation in non-0 values of $\hat{x}_{i,j}$ to be important for describing cell type attractors. It could be, however, that the real effect of CPM normalization is to stretch the difference between 0 and non-0 values. The latter perspective is supported by the fact that, just visually in the data, there are *no clusters in the raw UMI counts*. But when we apply the transformation, the separation between 0 and non-0 values clearly generates two clusters.

To consider this possibility, we also performed our $\epsilon$ network analysis on the Hamming distance between a different transform of the gene expression space. Here, we set the gene expression value to “0” if the gene was not expressed, and “1” if the gene was expressed in that cell. This generates a scenario where there is no difference between cells in terms of their *expression levels*; we just care if the gene is expressed or not. Such a transformation is equivalent to taking all the cells in the non-0 group in the histograms in Fig. 5 and assigning them an expression value of 1. This preserves the idea that there are two broad groups (expressing and non-expressing) but disregards any differences in expression levels among expressing cells.

Note that this approach works just as well as the CPM normalization on the A20 & NIH3T3 data (Fig. S5.17). This suggests that, rather than working because it converts a space of counts to a space of concentrations, the CPM normalization generates two separate clusters because it emphasizes the distinction between 0 and non-0 values. Interestingly, in the one MERFISH dataset where we can actually calculate concentrations of mRNA based on known cell volumes, we see no separation whatsoever (Fig. S5.8F), suggesting that, in the few cases where CPM normalization on its own generates distinct groups in the $\epsilon$ network sense, it may do so because of this effect of separating expressing and non-expressing cells.

Interestingly, log transformation emphasizes this effect even further. If we take the histograms and log-transform them with a pseudocount of 1, we get the histograms shown in Fig. 5. These show an even greater distinction between 0 and non-0 values, again generating bimodal and “clustered” data from data that has no clusters at all. Note that changing both the value of the normalization constant $Q$ and the psuedocount $p$ can be used to modulate the distance between the 0 and non-0 values in the resulting distribution. In particular, reducing $p$ expands this gap significantly.

One might ask whether the above matters in the context of studying Waddington’s landscape. To begin, let us emphasize that CPM and log transformations, along with a host of other transformations and dimensionality reduction techniques we considered, *do not* work to separate most datasets into discrete cell-type groups (Figs. 5, S5.1-S5.8, S5.11-5.20). And we also do not generally see separation in the Hamming space for most datasets (Fig. S5.17). So the idea here is not that these CPM and log transformations *work* in most cases. When they do work, however, this seems to be due to the fact that the separation in the underlying cell groups is due to presence/absence patterns, rather than an attractor structure in *levels* of the macromolecules being measured. It may be that these transformations are popular because they can emphasize differences in such patterns that can be operationally useful for cell type clustering even though they don’t generate Waddington-like groups. Whether such on/off patterns are mechanistically relevant to the problem of cell fate determination is yet to be seen, but in any case, in most datasets even such on/off patterns are not sufficient to generate separation.

26. 40k Mixture of Mouse Cell Lines, Multiplexed Samples, 4 Probe Barcodes (Next GEM) 10x Genomics. https://www.10xgenomics.com/datasets/40k-mixture-of-mouse-cell-lines-multiplexed-samples-4-probe-barcodes-1-standard.
